# The retinoblastoma tumor suppressor limits ribosomal readthrough during oncogene induced senescence

**DOI:** 10.1101/788380

**Authors:** Neylen del Toro, Frédéric Lessard, Sarah Tardif, Jacob Bouchard, Véronique Bourdeau, Gerardo Ferbeyre, Léa Brakier-Gingras

## Abstract

The origin and evolution of cancer cells is considered to be mainly fueled by mutations affecting the DNA sequence. Although errors in translation could also expand the cellular proteome, their role in cancer biology remains poorly understood. Tumor suppressors called “caretakers” block cancer initiation and progression by preventing DNA mutations and/or stimulating DNA repair. If translational errors contribute to tumorigenesis, then caretakers genes will prevent such errors in normal cells in response to oncogenic stimuli. Here, we show that the retinoblastoma protein (RB) acts as caretaker tumor suppressor by preventing the readthrough of termination codons, a process that allows proteins to be synthetized with additional domains. In particular, we show that expression of oncogenic *ras* in normal human cells triggers a cellular senescence response characterized by a significant reduction of basal ribosomal readthrough. However, inactivation of the RB tumor suppressor pathway in these cells, using the viral oncoprotein E7 or the oncogenic kinase CDK4 increased readthrough. Conversely, activation of the RB pathway by the tumor suppressor PML, the ribosomal proteins RPS14/uS11 and RPL22/eL22 or the CDK4/6 inhibitor palbociclib reduced readthrough. We thus reveal a novel function for the RB pathway as a caretaker of translational errors with implications for tumor suppression and cancer treatment.

## Introduction

Cellular senescence is a tumor suppressor mechanism that prevents proliferation in cells bearing oncogenic stimuli (1). Senescent cells can efficiently halt tumor progression by remaining out of the cell cycle permanently as benign lesions (1, 2). Ideally, they also activate their elimination through immune mediated clearance (3, 4). However, if not cleared, some senescent cells can occasionally escape from their dormancy and progress into malignant tumors (5, 6). There is a great heterogeneity in the senescence response depending on the inducer, tissue type and genetic background (7). Nevertheless, most senescent cells activate the p53 and retinoblastoma (RB) tumor suppressor pathways that block cell cycle progression (1,2,8). In epithelial cells, reactive oxygen species and DNA damage were linked to senescence bypass and malignant progression (9, 10). These results also suggest that mutations that inactivate the major tumor suppressor pathways mediating senescence contribute to circumvent this process. Yet, in many human cell types, inactivation of either p53 or RB is not sufficient to bypass senescence (11, 12), indicating that senescence is a robust process and that disabling multiple genes is required to bypass it. Consistent with this idea, it has been recently demonstrated that epigenetic changes controlling the expression of several tumor suppressors contribute to senescence bypass and tumor progression (13–16). In addition to epigenetics, the cellular protein repertoire and thus their activities can be extended at the translational level (17). However, little is known about the contribution of translational errors in tumorigenesis.

In normal cells, translational errors in the form of aminoacid misincorporation are estimated to occur at a reduced frequency and unlikely to significantly affect the proteome (18). However, translational recoding by frameshifting or readthrough can generate novel protein variants with the potential to notably alter a cellular phenotype. In yeast, a prion protein called PSI+ impairs translation termination and confers advantage under stress conditions (19). Moreover, translational recoding is often used by viruses to increase the coding potential of their small genomes (17). For example, retroviruses use frameshifting to control the synthesis of viral replication proteins (20). Overall, translational readthrough, originally discovered in viruses and now extended to metazoans, can add a new C-terminal signal to proteins, changing their function and localization (21–23). The extensions usually contain potential signals to target proteins to the nucleus, peroxisomes, or the membrane (22, 23). Taken together, the available evidence suggests that tumor cells could use such recoding mechanisms to generate new protein variants and evolve. Therefore, tumor suppressor pathways could counteract readthrough as part of their mechanism of action.

Here we describe that translational readthrough is significantly reduced during oncogene-induced senescence, an anticancer response controlled by the tumor suppressors RB, p53 and PML. Using defined genetic tools, we also show that senescence-associated readthrough suppression is mediated by the RB tumor suppressor pathway.

## Materials and Methods

### Cell culture and materials

IMR-90 human diploid fibroblasts were obtained from Coriell Institute for Medical Research (New Jersey, NY) and ATCC. IMR-90 containing human telomerase reverse transcriptase (hTERT) were generated in our laboratory by transducing cells with the FG12-hTERT lentiviral vector. Phoenix ampho packaging cells used for retroviral infections were given by S.W. Lowe (Memorial Sloan Kettering Cancer Center, New York, NY). Cells were cultured in Dulbecco’s modified Eagle medium (DMEM; Wisent) supplemented with 10% fetal bovine serum (FBS; Wisent), 1% penicillin G/streptomycin sulfate (Wisent) and 2 mmol/L L-glutamine (Wisent). The translation elongation inhibitor cycloheximide and the aminoglycoside gentamicin sulfate were purchased from Sigma-Aldrich. The CDK4/6 inhibitor palbociclib (PD**-**0332991) was purchased from Chemietek. A phthalamide derivative named CDX5-1 was donated by M. Roberge (University of British Columbia, Vancouver, BC, Canada).

### Plasmids and cloning

Retroviral vectors pMSCV, pBabe/pBabe-ER, pBabe-H-RasV12, pWZL/pWZL-H-RasV12 were described in (24), pBabe-PML-IV-ER in (8), pLXSN, pLXSN-E6, pLXSN-E7, pLXSN-E6/E7 and pLXSN-E7 **Δ** 21-24 in (25), pBABE-RPL22(WT)-Myc in (26), pBABE-RPS14(WT)-Myc in (27). pBABE-CDK4(WT) was a gift from Scott W. Lowe (Memorial Sloan-Kettering Cancer Center, New-York, NY). The genes coding for Renilla luciferase (Rluc) and Firefly luciferase (Fluc) were linked by an intercistronic region in order to obtain a 96 kDa bifunctional protein (28). These reporters were PCR-amplified and subcloned in NotI/NsiI restriction sites to obtain pMSCV-Rluc-Fluc variants. A UGA stop codon, created by site-directed mutagenesis using PfuUltra II fusion HS DNA polymerase (Agilent, Canada), was inserted in the intercistronic sequence. Primers for PCR are provided in Supplementary table 1. The readthrough region from Moloney murine Leukemia Virus (MMuLV) as well as sequences flanking the stop codon from the readthrough sequence of Aquaporin 4 (AQP4, NM_001650.4) (29), were chemically synthesized (Biocorp, Canada) and subcloned in XhoI/ApaI restriction sites between Rluc and Fluc genes. The readthrough region sequences are provided in Supplementary table 2. The non-readthrough control was performed by mutating the stop codon TGA to CGA.

### Identification of novel readthrough events in mammalian cells using bioinformatic analysis

In order to look for endogenous readthrough candidates we first predicted in silico candidate readthrough events that could be detected by proteomics experiments. We built a database of peptide sequences located between the first arginine/lysine after the canonical stop codon and the next stop codon, always in the same reading frame using sequences deposited in RefSeq hg38.2. We discarded sequences shorter than 5 aminoacids. Candidate readthrough peptides were then matched to the data set of peptides identified by trypsin digestion of human proteins and mass spectrometry at (30). We found 278 readthrough specific peptides above the false discovery rate (FDR) threshold (Supplementary table 3). Following a readthrough propensity predictor algorithm (22), four candidates showing the highest readthrough propensity values were chosen. This algorithm assigns regression coefficients to the stop codon and all possible nucleotides in the stop codon context based on experimental data. The stop codon context comprises the nucleotide sequences from −6 to +9 positions surrounding the stop codon. The new readthrough candidates are: Vasodilator-stimulated phosphoprotein (VASP, NM_003370.3), Aspartate beta hydroxylase (ASPH, NM_004318.3), Hepsin (HPN, NM_182983.2) and Fibrillarin (FBL, NM_001436.3). Oligonucleotides containing 30 up-stream and 30 down-stream nucleotides flanking the stop codon, as well as the stop codon of those readthrough candidates were chemically synthesized (Biocorp, Canada) and subcloned in XhoI/ApaI restriction sites between Rluc and Fluc genes of our pMSCV-Rluc-Fluc reporter construct. The readthrough region sequences are provided in Supplementary table 2.

### Polysome fractionation

The protocol was performed as previously described (31). Briefly, 60% (w/v) sucrose stock solution was used to make sucrose gradients (5 to 50%) in a buffer containing 200 mM Tris-HCl (pH 7.6), 1 M KCl, 50 mM MgCl_2_, 100 µg/mL cycloheximide, 1X protease inhibitor cocktail (EDTA-free) (Roche) and 200 units/mL of RNase inhibitor (abm-Applied Biological Materials, BC). After 12 days post-infection, proliferating and senescent fibroblasts at 80-90% confluence were treated with cycloheximide at a final concentration of 100 µg/mL for 5 min at 37°C. Cells were washed with ice-cold 1X PBS - 100 µg/ml cycloheximide, scratched and lysed in a hypotonic buffer [5 mM Tris-HCl (pH 7.5), 2.5 mM MgCl_2_, 1.5 mM KCl, 1X protease inhibitor cocktail (EDTA-free), 100 μg/mL cycloheximide, 1 mM DTT, 100 units of RNAse inhibitor, followed by addition of 25 μl of 10% Triton X-100 (final concentration of 0.5%) and 25 μl of 10% sodium deoxycholate (final concentration of 0.5%). Cells were vortexed and then centrifuged at low speed (140 x *g*) for 15 min at 4°C. Sample supernatants were adjusted so that they contain the same OD (10-20 OD at 260 nm). The lysate was loaded onto the ultracentrifuge tubes containing sucrose gradients, then centrifuged at 22,223 x *g* (36,000 rpm) for two hr at 4°C using SW41Ti rotor Beckman Coulter (Optima L80 XP ultracentrifuge). After ultracentrifugation, the samples were placed in the UV detector (Brandel #IV-22-117140) and fraction collector (Retriever 500, Teledyne Isco). Each fraction was collected and monitored from the top to the bottom of the ultracentrifuge tube.

### Dual-luciferase Assays

IMR-90 transduced cells were washed with 1X PBS, scratched and lysed in 1X Passive lysis buffer supplied in the Dual Luciferase Stop & Glo® Reporter Assay System (Promega). A 20 μl cell lysate sample was used for luminescence measurements with Lumin Triathler-Hidex. After adding 50 μl of the Fluc reagent (Substrate + Buffer) to the samples, luminescence was measured for 10 seconds. Addition of 50 μl of the Rluc reagent-Fluc quenching (Stop & Glo®) preceded another 10 seconds measurement. In order to compare the readthrough fold-changes, relative Fluc activities (Fluc/Rluc) of each condition were normalized to the relative Fluc activities from non-senescent cells:

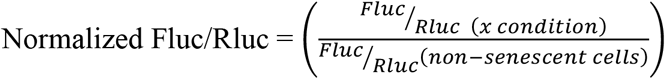

All assays were performed in technical triplicates. Normalized relative Fluc fold changes and Fluc/Rluc ratios are provided in Supplementary table 4.

### Immunoblotting

To prepare total cell extracts, cells were washed with 1X PBS containing 1X protease and phosphatase Inhibitor cocktails (Roche), scratched and collected by a centrifugation at low speed (140 x *g*) for 5 min, lysed in 200 μl of SDS sample buffer (62.5 mM Tris-HCl, pH 6.8, 10% glycerol, 2% SDS and 5% 2-mercaptoethanol), and boiled for 5 min. Fifteen μg of total cell proteins were separated on SDS-PAGE and transferred to nitrocellulose membranes (Millipore). The primary antibodies used were anti-Renilla Luciferase rabbit polyclonal (1:3000, Code No. PM047, MBL), anti-H-Ras mouse monoclonal (1:250, clone F235, Sc-29, Santa Cruz Biotechnology), anti-phospho-H3^S10^ rabbit polyclonal (1:1000, #06–570, Millipore, Billerica, MA), anti-RB mouse monoclonal (1:1000, clone 4H1, #9309, Cell Signaling), anti-MCM6 rabbit polyclonal (1:1000, A300-194A, Bethyl Laboratories), anti-p53 mouse monoclonal (1:1000, clone DO-1, sc-126, Santa Cruz Biotechnology), anti-c-Myc rabbit polyclonal (1:1000, clone A-14, sc-789, Santa Cruz Biotechnology) and anti-α-tubulin mouse monoclonal (B-5-1-2, 1:20000, Sigma). Signals were revealed after incubation with goat anti-mouse IgG (1:3000, #170-6516, Bio-Rad, Mississauga, ON) or goat anti-rabbit IgG (1:3000, #170-6515, Bio-Rad, Mississauga, ON) secondary antibodies, and by using enhanced chemiluminescence (ECL, Amersham) or Lumi-LightPLUS (Roche).

### Cell proliferation assay and Senescence-associated β-galactosidase assay

Relative density of cells was assessed from estimations of cell number according to crystal violet retention assay (8). The senescence associated β-galactosidase (SA-β-Gal) activity was measured at day 12 or 35 post-infection as previously described (8). All assays were performed in technical triplicates.

### RT-qPCR

Senescent and non-senescent IMR-90s were collected at 12-or 35-days post-infection, with either pBabe-H-RasV12 or pBabe control vector in TRIzol reagent (Invitrogen) and total RNA was extracted following manufacturer’s instructions. RNAs were reverse-transcribed into first-strand cDNA using All-in-One^TM^ First-Strand cDNA Synthesis Kit (abm-Applied Biological Materials, BC), diluted and then amplified by qPCR using LightCycler® 96 (Roche Diagnostics) and SYBR Green PCR mix previously described in (8). Analysis for indicated genes were done, using HMBS and TBP as reference genes. Primers for qPCR are provided in Supplementary table 5. All assays were performed in technical triplicates.

### Statistics and reproducibility

Statistical analysis (One-way ANOVA with post-hoc Tukey HSD (Honestly Significant Difference)) was done using the One-way ANOVA test calculator at: http://astatsa.com/OneWay_Anova_with_TukeyHSD/. Two-tailed Student’s *t-tests* were performed using GraphPad Prism version 6.0c software. A p<0.05 was considered statistically significant. Each experiment was repeated at least three times, except for those in Figures: 2C-D; 3B (UGA), and Supplementary figure S1C-E and S2D, which were done twice. SA-β-Gal assays were quantified from many fields within one experiment to represent the entire petri dish and confirmed as described in figures 5C and S1A.

## Results

### Translation termination is improved in oncogene-induced senescence

To investigate the effect of cellular senescence on readthrough efficiency, we first used a model of oncogene-induced senescence (OIS) in human primary cells. Primary fibroblasts were retrovirally transduced with pBabe-H-RasV12 (senescence inducer) and the control empty vector pBabe. They were also transduced with luciferase reporter plasmids containing either: 1) a TGA within an artificial intercistronic region (UGA readthrough); 2) the stop codon context from AQP4, a well-known mammalian readthrough candidate (29); or 3) the non-readthrough controls (TGA mutated to CGA) between Rluc and Fluc fusion protein gene (Figure 1A). Immunoblotting shows a band at 35 kDa corresponding to Renilla luciferase, as well as a weak band at 100 kDa indicating the fusion protein Renilla-Firefly luciferase produced after a ribosomal readthrough in cells transduced with the AQP4 readthrough reporter. For UGA reporter, the fusion protein is not detectable by the western blot technique. On the other hand, in cells transduced with the non-readthrough control only the fusion protein can be detected (Figure 1B). At early stages of OIS, cells present a cancer-like behavior characterized by a hyperproliferation phase. However, around 6 days after introduction of oncogenic *ras*, cells stop proliferation and enter a well-characterized and stable cell cycle arrest (27). We confirmed this senescent cell cycle arrest at day 12 post-infection, using an assay for the senescence-associated β-galactosidase activity that allowed visualizing characteristic large and flat cells that stained positive for this biomarker (Figure S1A). RT-qPCRs were carried out to evaluate the decrease of Ki67 expression, indicating the proliferation arrest, and the activation of p53 and RB pathways in senescent cells (Figure S1B). Global translation was not affected in senescent cells since their polysome profiles (Figure 1C) were similar to control growing cells. This is consistent with previous reports and with the fact that senescent cells are actively secreting multiple pro-inflammatory mediators (32). In addition, the non-readthrough (CGA) controls presented similar relative Fluc activities both in senescent and proliferating cells (Figure S1C). Cap-dependent Rluc luciferase expression increased with time (Figure 1D). Readthrough-dependent translation of the firefly luciferase reporter increased early after introduction of oncogenic *ras* in normal human fibroblasts when cells proliferate rapidly but it decreased when the cells underwent OIS as measured at days 12 or 20 after H-Ras-introduction (Figure 1E). The efficiency of readthrough was calculated by dividing Fluc by Rluc values, then normalized to the relative Fluc activities from non-senescent cells (Vector). The results clearly show that the senescent program reduced the readthrough efficiency at later stages (Figure 1F). In contrast, control cells did not change readthrough efficiency during the same time in culture (Figure S1D). These results suggest that oncogenic stimulation decreases the efficiency of translation termination but cellular senescence counteracts this potentially oncogenic mechanism. To discard the possibility that readthrough efficiency might be decreased due to a cell-cycle arrest independent from the senescent phenotype, proliferating cells were starved for a week to induce quiescence. Starved normal cells compared to cells supplied with fetal bovine serum (FBS) showed no differences in the efficiency of readthrough. Besides, starved and FBS-supplied senescent cells showed similar readthrough values (Figure S1E). This observation leads us to conclude that the decrease of readthrough efficiency is senescence-specific. Immunobloting shows the gradual reduction of phosphorylated RB as well as the absence of MCM6, and the reduction of the mitotic marker H3-pS10 in H-Ras cells, confirming the establishment of senescence. P53 accumulates in senescent cells but decreases after 20 days post-infection in H-Ras cells (Figure 1G).

**Figure 1.**
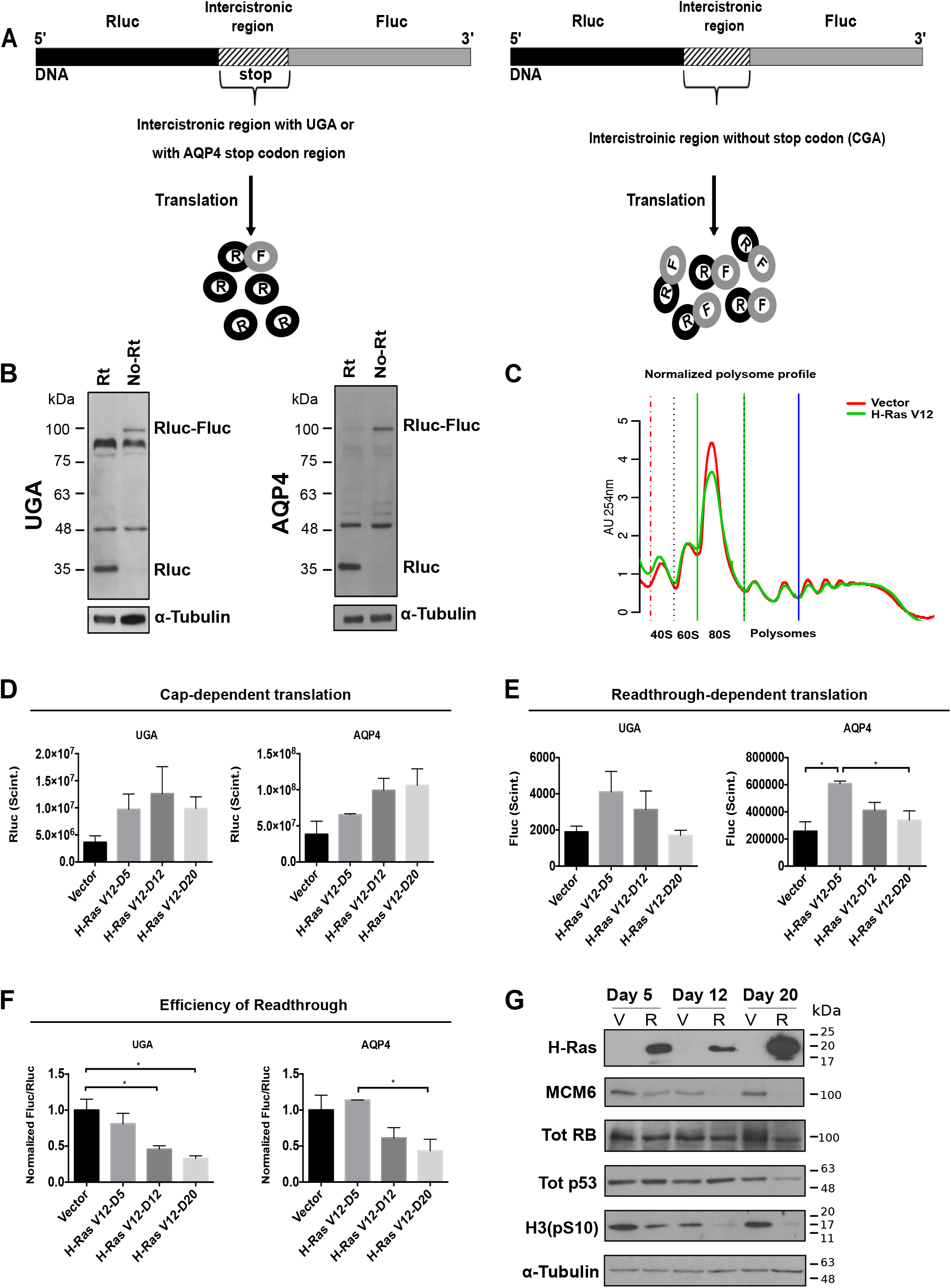
Readthrough is reduced in OIS. **A.** Rluc (Renilla luciferase) in black is linked to Fluc (Firefly luciferase) in gray, by an intercistronic streaked region. Rluc is the internal control of the gene, while Fluc is the readthrough sensor. A UGA codon within an artificial context or the AQP4 stop codon region are inserted in the intercistronic region (left). The expression of Fluc indicates the efficiency of readthrough. The intercistronic region from the non-readthrough control lacks the stop codon (TGA mutated by CGA) (right). **B.** Immunoblots for Rluc in IMR-90 cells transduced with UGA and AQP4 reporters (Rt) or with the non-readthrough controls (No Rt), 12 days post-infection. Tubulin was used as a loading control. **C.** Polysome profiles were performed 12 days post-infection with an empty vector (Vector) or H-Ras V12 oncogene (H-RasV12) showing that global translation is similar in senescent and non-senescent IMR-90 cells. **D.** Rluc plots indicate that luciferase cap-dependent translation is not affected in IMR90 infected cells with an empty vector (Vector) or in senescent cells induced with the oncogene H-Ras V12 (H-RasV12) at day 5 (D5), 12 (D12) or 20 (D20). **E.** Fluc plots indicate the decrease of readthrough dependent-translation following induction of senescence with the oncogene H-Ras V12 (H-RasV12) at day 5 (D5), 12 (D12) or 20 (D20). **F.** Normalized Fluc/Rluc ratios indicate that readthrough efficiency decreases progressively in senescence. IMR-90 cells were transduced with an empty vector (Vector) or H-RasV12 oncogene (H-RasV12) to establish senescence and with UGA/AQP4 dual luciferase reporter plasmids. Luciferase activities were measured at days 5, 12 and 20 post-infection. Normalizations are presented as means relative to empty vector-infected cells from three independent experiments with technical triplicates for each experiment. One-way ANOVA with post-hoc Tukey HSD (Honestly Significant Difference) tests were performed. Error bars indicate SD of biological triplicates. Tukey HSD p-values indicate that * = p<0.05, is significantly different. **G.** H-Ras, total RB (Tot RB), Total p53 (Tot p53), MCM6, H3(pS10) and tubulin at day 5, 12 and 20 post-infection from non-senescent (V) and senescent cells (R). Blots in **G** are representative of 3 independent experiments with similar results.

The aminoglycoside gentamicin disrupts prokaryotic protein synthesis by binding to 16S ribosomal RNA, but also affects eukaryotic ribosome proofreading, inducing a conformational change in the ribosome-mRNA complex stimulating the efficiency of readthrough (33). We sought to investigate whether OIS can decrease readthrough stimulated by gentamicin. We added 900 μg/ml of gentamicin sulfate to OIS cells 24 hours before luciferase assays and found that the readthrough efficiency was still reduced in senescent cells both at UGA stop codons and the natural readthrough sequence of AQP4 (Figure 2A). Strikingly, readthrough levels were the same in senescent and proliferating cells with the reporter containing the programmed readthrough region from Moloney Murine Leukemia Virus (MMuLV) (Figure 2A). This region contains a UAG stop codon followed by a pseudo-knot structure (34) and it allows up to 10% of ribosomes to suppress translation termination in order to synthesize the Gag-Pol polyprotein in cells infected with the virus (35). Also, the aminoglycoside gentamicin did not affect MMuLV readthrough efficiency. The non-readthrough controls were unaffected by gentamicin stimulation (Figure S2A). We then treated cells 72-hours before cells lysis with 600 μg/ml of gentamicin and 25 μM of CDX5-1, a novel small molecule that potentiates readthrough efficiency when combined with aminoglycosides although by itself has no effect (36). We observed that while single gentamicin treatment increased UGA readthrough efficiency up to 6.6-fold in control cells, the combination increased it up to 16.5-fold. Surprisingly, the readthrough efficiency was still limited in senescent cells after the combination, which improved the efficiency only 1.7-fold over gentamicin treatment alone (Figure S2B).

To investigate more broadly whether senescence affects endogenous readthrough signals, we made luciferase reporters using the stop codon context of several mRNAs, which we predicted *in silico* as candidates for readthrough (see Material and Methods). We tested both their basal and gentamicin-stimulated readthrough efficiency. We found that the readthrough of VASP and HPN were significantly reduced during OIS while those of FBL showed a tendency to be reduced (Figure S2C). ASPH readthrough was undetectable neither in proliferating nor in senescent cells. Nevertheless, after gentamicin stimulation, we could observe the readthrough decrease in senescent cells (Figure S2D).

**Figure 2.**
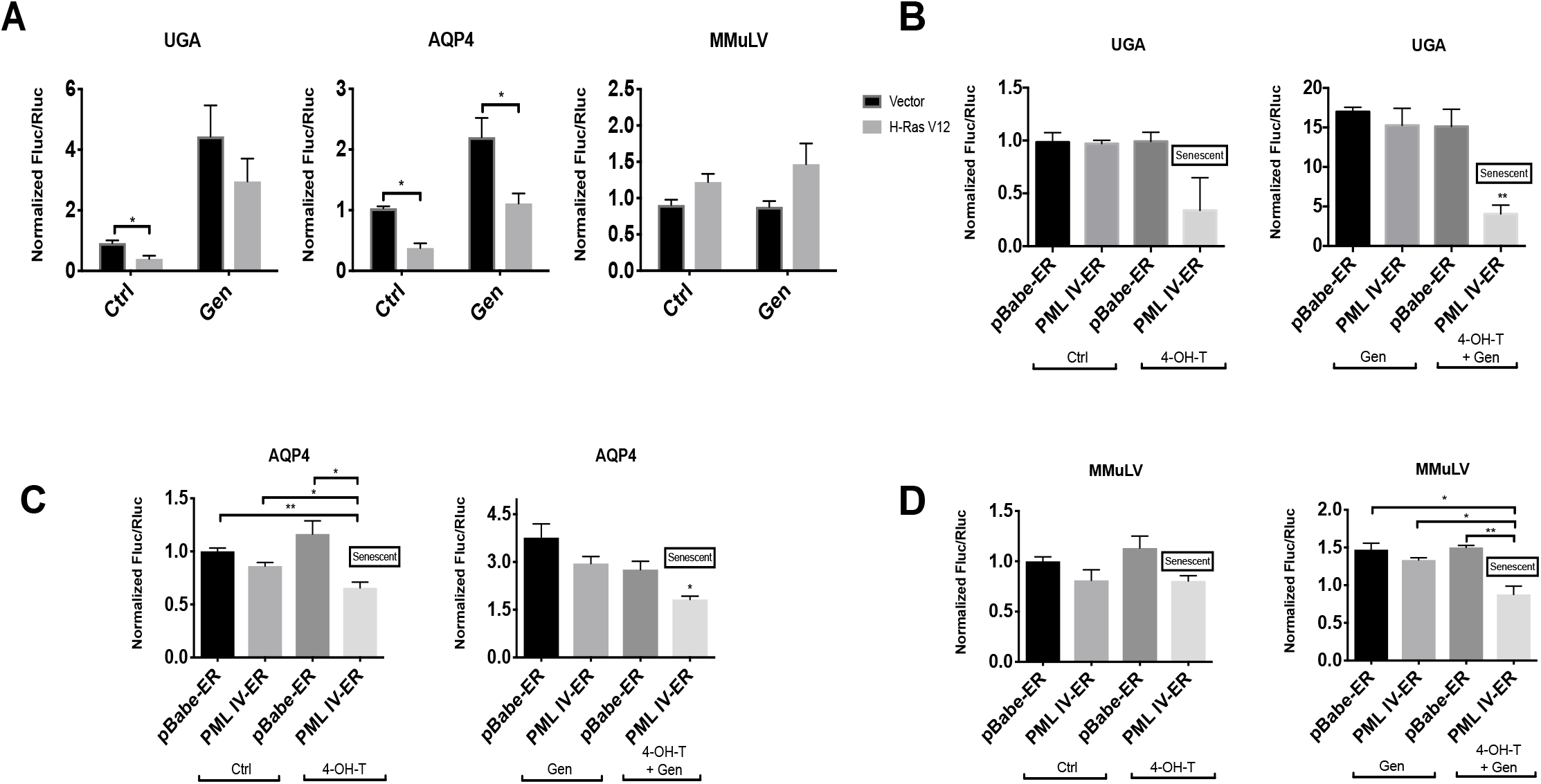
Gentamicin - dependent translation errors are reduced in OIS and PML - induced senescence. **A.** IMR-90 cells were transduced with an empty vector (Vector) or H-RasV12 oncogene (H-RasV12) to induce OIS and with luciferase reporters. A UGA within an artificial context, a stop codon within the natural context from AQP4 or from MMuLV gag-pol protein were inserted in the intercistronic region of the reporter. Cells were treated with vehicle (Ctrl) or 900 μg/ml of gentamicin sulfate (Gen) for 24 hours before measuring luciferase activities at day 12 post-infection. Unpaired Student’s *t*-test with equal SD were performed. Error bars indicate SD of biological triplicates. * = p<0.05 is significantly different, using two-tailed Student’s *t*-test. **B-D.** IMR-90 cells were transduced with pBabe-ER empty vector (pBabe-ER) or pBabe-PML-IV-ER (PML IV-ER), and with luciferase reporters UGA, AQP4 or MMuLV. Cells were treated with vehicle (Ctrl) or 100 nM 4-OH-Tamoxifen (4-OHT), inducing PML-IV nucleus translocation to induce senescence. Moreover, cells were treated with vehicle or 900 μg/ml of gentamicin sulfate (Gen) for 24 hours before measuring luciferase activities at day 12 post PML-IV induction. Normalized Fluc/Rluc ratios indicate the efficiency of readthrough. Normalizations are presented as means relative to empty vector-infected cells from three independent experiments with technical triplicates for each experiment, except for **C, D** which are representative of 2 independent experiments with similar results. **B-D**: One-way ANOVA with post-hoc Tukey HSD were performed. Error bars indicate SD of biological triplicates (**B**) or technical triplicates (**C, D**). Tukey HSD p-values indicate that * = p<0.05, ** = p<0.01 are significantly different.

### Translation termination is improved in PML-induced senescence

The tumor suppressor PML controls p53 and RB which are central for the establishment of OIS response (8). Also, PML plays a role in antiviral responses (37, 38) and, since many viruses use readthrough to monitor gene expression, we wanted to investigate whether PML modulates this process. Normal human fibroblasts IMR-90 cells were retrovirally transduced with a vector that allows a conditionally inactive PML (pBabe-PML-IV-Estrogen Receptor) to be expressed as well as with UGA, MMuLV and AQP4 luciferase reporters, in order to analyze the readthrough efficiency in this model. Transduced cells were stimulated with 100 nM of the estrogen antagonist 4-OH-Tamoxifen, to induce PML-IV nuclear translocation and senescence (13). We found that induction of PML dramatically reduced basal or gentamicin-induced readthrough at UGA stop codons (Figure 2B). Intriguingly, the effect of PML on gentamicin-induced readthrough was much more important than the effect of OIS, suggesting that PML could be one important regulator of readthrough efficiency. Readthrough levels were also reduced after 4-OH-Tamoxifen stimulation, with or without gentamicin, in AQP4 and MMuLV transduced cells (Figure 2 C-D). To our surprise, the decrease of readthrough efficiency in MMuLV reporter leads us to speculate about an antiviral response role of PML-IV.

### Circumventing senescence decreases the fidelity of translation termination

Although senescence in response to oncogenes includes a very stable cell cycle arrest, some cells escape from the process and progress towards malignant transformation. Cells that circumvent OIS display a gene expression profile typical of malignant cells and chromosomal aberrations (39). We sought to determine whether these cells loss the tight control over readthrough. We obtained cell populations that bypassed OIS from long-term (35 days) cultures of IMR90 cells expressing oncogenic *ras*. Cells that escaped senescence revert their morphology to that of normal growing cells (not shown) and express lower levels of oncogenic *ras* (Figure 3A), indicating a mechanism that allow them to prevent pro-senescence *ras*/ERK signaling. In fact, we showed previously that decreasing ERK signaling accelerates senescence bypass in H-Ras-expressing cells (39, 40). Cells that bypassed senescence also had lower levels of the CDK inhibitor p16INK4a and higher levels of the proliferation marker KI67 (Figure 3A). We next measured readthrough efficiency in these cells using the UGA stop codon or the natural signals for AQP4 (29), VASP and HPN. The last two signals were found using bioinformatic analysis as described in Materials and Methods. In all cases, the cells that bypassed senescence displayed an increased readthrough level even higher than control non-senescent cells (Figure 3B). Actually, the cells that bypassed senescence had readthrough ratios similar to gentamicin-treated cells (excluding UGA reporter).

**Figure 3.**
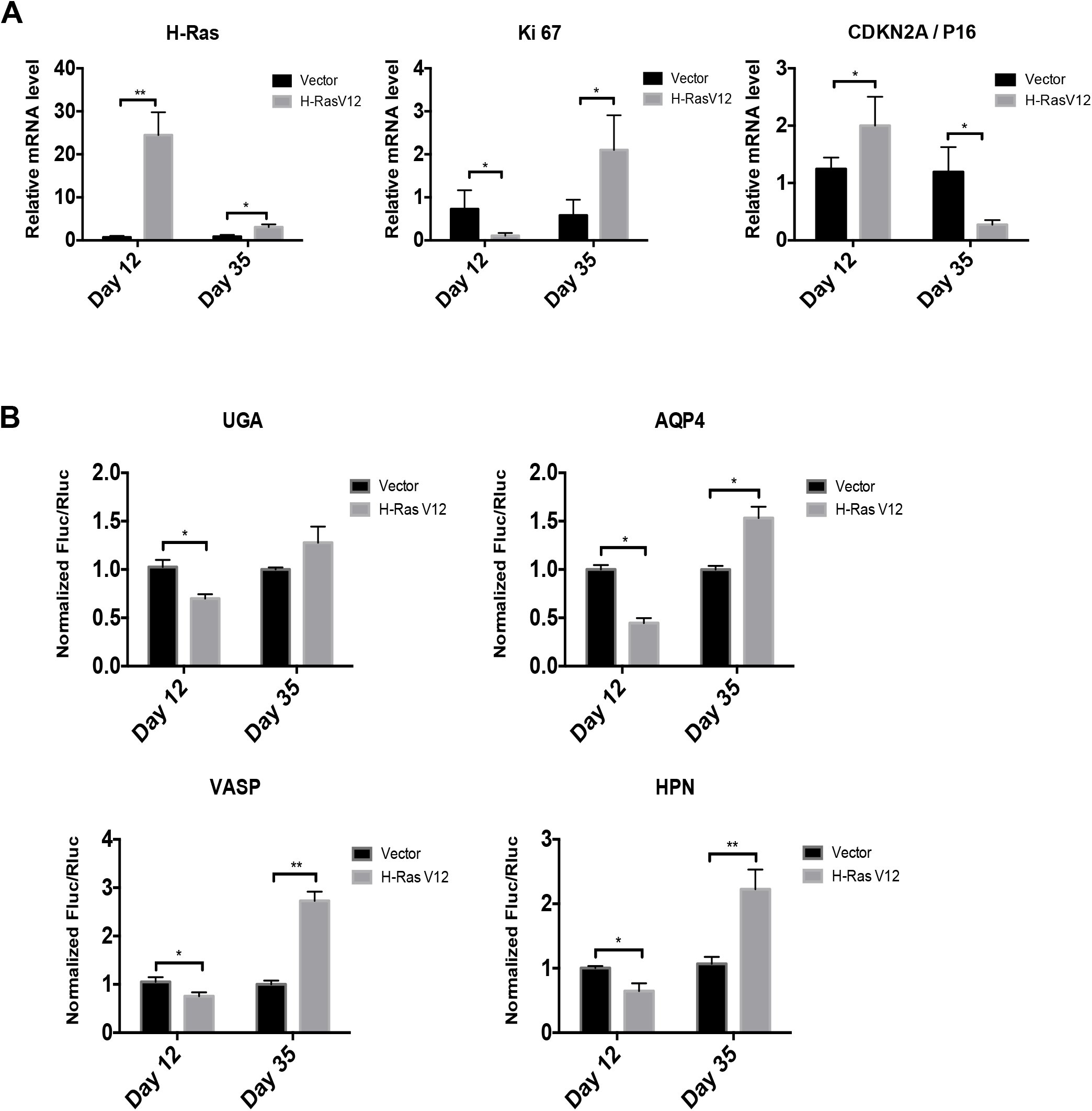
Bypassed-OIS shows increased readthrough. **A.** RT-qPCR for senescent marker mRNAs were performed in IMR-90 cells at day 12 and 35 post-infection with an empty vector (Vector) or H-RasV12 oncogene (H-RasV12). Data are normalized over TBP and HMBS, and presented as means relative to vector infected cells. Error bars indicate SD of biological triplicates. * = p<0.05, using two-tailed Student’s *t*-test. **B.** Luciferase activities in non-senescent and senescent cells measured at day 12 and 35 post-infection in cells having the indicated reporters. Normalizations are presented as means relative to vector-infected cells from three independent experiments with technical triplicates for each experiment, except for UGA reporter, which are representative of 2 independent experiments with similar results. Error bars indicate SD of biological triplicates (AQP4, HPN, VASP) or technical triplicates (UGA). * = p<0.05, ** = p<0.01 are significantly different, using two-tailed Student’s *t*-test.

### E7 Oncoprotein increases readthrough

Since low translation fidelity and increased readthrough are associated to malignant transformation (41) and the escape from senescence, we next investigated which tumor suppressor pathways activated in senescent cells modulate readthrough. To investigate whether p53 and/or RB pathway is implicated in translation termination efficiency, we used E6 (inhibits p53) and E7 (inhibits RB) oncoproteins from HPV-16 virus. Transduction of E6 did not affect the readthrough levels in senescent cells. However, senescent cells infected with E7 oncoprotein, as well as with both E6 and E7, showed a significant increase in readthrough efficiency at UGA stop codons and the natural readthrough signals from AQP4 and VASP (Figure 4A-C). These results indicate that E7 targets, likely the pocket proteins family including RB, p107 and p130, mediate the readthrough decrease in senescent cells. We also knocked-down the p53 and RB mRNAs’ expression with shRNAs in H-Ras cells and we failed to see the increase of readthrough levels (data not shown) in cells transduced with shp53. This reproduced the E6–H-Ras transduced cells result. The readthrough efficiency in H-Ras cells with shRB knock-down was increased, although not significantly, suggesting that RB in coordination with p107 and p130 must influence the reduction in readthrough efficiency.

**Figure 4.**
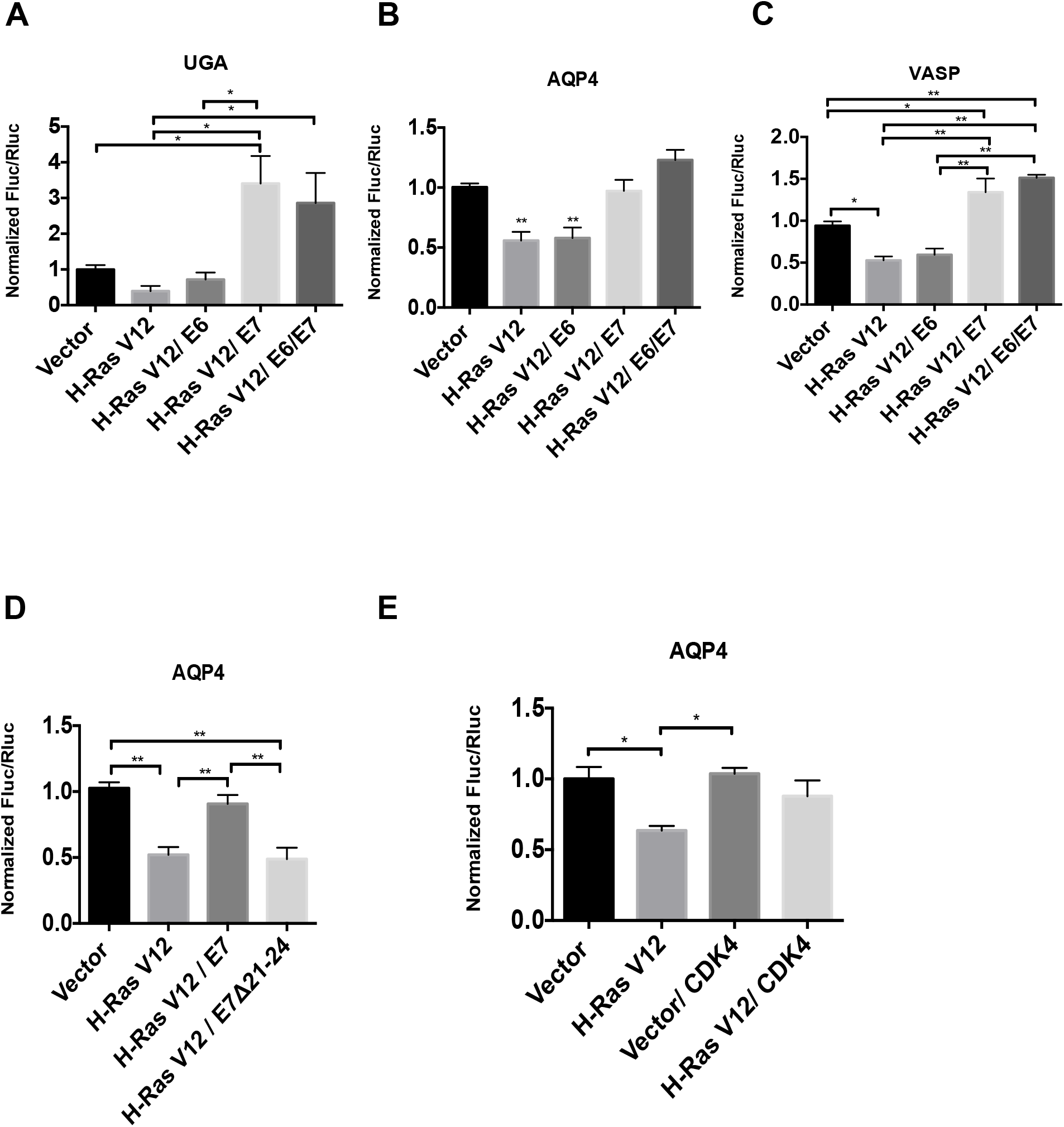
RB pathway disruption affects the efficiency of readthrough. Normalized Fluc/Rluc ratios to indicate the efficiency of readthrough. A UGA within an artificial context, a stop codon within natural contexts from AQP4 or VASP were inserted in the intercistronic region of Rluc-Fluc luciferase reporter. **A-C.** IMR-90 cells containing HTERT were transduced with an empty vector (Vector) or H-RasV12, an empty control vector (pLXSN), E6, E7 or E6/E7 oncogenes and luciferase reporters to study the readthrough efficiency in proliferating *vs* senescent *vs* transformed-like cells. **D.** IMR-90 cells were transduced with an empty vector (Vector) or H-RasV12, an empty control vector (pLXSN), wild-type E7 or E7 Δ21-24 mutant oncogene. **E.** IMR-90 cells were transduced with an empty vector or H-RasV12 or CDK4 and luciferase reporters to study the readthrough efficiency variations in proliferating *vs* senescent *vs*. cells overexpressing CDK4. **A-E.** Luciferase activities were measured in non-senescent and senescent cells at day 12 post-infection. Normalizations are presented as means relative to vector-infected cells from three independent experiments with technical triplicates for each experiment. One-way ANOVA with post-hoc Tukey HSD were performed. Error bars indicate SD of biological triplicates. Tukey HSD p-values indicate that * = p<0.05, ** = p<0.01 are significantly different.

To further implicate the RB pathway in the control of readthrough in senescent cells, we used an in-frame deletion mutant of E7 whose interactions with RB, P107 and p130 are disrupted (E7 **Δ**21-24) (42). As expected, H-Ras cells transduced with wild type E7 showed a two-fold increase in readthrough levels compared to senescent cells but the mutant E7 **Δ**21-24 failed to do so (Figure 4D). Besides, the proliferating cells transduced with E7 presented a tendency to increase readthrough, while the readthrough levels were not affected in proliferating cells transduced with the mutant E7 **Δ**21-24 (Figure S3A). RT-qPCRs were carried out to evaluate the expression increase of E2Fs targets in E7 transduced cells, as well as their decrease in E7 **Δ**21-24 mutant (Figure S3B). Next, we used a retroviral vector for CDK4 overexpression to block pocket proteins activity in senescent cells (43) and, as expected, we observed an increase in readthrough efficiency (Figure 4E). Relative cell growth and RT-qPCRs show that CDK4 overexpression does not affect proliferation arrest behavior at day 12 post-infection in senescent IMR-90s (Figure S3C-D). Taken together, these results indicate that the observed reduction in readthrough is not a simple consequence of the growth arrest of senescent cells and strongly suggest that the activation of the RB tumor suppressor pathway increases the fidelity of translation termination during OIS.

### The CDK4/6 inhibitor palbociclib contributes to readthrough reduction

Palbociclib blocks the cyclin-dependent kinase (CDK) 4/6 activity by competing with ATP binding and, as a result, RB remains in its hypo-phosphorylated active form (44). Treatment of proliferating fibroblasts with palbociclib reduced readthrough of the AQP4 reporter and gentamicin-stimulated readthrough at the UGA stop codon (Figure 5A-B). Interestingly, palbociclib reduces readthrough in cells that spontaneously escaped from senescence (Figure 5C). As expected, palbociblib also inhibits cell proliferation and strongly activates RB pathway in proliferating and H-Ras-escaped IMR-90s (Figure 5D, S4). Besides 16% of the proliferating population was positive to the blue-stained β-gal assay and the H-Ras-bypassed cells rapidly restored their senescent phenotype (50 % of cell population positive to β-gal staining) (Figure 5E).

**Figure 5.**
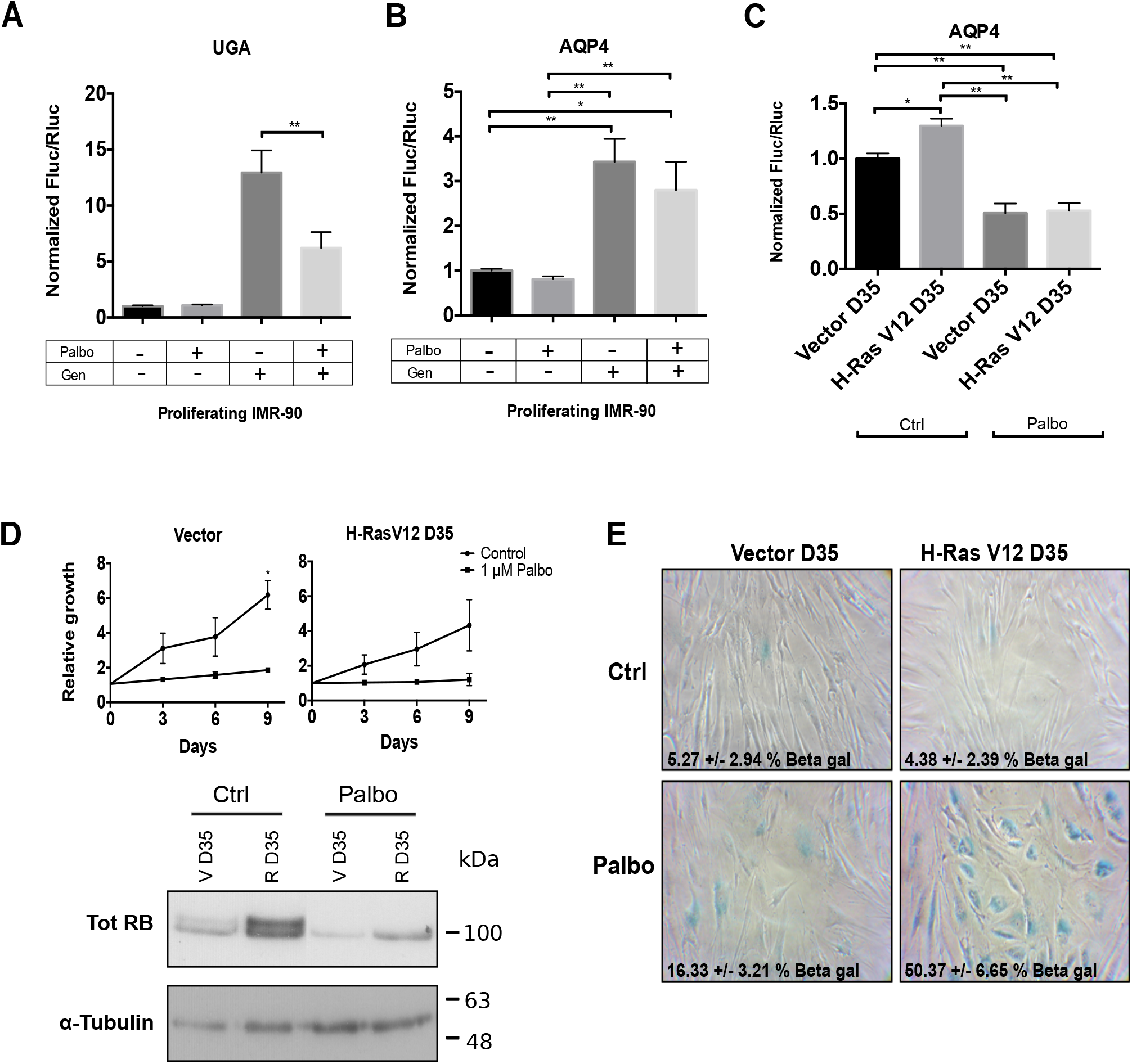
Palbociclib, a CDK4/6 inhibitor, contributes to readthrough reduction. **A.** Normalized Fluc/Rluc ratios of IMR-90 cells transduced with luciferase reporters (UGA, AQP4) and treated with vehicle and/or 1 μM of palbociclib (Palbo) and/or 900 μg/ml of gentamicin sulfate (Gen) for 5 days before measuring. **C.** Normalized Fluc/Rluc ratios of IMR-90 fibroblasts transduced with an empty vector (Vector) or H-RasV12 oncogene (H-RasV12) to induce OIS and with AQP4 luciferase reporter. Cells were treated with vehicle (Ctrl) or 1 μM of palbociclib (Palbo) for 5 days before measuring luciferase activities at day 35 post-infection. Normalizations are presented as means relative to vector-infected cells from three independent experiments with technical triplicates for each experiment. One-way ANOVA with post-hoc Tukey HSD. Error bars indicate SD of biological triplicates. Tukey HSD p-values indicate that * = p<0.05, ** = p<0.01 are significantly different. **D.** (Top) Growth curves of proliferating and *ras* by-passed IMR-90 cells treated with vehicle (Control) or 1 μM of palbociclib (Palbo) for 5 days are shown. Data are presented as means normalized to day 0 of each condition and error bars indicate SD, n = 3 independent experiments. ** = p<0.01 is significantly different, using two-tailed Student’s *t*-test. (Bottom) Immunoblots for indicated proteins: total RB (Tot RB) and tubulin from non-senescent (V D35) and *ras* by-passed cells (R D35) following treatments with vehicle (Ctrl) or 1 μM of palbociclib (Palbo) for 5 days. Blots are representative of 3 independent experiments with similar results. **E.** SA-β-gal of proliferating (Vector D35) IMR90 cells and IMR90s that have by-passed the senescent stage (H-RasV12 D35) treated for 5 days with vehicle (Ctrl) or 1 μM of palbociclib (Palbo) and fixed at day 35 (D35) post-infection. Data were quantified from many fields within one experiment to represent the entire petri dish. Three independent cell counts up to a total of at least 100 cells are presented as the mean percentage of positive cells.

### Ribosomal proteins RPS14/uS11 and RPL22/eL22 limit readthrough

We recently demonstrated that extra-ribosomal functions of RPS14/uS11 and RPL22/eL22 were linked to the regulation of the cell cycle and senescence. We found that these two proteins interact with the CDK4-Cyclin D1 complex inhibiting its activity and consequently activating the RB pathway (26, 27). These results suggest that RPS14/uS11 and RPL22/eL22 could, as palbociclib, affect readthrough via the RB pathway. We transduced IMR-90 cells with Myc tagged RPL22/eL22 or RPS14/uS11 as well as with the AQP4 readthrough reporter and measured readthrough efficiency at day 7, 12 and 14 after transduction. The expression of RPL22/eL22 or RPS14/uS11 was confirmed by immunoblots for the Myc tag. We also measured biomarkers of senescence such as RB, the E2F target MCM6 and the mitosis marker phospho H3 (Figure 6A). Cells bearing RPS14/uS11 showed a 15% decrease of readthrough at days 7 and 12 after transduction, and 30% decrease of readthrough 14 days after transduction compared to control cells (Figure 6B). In contrast, RPL22/eL22-transduced cells showed a peak of readthrough reduction at day 7 post-infection, and a progressive recovery 12-and 14-days post-infection (Figure 6C). We already reported that RPS14/uS11 is more efficient than RPL22/eL22 for inhibition of CDK4 and induction of senescence (26), explaining this difference. These results suggest that RPS14/uS11 haploinsufficiency in the 5q-syndrome (45) and frequent RPL22/eL22 hemizygous gene deletions found in cancer (46) could play a role in proteome diversity and tumorigenesis.

**Figure 6.**
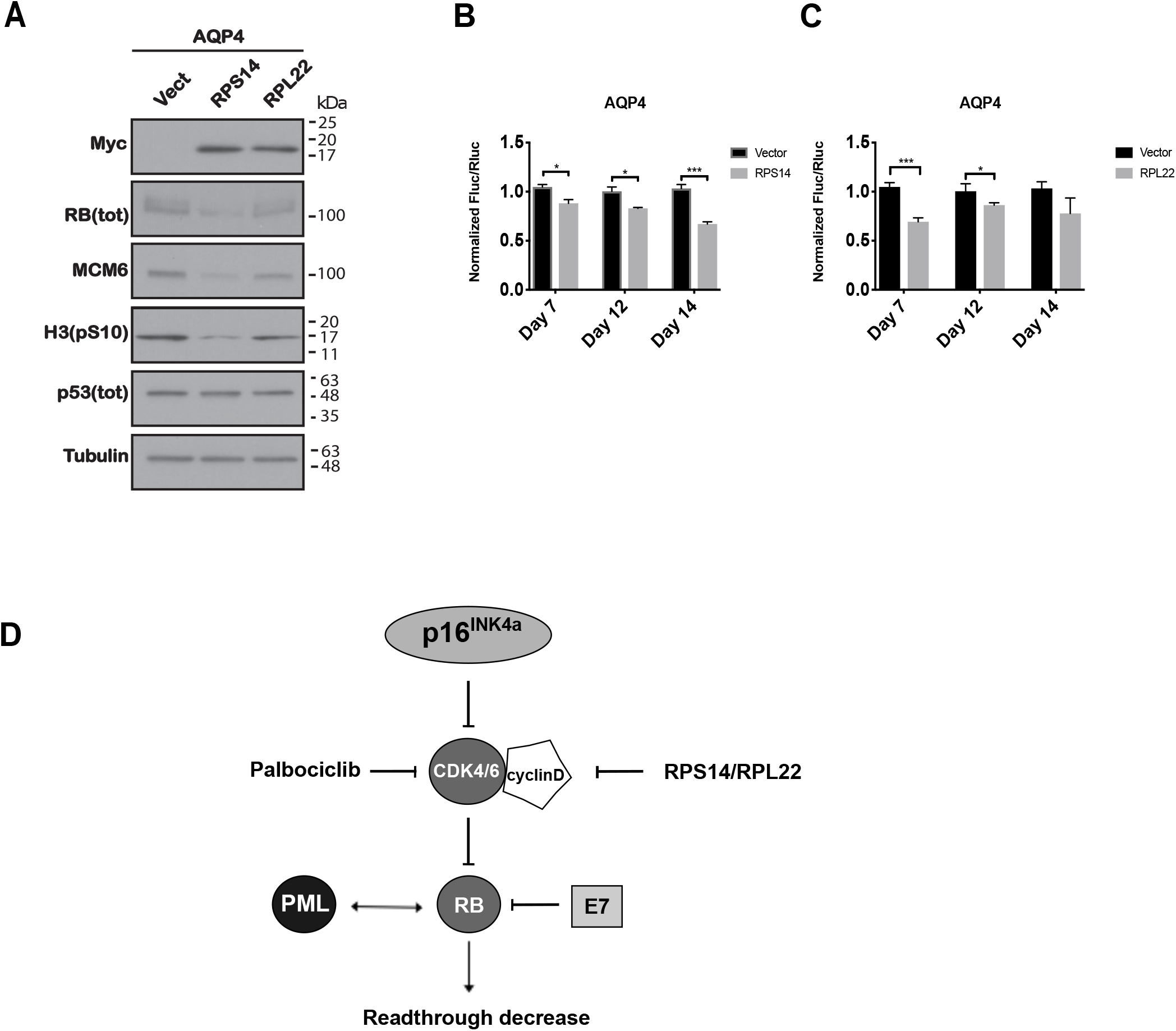
Ribosomal proteins RPS14/uS11 and RPL22/eL22 limit readthrough. **A.** Immunoblots for indicated proteins at day 7 post-infection with an empty vector (Vect), or pBABE-RPS14(WT)-Myc, or pBABE-RPL22(WT)-Myc: Myc (Myc-tag), total RB (RB tot), MCM6, H3(pS10), total p53(p53 tot). Tubulin was used as a loading control. Blots are representative result from three independent experiments with similar results. **B-C.** Normalized Fluc/Rluc ratios of IMR-90 cells transduced with an empty vector (Vector), or **(B)** pBABE-RPS14(WT)-Myc, or **(C)** pBABE-RPL22(WT)-Myc, and with luciferase reporter AQP4. Luciferase activities were measured in Vector and RPs-transduced cells at day 7, 12 and 14 post-infection. Normalizations are presented as means relative to vector-infected cells from three independent experiments with technical triplicates for each experiment Unpaired t tests with equal SD were performed. Error bars indicate SD of biological triplicates. * = p<0.05, ** = p<0.01, *** = p<0.001 are significantly different, using two-tailed Student’s *t*-test. **D.** Schema showing the RB activation/inhibition factors that modulate readthrough.

In conclusion, our results suggest that the RB pathway is implicated in translation termination accuracy, since RB degradation mediated by E7, or CDK4 overexpression, increases readthrough efficiency. On the contrary, RB activation after palbociclib treatment, PML-IV inducible expression as well as RPS14/uS11 and RPL22/eL22 overexpression improves translation termination (Figure 6D).

## Discussion

We show here that the RB tumor suppressor pathway reduces translational readthrough at UGA stop codons from dual luciferase reporters in the context of cellular senescence, a tumor suppressor response. In addition, the RB pathway also reduces readthrough in reporters containing the readthrough context of mRNAs coding for AQP4, VASP, FBL, ASPH and HPN. We propose a novel caretaker tumor suppressor activity for the RB pathway preventing the generation of C-terminally extended proteins by readthrough a process that can potentially confer new localization signals or interaction partners to many proteins (22, 47). The potential for readthrough to confer advantages to cells in stress was previously reported in yeast cells expressing the prion PSI that enhances both readthrough and resistance to several stresses (19). Our results suggest that cellular senescence and, in particular, the RB pathway prevents the oncogenic potential of proteome reprogramming via translational readthrough.

Senescence reduces translational readthrough in dual luciferase reporters by a factor of two, or four (for AQP4 reporter). This is around the same level of regulation reported by miRNAs. miRNAs regulate multiple genes to affect a particular phenotype (48) and by analogy, we propose that the effects of readthrough depend also on multiple genes. Studies with ribosome profiling in *Drosophila* and human fibroblasts reveal that more than 300 mRNAs undergo translational readthrough (49), suggesting that many new functions could be generated by this mechanism.

Senescence also reduced the stimulatory effect of aminoglycosides and the enhancer effect of CDX5-1 on translational readthrough (33). It has been reported that CDX5-1 potentiates G418-induced readthrough up to 180-fold compared to the aminoglycoside G418 alone (36). Here we showed a clear resistance of senescent cells to induce C-terminal extended proteins. Green and Goff reported that aminoglycosides can increase readthrough efficiency of the MMuLV gag-pol junction in HEK-293 cells (50). However, using normal fibroblasts, we found that the same drugs did not have such effect. HEK-293 cells have an inactivation of the RB and PML tumor suppressor pathways due to expression of adenovirus E1A oncoprotein. In our normal cells, disruption of these pathways increases readthrough. It is thus plausible that in the context of an altered control in HEK293 cells, the MMuLV gag-pol junction readthrough is stimulated by aminoglycosides. In addition, Green and Goff used several aminoglycosides in their studies, gentamicin being the less active and its effects were reported as dose-independent and not consistent. Since gentamicin does increase readthrough in other stop codon contexts, we suggest that the pseudoknot structure of the viral signal may interfere with the binding or the action of gentamicin on the ribosome.

Our results indicate that translation termination tends to be inaccurate in cancer-like cells where the RB pathway is dysfunctional but is particularly efficient in senescent cells. Early work by Stanners and colleagues showed that SV40-mediated transformation increased mistranslation in mammalian cells (51). SV40 encodes for the large T antigen that binds and inactivates the RB family of tumor suppressors (52) like the E7 oncoprotein we used in this study. Diaz and colleagues found that translational fidelity is decreased in the most aggressive breast cancer cell lines (53). They also reported that the p53 tumor suppressor pathway controls translational fidelity and in particular the translation of oncogenic proteins from IRES (41). In our experimental system, the RB pathway was more important to control readthrough levels in OIS. However, we did notice that inactivation of p53 in cells where RB is also inactivated further increased readthrough levels, suggesting also an important role for p53 in controlling readthrough levels (Figure 4A-C). The translation initiation factor eIF3 interacts with post-initiation ribosomes and controls readthrough (54, 55). The RB tumor suppressor pathway may repress the transcription of eIF3 subunits or other factors that modulate readthrough. An additional interesting mechanism relevant to senescence is the sequestration of eIF3 in PML bodies, which also contain RB (8). Consistent with the latter mechanism the eIF3 subunits EIF3K and EIF3E (Int-6) localize to PML bodies (56, 57). Viral proteins such as HTLV-I Tax can delocalize EIF3E from PML bodies (58), an event that could facilitate readthrough.

Of special interest for a plausible pharmacological modulation of readthrough levels, we found that the CDK4 inhibitor palbociclib was able to reduce readthrough and re-induce the stable cell cycle arrest in cells that bypassed senescence. If the phenotypic plasticity conferred by C-terminal extended proteins plays a causal role in the origin of human cancers, palbociclib could be used as a chemopreventive agent in patients with premalignant lesions susceptible to progress into malignant tumors.

## Acknowledgements

This study is supported by a NSERC grant to L B-G. N.D.T. acknowledges a studentship support from the GRUM (Université de Montréal), F.L. acknowledges support from the FRSQ and CRS (Cancer Research Society), S.T. acknowledges a summer NSERC studentship, G.F. is the recipient of a CIBC chair for breast cancer research at the CR-CHUM. We are grateful to I. Topisirovic, K. Tandook, and M. Leibovitch at the Lady Davis Institute, Department of Oncology, McGill University, for their help and advices in polysome fractionation assays, M. Roberg at the University of British Columbia for CDX5-1 and X. Roucoux, J-F. Lucier and F-M. Boisvert at the Université de Sherbrooke, Département de Biochimie, for their help in the identification of readthrough events by bioinformatic analysis.

---

**Supplementary Figure 1.**
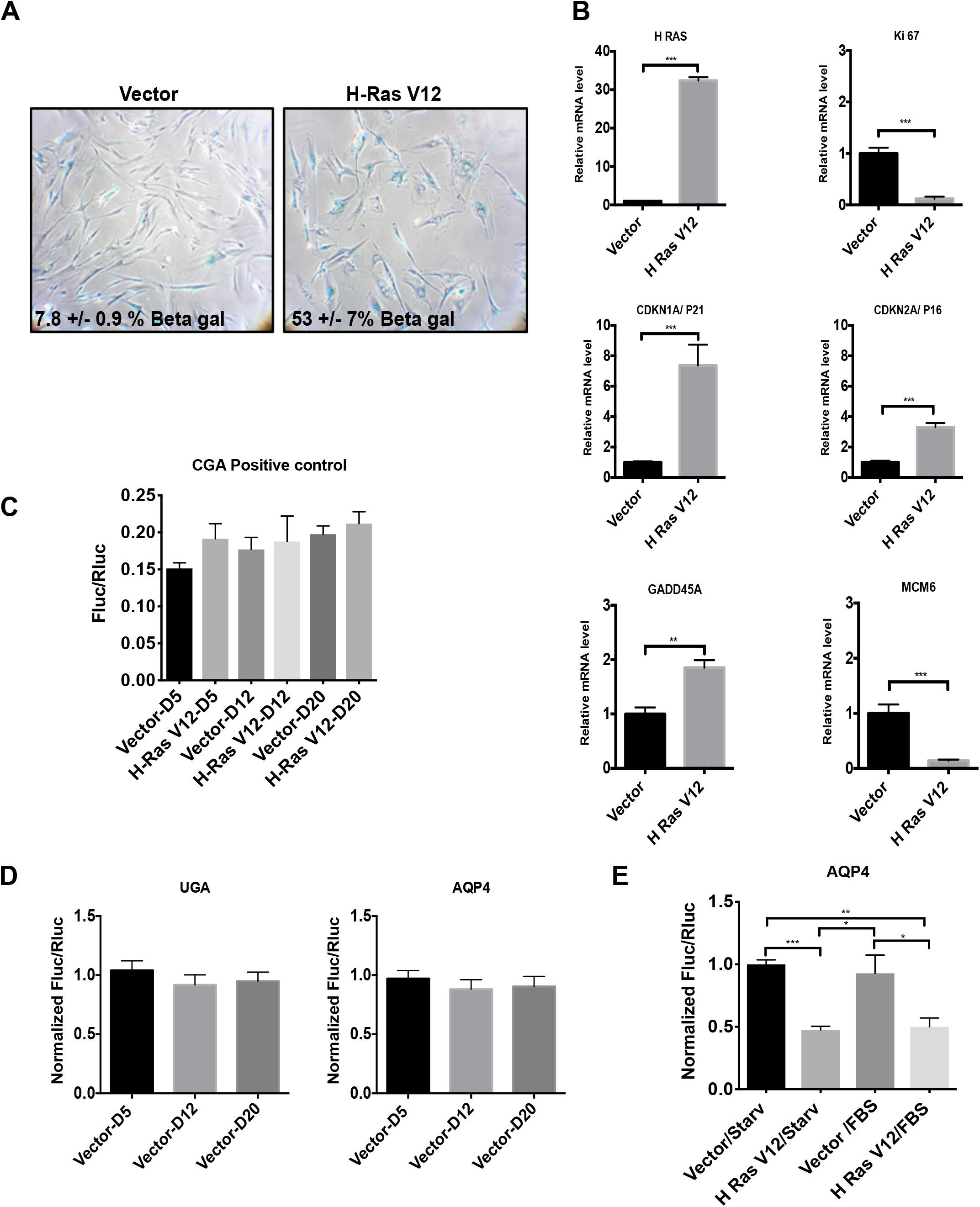
Readthrough is reduced in OIS. **A.** SA-β-gal of proliferating (Vector) and senescent IMR90 (H-RasV12 oncogene) cells, fixed at day 12 post-infection. Data were quantified from many fields within one experiment to represent the entire petri dish. Three independent cell counts up to a total of at least 100 cells are presented as the mean percentage of positive cells. **B.** RT-qPCRs for senescent marker mRNAs were performed in IMR-90 cells at day 12 post-infection. Data are normalized over TBP and HMBS, and presented as means relative to vector infected cells. Error bars indicate SD of technical triplicates. ** = p<0.01, *** = p<0.001 are significantly different, using two-tailed Student’s *t*-test. **C.** Luciferase activities from non-readthrough control (CGA-Positive control) in proliferating (Vector) and H-RasV12 senescent cells (H-RasV12) were measured at days 5 (D5), 12 (D12) and 20 (D20) post-infection. Assays are representative of 2 independent experiments with similar results with technical triplicates for each experiment. One-way ANOVA with post-hoc Tukey HSD were performed. Error bars indicate SD of technical triplicates. **D.** Normalized Fluc/Rluc ratios indicate that readthrough level does not change in non-senescent cells (Vector) at day 5 (D5), 12 (D12) or 20 (D20) post-infection. Error bars indicate SD of biological triplicates. **E.** Luciferase activities from AQP4 reporter were measured at day 12 post-infection. Normalized Fluc/Rluc ratios from non-senescent (Vector) and senescent cells (H-RasV12), starved (Starv) or supplied with fetal bovine serum (FBS) are represented. Starvation was performed for a week to induce quiescence. Data are presented as means relative to empty vector-infected cells, which are representative of 2 independent experiments with similar results with technical triplicates for each experiment. One-way ANOVA with post-hoc Tukey HSD were performed. Error bars indicate SD of technical triplicates. Tukey HSD p-values indicate that ** = p<0.01 is significantly different.

**Supplementary Figure 2.**
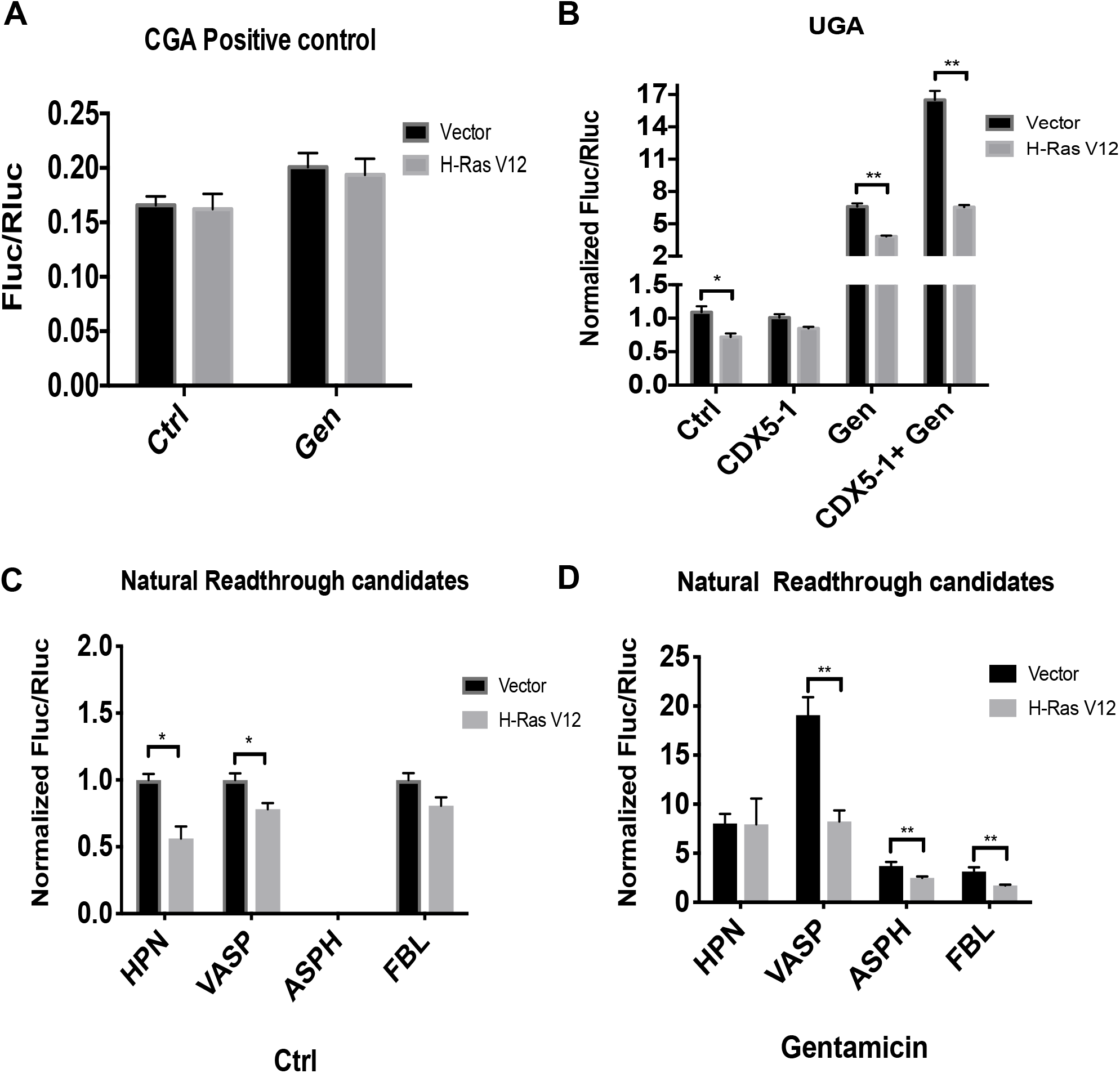
Senescent cells are resistant to gentamicin-induced readthrough. **A.** IMR-90 cells were transduced with an empty vector (Vector) or H-RasV12 oncogene to induce OIS and with indicated non-readthrough control luciferase reporter. Cells were treated with vehicle (Ctrl) or with 900 μg/ml of gentamicin sulfate (Gen) 24 hours before measuring luciferase activities at day 12 post-infection. Error bars indicate SD of biological triplicates. * = p<0.05, ** = p<0.01 are significantly different, using two-tailed Student’s *t*-test. **B.** IMR-90 cells were transduced with an empty vector (Vector) or H-RasV12 oncogene to induce OIS and with a UGA luciferase reporter. Cells were treated with vehicle (Ctrl) and/or 600 μg/ml of gentamicin sulfate (Gen) and/or 25 μM of CDX5-1 72 hours before measuring luciferase activities at day 12 post-infection. Error bars indicate SD of biological triplicates. * = p<0.05, ** = p<0.01 are significantly different, using two-tailed Student’s *t*-test. **C.** A stop codon within natural contexts from VASP, ASPH, HPN and FBL, was inserted in the intercistronic region of Rluc-Fluc luciferase reporter. IMR-90 cells were transduced with an empty vector (Vector) or H-RasV12 oncogene to induce OIS and with luciferase reporters. Luciferase activities were measured in non-senescent and senescent cells at day 12 post-infection. Error bars indicate SD of biological triplicates. * = p<0.05, ** = p<0.01 are significantly different, using two-tailed Student’s *t*-test. **D.** Cells, as in (**C**), were treated with 900 μg/ml of gentamicin sulfate (Gen) 24 hours before measuring luciferase activities at day 12 post-infection. Normalized Fluc/Rluc ratios indicate the efficiency of readthrough. Normalizations are presented as means relative to vector-infected cells from three independent experiments with technical triplicates for each experiment. Unpaired Student’s *t*-test with equal SD were performed. Error bars indicate SD of biological triplicates.

**Supplementary Figure 3.**
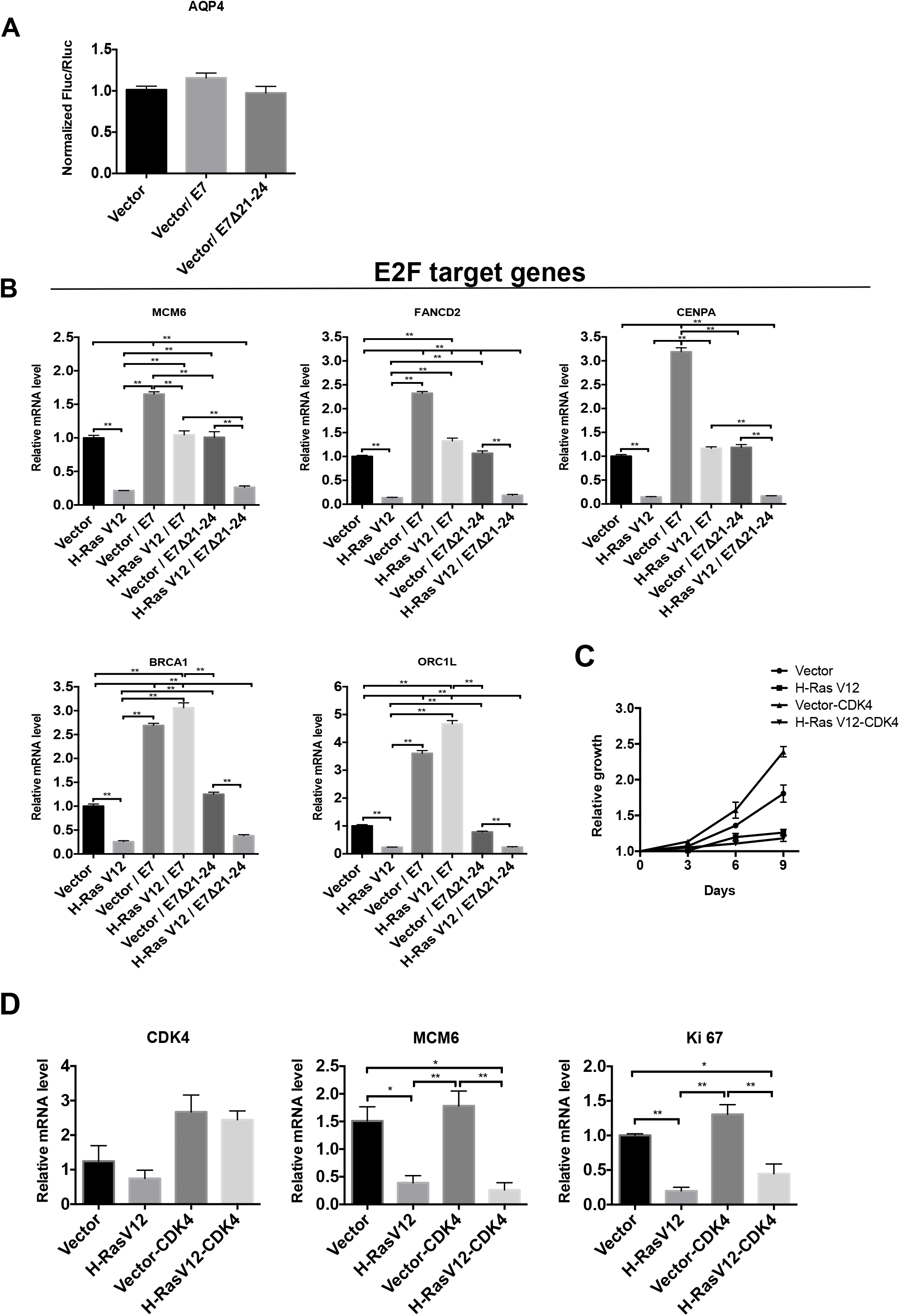
RB pathway disruption affects the efficiency of readthrough. **A.** IMR-90 cells were transduced with luciferase reporter AQP4 and an empty vector (Vector), wild-type E7 or E7 Δ21-24 mutant oncogene. Luciferase activities were measured at day 12 post-infection. Normalized Fluc/Rluc ratios indicate the efficiency of readthrough. Normalizations are presented as means relative to vector-infected cells from three independent experiments with technical triplicates for each experiment. Error bars indicate SD of biological triplicates. **B.** Representative RT-qPCR from E2Fs’ target mRNAs were performed in IMR-90 cells transduced with an empty control vector (pLXSN), wild-type E7 or E7 Δ21-24 mutant oncogene, and with an empty vector (Vector) or with H-RasV12 oncogene to induce OIS at day 12 post-infection. Data are normalized over TBP and HMBS, and presented as means relative to vector infected cells. Experiments were done three times with technical triplicates for each experiment. Error bars indicate SD of technical triplicates. Tukey HSD p-values indicate that ** = p<0.01 is significantly different. **C.** Growth curves of non-senescent (Vector) and H-RasV12 senescent IMR-90s overexpressing or not CDK4. Data are presented as means normalized to day 0 of each condition. Assays are representative of 2 independent experiments with similar results with technical triplicates for each experiment. Unpaired Student’s *t*-test with equal SD were performed. Error bars indicate SD of technical triplicates. **D.** RT-qPCRs for CDK4, MCM6 and KI67 mRNAs were performed in IMR-90 cells as in **(C)** and at day 12 post-infection. Data are normalized over TBP and HMBS, and presented as means relative to vector infected cells from three independent experiments with technical triplicates for each experiment. One-way ANOVA with post-hoc Tukey HSD were performed in **A, B, D** and Unpaired Student’s *t*-test with equal SD was performed in **C**. Error bars indicate SD of biological triplicates. Tukey HSD p-values indicate that * = p<0.05, ** = p<0.01 are significantly different.

**Supplementary Figure 4.**
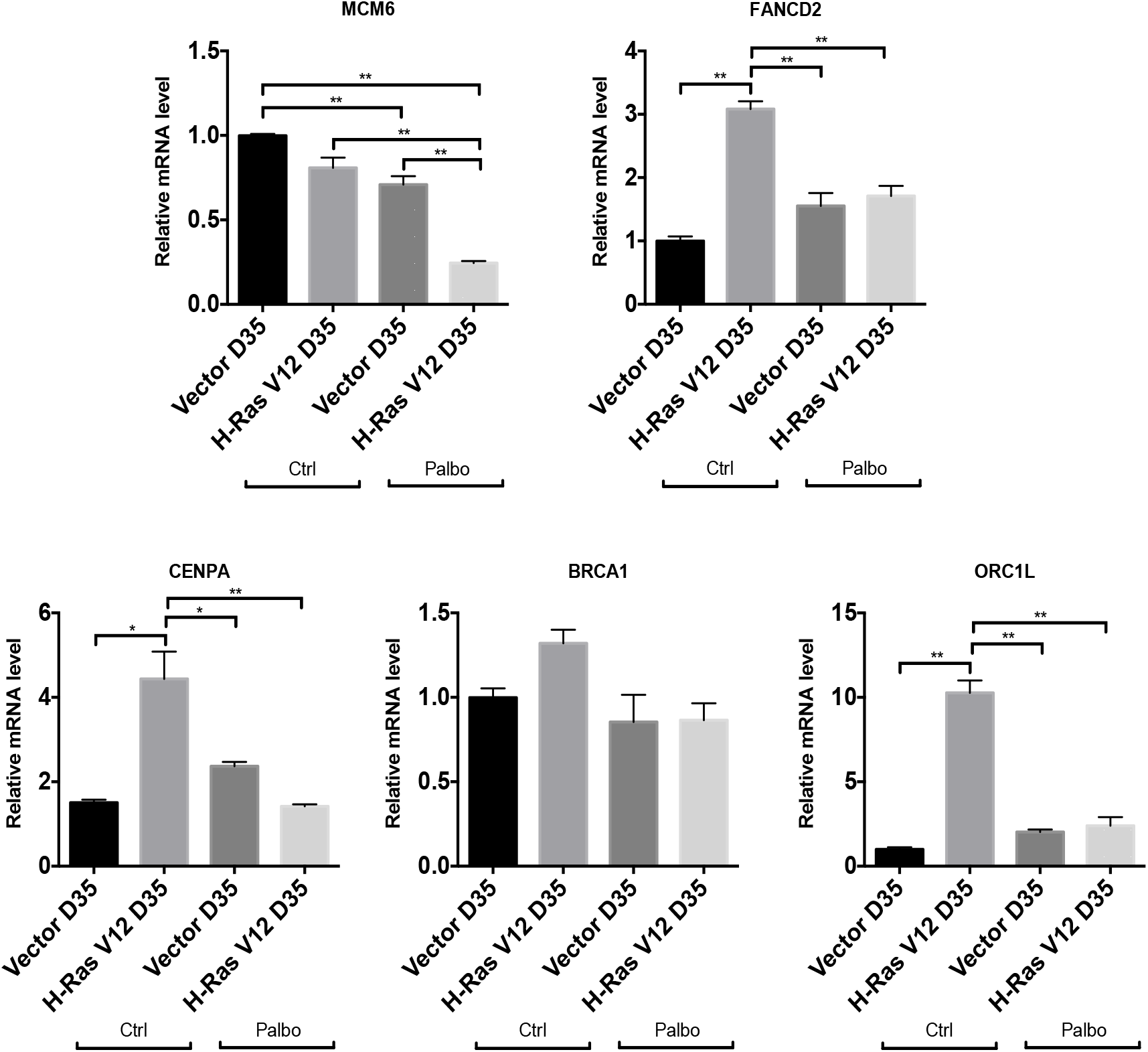
E2Fs target genes. RT-qPCR from E2Fs’ target mRNAs were performed in IMR-90 at day 35 post-infection with an empty vector (Vector D35) or with the oncogene H-RasV12 (H-RasV12 D35) and treated with vehicle (Ctrl) or with 1 μM of palbociclib (Palbo) for 5 days before cell lysis. Data are normalized over TBP and HMBS, and presented as means relative to vector infected cells from three independent experiments with technical triplicates for each experiment. One-way ANOVA with post-hoc Tukey HSD were performed. Error bars indicate SD of technical triplicates. Tukey HSD p-values indicate that * = p<0.05, ** = p<0.01 are significantly different.

