## Supplemental tables for "The retinoblastoma tumor suppressor limits ribosomal readthrough during oncogene induced senescence"

**Supplementary table 1. PCR cloning primers**

| Name | 5' forwad primer (5'-3') | 3' reverse primer (5'-3') |
|---|---|---|
| pMSCV-RLUC-FLUC | GGAAGAAAGCGGCCGCTTCAGATCCGCTAGAGCC | TGGATGCATGAATTTCGTCATCGCTGAATAC |
| UGA STOP CODON | CCAAGCTTGCACCAGTGACCAGCAAGGTGTACG | CGTACACCTTGCTGGTCACTGGTGCAAGCTTGG |

**Supplementary table 2. Readthrough region**

| Gene name | | | Readthrough context |
|---|---|---|---|
| AQP4 | Aquaporin 4 | NM_001650.4 | ATCTGGAGAGGTATTGTCTTCAGTATGACTAGAAGATCGCACTGAAAGCAGACAAGACTCCTTAGAA |
| ASPH | Aspartate beta hydroxylase | NM_004318.3 | AGAACAGCAGGAAGTACCACCAGATACTTAAAGCTTCAAAAAGACTGCCCCTACCACCGGGCC |
| FBL | Fibrillarin | NM_001436.3 | GTACAGGCCACCCCCCAAGGTGAAGAACTGAAGTTCAGCGCTGTCAGGATTGCGAGAGATGGG |
| HPN | Hepsin | NM_182983.2 | TCGAGGCCAGCGGCATGGTGACCCAGCTCTGACCGGTGGCTTCTCGCTGCGCAGCCTCCAGGG |
| MMuLV | Moloney Murine Leukemia Virus | | CTAGGGAGGTCAGGGTCAGGAGCCCCCCCCTGAACCCAGGATAACCCTCAAAGTCGGGGGGCC |
| VASP | Vasodilator-stimulated phosphoprotein | NM_003370.3 | GGAGCTGAGGAAGCGGGGTTCTCCCTGACCACAGGGACCCAGAAGACCCGCTTCTGGGCCCCA |

### SUPPLEMENTARY TABLE 3. Readthrough candidates

| Readthrough code | Gene name | peptides found by MS analysis | | | peptides « confi | stop- DNA 3'UTR | RTP (readthough propensity)(Ref 22) | RTP Linfs3 (Ref 22) |
|---|---|---|---|---|---|---|---|---|
| | | | | | | TGAcaaacacagcccctgaggggccccgggagtggccttggctccctggagagcccacgtctcagccacagttctccact | | |
| RT_012331.1 | FAM19A5 family with sequence similarity 19 (chemokine (C-C motif) | APSEDGLRLR | NP_001076436. | | 3608 | cgcctcggacttcacccgttctctgccgcccgcccactccgtttccctgtggtccgtgaaggacggcctcaggccttggcatcctgagcttcggtctgtccagccgacccgaggaggccggactcagacacataggcgggggcggcacctggcatcagcaatacgcagtctgtgggagcccggccgcgcc | 0.309 | 0.646 |
| RT_006854.1 | ZNF182 zinc finger protein 182 | YGSKFSLSSFHLSHNPK | NP_001171570.1, NP_001 | | 3732 | TAGagagctaatcagcagtatctattatggacactgaaagaaagttgtgtcaattttactcacattttaaaagtatattttgacttgtggtttaaatatgggagtaaattcagcctttcttcttttcatctcagtcataacccaaaagctacgggagaaaaagcagaagtgtaa | 0.143 | 0.215 |
| | | | | | | TGAgcccgcggcggactgccccaggactgcgggaaggcgcgggagccaggggcctccaccccaaccccggctcagagactgggcggcccggcgggggctccctggggaccccagatgggctcagtgctgccacccagatcagatcacatgggaatgaggc | | |
| RT_020801.1 | ACADS acyl-CoA dehydrogenase, C-2 to C-3 short chain | LGGPAGAPWGPQMGSVLPPR | NP_000008.1, N | | 8117 | cctccgaccattggcagctccgcctctgggcctttccgcctcctcaccactgtgcctcaagttcctcatctaagtggccctggcctcctgggggcggggttgtggggggctgagcgacactcagggacacctcagttgtcctcccgcgggccctggtgccctggcatga | -0.077 | 0.066 |
| RT_017649.1 | ACSL5 acyl-CoA synthetase long-chain family member 5 | VLKYLPAHCALLVR | NP_976313.1, NP_976314 | | 690 | TAGgataaggtacttaagtacctgccggcccactgtgcactgcttgtgagaaaatggattaaaaactattcttacatttgttttgcctttcctcctattttttttaa | -0.08 | -0.226 |
| | | | | | | TGAtggggcagtggcctctgtgggcccttgtcaagagctggaggccactcccaagagcctctcctatggcagctgggacg | | |
| RT_009441.1 | ADCK2 aarF domain containing kinase 2 | AGGHSQEPLLWQLGR | NP_443085.2 | XP_01151 | 464 | ttttaaaattgggacaccaatttcaaatgtaaccctccagtggtggaaggcacaccatggcttcctctgcttggtttga | 0.237 | 0.065 |
| | | | | | | TGAgggctcctgctggattccgaaaaggccgggaagccagtctccttccctgaagcaagcccaggagaagactctccgcc | | |
| | | | | | | ccacgccaatcccaaaggcatgcagatggctgtgcatgttggcttctttggacctgcactggaggatttctcagacacat | | |
| RT_013024.1 | ADCY7 adenylate cyclase 7 | GLLLDSEK | NP_001105.1 | XP_01152 | 9511 | gcaccagattctggctcgaagcagccactgagccataa | -0.029 | 0.028 |
| | | | | | | TGAaggcgcatggttctggacggcccagcagctcctgtggccacagcagctttgtacacgaagaccatccatcctcccctt | | |
| RT_020165.1 | ADGRE5 adhesion G protein-coupled receptor E5 | MVLDGPAAPVATAALYTK | NP_510966.1, N | | 6737 | cgtccaccactctactccctccaccctccctccctga | 0.084 | 0.12 |
| | | | | | | TAGcagggggccgggacagggaggttgggagggggacagaggggagacagaggcacggagagaaaggaatgtttagcacaagacacagcggagctcgggattggctaaactcccatagtatttatggtggccgccggcgggggccccagcccagcttgc | | |
| RT_004627.1 | AES amino-terminal enhancer of split | LGGGTEGRQR | NP_945320.1, N | XP_00672 | 402 | aggccacctctagctttcttcctaccccattcccggcttcctcctcctcccctgcagcctggttaggtggatacctgccctgacatgtgaggcaagctaa | 0.232 | 0.107 |
| | | | | | | TAGttgcccagcgggagagacacccccatccccacgcgggccgggcacgaccacaccgggcggaccgctggacagacgcacccactcacctctccgatccgcaccccgccccacgggagcacttcctacccccacgagggcacagcccccacgccttcca | | |
| RT_024040.1 | AGAP2 ArfGAP with GTPase domain, ankyrin repeat and PH domain 2 | EGLVHGR | NP_055585.1, N | XP_00526 | 6563 | gaagacacaaactcacctccctctccccaacgaggcatggagaccccagctcaacacgcccccctctccattgtttccacactcggattgctctgggtctacccgctcggggttcgcggaaggggaggtccccagcggtacgggctcctaggcgcttggacatgcccgcttgtgccccctaggggcggaaag | 0.059 | 0.072 |
| RT_009047.1 | AK1 adenylate kinase 1 | QRWSRFPSSEPRPTPS | NP_000467.1, | XP_01151 | 5273 | TAGcaacgctggagccgcttccccagctcagagccccgccccaccccgtcctga | 0.054 | 0.424 |
| RT_002208.1 | AKAP8L A kinase (PRKA) anchor protein 8-like | PELGAGGAR | NP_001278407. | XP_00525 | 924 | TGAcccgagctcggggcgggcggagcccgcgtggccgaagctggaaaccaaacctaa | 0.42 | 0.432 |
| RT_017612.1\| | alpha-(1,3)-fucosyltransferase 9 | NFSSLAHLINILMR | NP_006572.2 | XP_01153 | 652 | TAAaatttttcatcacttgcacacttgataaatatttgatgagatatcatccaagtattgaggataagaagagatgcaacatactacttttgtgtcacaatttatttttatcaccctctctagggtaa | -0.081 | -0.089 |
| RT_022128.1 | AMOT angiomotin | SRAADYQQK | NP_573572.1, N | XP_01152 | 3141 | TAAacggccaaatcaagagctgcagattatcagcaaaaatgcttttaa | 0.051 | -0.11 |
| RT_021756.1 | ANKLE2 ankyrin repeat and LEM domain containing 2 | EGRVICLLLNCQK | NP_055929.1, | XP_00526 | 2085 | TAGgcttggcgctgggctctcggtttgttcttcatttttaaagaaggaagggtcatatgtttattgctaaactgtcaaaaaggaatatattctgattaa | -0.028 | -0.215 |
| | | | | | | TGAgccatagccccgggagggctggccaggccactccctgcccgcttttgtaatttatttatttataaactctctgctgctgagcttggggcctggagccccaggaatgagcaggcaggggagactgagatggaaataaagagactgtcgcagcagggctgct | | |
| | | | | | | ctgctcactgggctggagcggtggggaccctgaaggtggggcgctgtggagaggcactgtggatggcggcctcagacctg | | |
| RT_002681.1 | ANKRD13D ankyrin repeat domain 13 family, member D | AIAPGGLARPLPAR | NP_997237.2, | XP_01154 | 10572 | ggcttgaatcctggtcatggtccccttagcctatctgagcctcagtttcttcagttgtaa | 0.041 | 0.066 |
| RT_001281.1 | ANO10 anoctamin 10 | GLCLCHTCHGGRGGTRGSIVAPEPF | NP_001191762. | XP_01153 | 7239 | TGAgtgcccagcgtgcccagctgccctgttggcagaggcctgtgtctgtgccacacctgccacggtggcaggggggtacccggggcagcatcgtggctcctgaacccagacccaatgcttag | 0.236 | 0.257 |
| RT_005833.1 | APLP1 amyloid beta (A4) precursor-like protein 1 | AQTSPLPGAPEPQLPA | NP_001019978.1, NP_005 | | 2785 | TGAcccggccccccttcacccccttcagccgagcccagacctcccctcttcctggagccccagaaccccaactcccagcctag | 0.271 | 0.432 |
| RT_021610.1 | APPL2 adaptor protein, phosphotyrosine interaction, PH domain and | LTCACGGR | NP_060641.2, N | XP_00671 | 20075 | TAActcacttgcgcctgtgggggaagagcaaacaggaaggagagctacctcctaagggttttaacgtctctgacatacaggcacactgacctgatttccgaaggctgacaatcgtttgtggaatgtaa | 0.027 | 0.147 |
| | | | | | | TGAgccctggggagctccaaggctgcagccaggagccccgtagccagggatggtcctggctgtgctgctgcaccagggcc | | |
| RT_023876.1 | AQP12B aquaporin 12B | ALGSSKAAARSPVAR | | XP_01150 | 9755 | gccttccccaccttttccagaggaacctgttctacggccagaagaacaagtaccGAGCACCCCGAGGGAAGCCGGCCCCGGCCTCAGGGGACACCCAGACCCCTGCAAAGGGGTCCAGTGTCCGGGAGCCTGGGCGCAGTGGTGTTGA (alternative spli | 0.214 | 0.066 |
| RT_007575.1 | ARHGAP17 Rho GTPase activating protein 17 | RTWRPPLK | NP_060524.4, N | XP_01154 | 3300 | TGAagaaagccctttcccagccctccaccacttccaccctggcgagtggagcaggggcaggcgaacctctttctttgcagaccgaacagtgaaaagctttcagtggaggacaaaggagggcctcactgtgcgggacctggccttctgcacggcccaaggagaacctggaggccaccactaaagctgaa | 0.319 | 0.437 |
| RT_018159.1 | ARHGEF1 Rho guanine nucleotide exchange factor 1 | IGGLFFNIYYLR | | XP_00525 | 8679 | TGAattggaggtttatttttaatatatattatctaagaagagatctgtgtgtgtga | 0.186 | 0.382 |
| RT_023604.1 | ARPP21 cAMP-regulated phosphoprotein 21kDa | KLQMERNSK | NP_001254545. | XP_00671 | 4165 | TGAaaactgcaaatggaaaggaattcaaaagaatttagattaaaagttaaataa | 0.28 | 0.373 |
| RT_026723.1 | ASB14 ankyrin repeat and SOCS box containing 14, predicted transcri | SSSAWEAEAGGLLHLGRSR | | XP_00526 | 2744 | TAAtcaaaccattctaatggaaaaggtataattttgcagctgtattccttaaaatttcacttatgtatgattcatatttttgctttacccaccaactaaaatctcaatgcagtctaagtcctttaaaaagtagctctgcttgggaggctgaggcaggaggattgcttcaccttgggaggtcaaggctgcagtga | 0.161 | 0.151 |
| RT_011768.1 | ASPH aspartate beta-hydroxylase | KTAPTTTGGPA | NP_001158227.1, NP_001 | | 5997 | TAAagcttcaaaaagactgccccctaccaccacaggaggaccagcctaac | 0.123 | 0.033 |
| | | | | | | TGAgggagggtattgccctccgtgagcacagagactctccatgggagggggagcagtattctcctggatcctggggcctg | | |
| RT_021372.1 | ASTN2 astrotactin 2 | GRVLPSVSTETLHGR | NP_054729.3, N | | 235 | ggtgggctgggggacagctgaggatgggcctagcagatga | 0.001 | 0.028 |
| RT_013285.1 | ATG9A autophagy related 9A | DSSLIPR | NP_076990.4, NP_001070 | | 11636 | TAGacaaggctgagcagggttcctgtggcccaggatggaggccaccgctgccctgccatcccgtctgcctgccatgggacggctcctctgagtgttccctggccccacgtgtgtggtgtttgtgtgtctgtgcctggccaagggaggtgccaacactgggcttgccacagccccaggagaggaatttgggg | 0.105 | 0.162 |
| RT_006517.1 | ATP10D ATPase, class V, type 10D | DFEEVSLQAR | NP_065186.3, | XP_00524 | 2260 | TAGataccctccttggagttgcaagtattctttcaaggttggaagagggattttgaagaggtatctctccaagcaagaatgacttgtttttccataagggacatgagcattttactaggcttggaagagctgacatga | 0.203 | 0.444 |
| RT_003127.1 | ATP11C ATPase, class VI, type 11C | QNPNLELPMLLSYK | NP_775965.2 | XP_01152 | 3127 | TAAcagaatccgaatcttgaactgcctatgttattgtcctacaagcatactgacagtggttacagctaa | 0.129 | 0.152 |
| | | | | | | TGAaagtgccttcctgagatggcagtgctggtacccactgcccaccctggctgccgctgggcgggaaccccaacagggcc | | |
| | | | | | | ccgggagggaaccctgccccaacccccacagcaaggctgtacagtctcgcccttggaagactgagctgggacccccac | | |
| | | | | | | agccatccgctggcttggccagcagaaccagccccaagccagcacctttggtaaataaagcagcatctgagattttaaaa | | |
| RT_007887.1 | ATP13A1 ATPase type 13A1 | CLPEMAVLVPTAHPGCR | | XP_01152 | 666 | aaaaaaaaaaaaaaaaa... | -0.032 | 0.056 |
| RT_027025.1 | ATP6V0B ATPase, H+ transporting, lysosomal 21kDa, V0 subunit b | LGVQGPPCTPLLLR | NP_004038.1, NP_001034 | | 2970 | TAGatgatatgtgtgggtggggccgtgcctcacttttatttattgctggttttcctgggacagctggagctgtgtcccttagcctttcagaggcttggtgttcagggccctccctgcactcccctcttgctgcgtgttgatttggaggcactgcagtccaggccgagtcctcagtgcggggagcaggctgctgctgc | 0.123 | 0.127 |
| | | | | | | TGAccgggaaaacagcccctcctcttctttctccttccgagttcgcccttcccccacctccttgtctttccccgaccgag | | |
| RT_025167.1 | BCORL1 BCL6 corepressor-like 1 | LQNEAIIR | NP_068765.3 | XP_00526 | 6940 | caccagactgcagaatgaggcaataatacggaccaacaagaagccgccttatcaatgccagcattag | 0.176 | 0.341 |
| RT_020389.1 | C10orf105 chromosome 10 open reading frame 105 | WGVHKHAPTSP | NP_001161862. | XP_01153 | 9533 | TAAtggggtgtccacaaacatgcccccacatcccctag | 0.056 | -0.113 |
| RT_023564.1 | C1GALT1 core 1 synthase, glycoprotein-N-acetylgalactosamine 3-beta | KKIMNEQR | NP_064541.1 | XP_00524 | 1008 | TGAaagaaaatcatgaatgaacaaaggtaa | 0.112 | 0.056 |
| RT_024012.1 | C1orf186 chromosome 1 open reading frame 186 | IPNIFNGVQFSMDSYI | NP_001007545. | XP_00527 | 10908 | TGAattccaaatatttttaatggggtccagttctctatggattcttacatttaa | 0.197 | 0.382 |
| RT_021492.1 | C1S complement component 1, s subcomponent | YIPPASPR | NP_958850.1, N | XP_00525 | 4939 | TAAtccagatacatcccaccagcctctccaagggtggtgaccaatgcattaccttctgttccttatgatattctcattatttcatcatga | 0.001 | -0.075 |
| RT_007816.1 | C22orf15 chromosome 22 open reading frame 15 | GDGLHTVVR | NP_872326.2, | XP_00526 | 8506 | TAAggggatggattgcacactgtagtgagacatccatcctgac | 0.116 | -0.15 |
| RT_019768.1 | C22orf23 chromosome 22 open reading frame 23 | LTAPSAEK; QSPSIPTRTCV | NP_001193991. | XP_00526 | 1593 | TAAtctgtctccagagacgggcagggtagcccaagctcgaccactggagtccactcctctgccaggcagggccatcctcaggccagcccatcctaccagccttctgccacgtcaggtggtggcatagcactgccctgcagtaccccaaggcatactcagcagcctagcccctgccccggaatgcagatgg | 0.131 | -0.134 |
| RT_013060.1 | C5orf60 chromosome 5 open reading frame 60 | VTGITGLCHR | | XP_01153 | 7069 | TAGggagaccccacatgcaggcaggtggaggcaggggtcccgctttcctcagcctggatgtgccaacgctgctggacgcataagacaccaaaatcccagacactcccatggacaccgaatatccgacatccacgaacaattgggaatctcctgaggacaacttggagatggagtctcactctgttgctc | 0.364 | 0.122 |
| RT_024351.1 | CACNA1A calcium channel, voltage-dependent, P/Q type, alpha 1A su | THPTRGAAQRPR | NP_001120694.1, NP_075 | | 2949 | TAAgcccgggcgaggtggcgcccgcccggcccccccacgcaccccacgcacacaccccacccgaggagccgcgcagaggccgcgggggcccagcacagagggcccgggagagggccagccgggagaccccagactctggagaggccagggctgggccacaagggtgtcccgcagagaccctc | -0.086 | -0.112 |
| | | | | | | TGAggcagggactcggtccccttgccgtgctcccctccctcctcgtctgccaagcctcgcctcctaccacaccacacca | | |
| | | | | | | ggccaccccagctgcaagtgccttccttggagcagagaggcagcctcgtcctcctgtcccctctcctcccagccaccatc | | |
| | | | | | | gttcatctgctccgggcagaactgtgtggcccctgcctgtgccagccatgggctcgggatggactccctgggccccaccc | | |
| RT_018787.1 | CAPN1 calpain 1, (mu/l) large subunit | QPRPPVPSPPSHHR | NP_001185797. | XP_00671 | 1683 | attgccaagccaggaaggcagctttcgcttgttcctgcctcgggacagccccgggtttccccagcatcctga | 0.169 | 0.119 |
| | | | | | | TAGgcctctccccagatatctgagcataactgggagcacctcatttgtggaaaagcctctttgttatcggccttgtgtca | | |
| RT_005409.1 | CASK calcium/calmodulin-dependent serine protein kinase (MAGUK f | ASLLSALCQQVMVPR | NP_001119527. | XP_00672 | 1200 | gcaggtcatggtccctagagactacctagttgtagtgtga | -0.161 | -0.156 |
| RT_010840.1 | CCDC58 coiled-coil domain containing 58 | REILFFFFFFFFLRTGSSHK | NP_001017928. | XP_00524 | 3768 | TAAagagagattcttttttttttttttttttttttttttaaggactgggtcatctcataagagctaa2 | 0.285 | 0.259 |
| | | | | | | TGAggggtccagaagcttcgaggcccagcgacctcagtgggcccagtggggaggagcaggagcctgagccttgggaacat | | |
| RT_004671.1 | CCL3 chemokine (C-C motif) ligand 3 | GVQKLR | NP_002974.1, N | | 287 | gcgtgtgacctccacagctacctcttctatggactggttattgccaaacagccacactgtgggactcttcttaacttaa | 0.291 | 0.028 |
| RT_012415.1 | CCNH cyclin H | PFEVDFSMLTNQEK | NP_001230.1, NP_001186 | | 7561 | TAAccatttgaagttgatttctcaatgctaactaatcaagagaagtag | 0.402 | 0.48 |
| | | | | | | TGAaactggactgttcctgggaaaagggtgaagccacctctagaagggactttggcctcccccaagaacttcccatggt | | |
| RT_004395.1 | CD72 molecule | NWTVPGK | | XP_00671 | 1058 | agaatggggtgggggaggagggcgcacgggctga | 0.231 | 0.147 |
| | | | | | | TGAggcagctgctatccccatctccctgcctggcccccaacctcagggctcccaggggtctccctggctccctcctccag | | |
| RT_016795.1 | CD82 molecule | TLLPILTESRGLSGA | NP_002222.1, N | XP_00671 | 4030 | gcctgcctcccacttcactgcgaagaccctcttgcccatcctgactgaaagtagggggctttctggggcctag | 0.065 | 0.119 |
| RT_006924.1 | CDH23 cadherin-related 23 | FILEGGK | NP_001165402. | XP_01153 | 7762 | TAGgtgggggcaggtgagggtggaaaagaggtcagggctctactgttgggctttagcctctggtggtgcctcccgaggatttgctcctggctcttcccaagggctttgcagctggatcactctggactggctccctggggacctcctgaacctgttggttgcagggacggggagcatctaccaaggttcattc | 0.136 | 0.035 |
| | | | | | | TAAcaggaagcgcctcggcctcagcgtctggacctatccggccactgcagagcacccgcttctccctggccttcatcccg | | |
| | | | | | | agttgcactaaccatcctgggcttcctgtcctgtgtcccttggtgggtcccctccaggaaccaaggagtggccctccagg | | |
| RT_019066.1 | CDK2AP2 cyclin-dependent kinase 2 associated protein 2 | LDLSGHCRAPASPWPSSR | NP_005842.1, N | | 4819 | tggcagcactaaggacacccccccacaacaagagttag | 0.153 | 0.152 |
| RT_008545.1 | CEACAM4 carcinoembryonic antigen-related cell adhesion molecule | KEGALWLFWTHLG | | XP_01152 | 3977 | TAAaggaaggagccttgtggttattttggacgcacctgggataa | -0.015 | -0.122 |
| RT_022767.1 | CEP57L1 centrosomal protein 57kDa-like 1 | QNSKTVTLMNFVSETLNCLKWF | NP_001077004. | XP_00526 | 8129 | TAAcaaaacagcaaaactgtcaccttaatgaactttgtcagtgagaccttgaattgtctaaagtggttttaa | 0.29 | 0.468 |
| RT_002593.1 | CFAP54 cilia and flagella associated 54 | ISYTRDDKTEK | | XP_01153 | 1470 | TGAatatcctatacacgggatgataaaactgagaaataa | 0.333 | 0.666 |
| RT_023936.1 | CFHR3 complement factor H-related 3 | GSIVTLNVCPTSTFPLLTLMVSK | NP_066303.2, NP_001160 | | 9546 | TAAggcagcattgttaccctaaatgtatgtccaacttccacttttccacttctcactcttatggtctcaaagcttgcaaagatagcttctgatattgttgtaatttctactttatttcaaagaaaattaatataa | 0.059 | -0.059 |
| | | | | | | TGAgcagaaaatgagctgagcttagctctgccctggaacctaccagagcagagaagggcaggagaggaagatttgtctac | | |
| RT_003463.1 | CHRNA1 cholinergic receptor, nicotinic, alpha 1 | AENELSLALPWNLPEQRRAGEEDLS | NP_000070.1, N | | 15187 | ttgctccactcgcacttatcaaacgtgttatattccatacttattattgatgataagatttacctttatgtaa | 0.219 | 0.292 |
| RT_021669.1 | CLEC12A C-type lectin domain family 12, member A | SVGGGGSRL | NP_001193939. | XP_01151 | 2968 | TGAggcatcaatcaaatacatttaaggagtgtagggggtgggggttctaggctatag | 0.238 | 0.119 |
| | | | | | | TAAgcaggccagccgggctgtgcacctaaatgtctgtctgggaggagcaggctgagaagtcttgcagtctgcaggacacc | | |
| RT_010570.1 | COG3 component of oligomeric golgi complex 3 | SLAVCRTPR | NP_113619.2, | XP_01153 | 3531 | gaggaatcgtatgtgggaacgtccccgagaaccacacgagcgtgctgctcagtgctga | 0.213 | 0.114 |
| | | | | | | TAAactccctccatcccaacctggctccctcccacccaaccaactttcccccaacccggaaacagacaagcaacccaaa | | |
| RT_016946.1 | COL1A1 collagen, type I, alpha 1 | TPSIPTWLPPTQPTFPPTR | NP_000079.2 | XP_01152 | 3390 | ctga | 0.046 | -0.078 |
| RT_004138.1 | COL4A4 collagen, type IV, alpha 4 | RMRNSPTR | NP_000083.3, | XP_00524 | 886 | TAGagaatgcgaaattcaccaacacgtggccaagagaaacttcctagggggctaagacttcctagactgtgctaag | 0.2 | 0.215 |
| RT_014034.1 | CORO2A coronin, actin binding protein, 2A | AETSALLTLR | NP_438171.1, N | XP_01151 | 20475 | TGAgcagagacctctgccctcctcaccctcagggacaccactcggctccatggggaggtttag | 0.259 | 0.292 |
| RT_024782.1 | CPS1 carbamoyl-phosphate synthase 1, mitochondrial | HPSPIIK | NP_001116105. | XP_01150 | 5966 | TAGagatgcagacaccccagccccattattaaatcaacctga | 0.222 | 0.215 |
| RT_011692.1 | CREB3L4 cAMP responsive element binding protein 3-like 4 | AGTDLPGPLPDHK | NP_570968.1, N | XP_00671 | 671 | TGAgctggaacagaccttcctggcccacttcctgatcacaaggaatcctgggcttccttatggctttgcttcccactgggattcctacttaggtgtctgccctcaggggtccaaatcacttcaggacaccccaagagatgtcctttagtctctgcctga | 0.075 | 0.008 |
| | | | | | | TGAggggccgcggccctggggcgggcagccaggctgggccgcaagggcctggacggggtcggacggggcaggcgctgcggacaccgtggcctgggcttggcccgcggcctgcaccgacccgggccggacctgagcccgcagcccagacccccctccacggtccgcggcctgcaccgacccgagg | | |
| RT_021761.1 | CUX1 cut-like homeobox 1 | GAAALGRAAR | NP_001189472. | XP_01151 | 2074 | cagggcaaaatcgccatag | 0.237 | 0.028 |
| | | | | | | TGAtgcgcccggccggccctgggggttcgcggggtgtcttcttgcctgcccctgctgaggcgtcttcaggactgcaggct | | |
| RT_013213.1 | CYB561 cytochrome b561 | RLQDCRLR | NP_001906.3, N | XP_00525 | 12262 | ccggagagtggctctggcagcaggcgggcgcgtgggtgcagctgcatctgtttga | 0.292 | 0.156 |
| RT_019282.1 | CYB5D2 cytochrome b5 domain containing 2 | AVASSVNNTQRALPST | NP_001241685. | XP_01152 | 4632 | TAAgccgtagcctcttctgttaataacacacagagagctctgccaagcacctga | 0.057 | -0.112 |
| RT_011989.1 | CYGB cytoglobin | APVNQKHMAQVPQHAARPALPTLP | | XP_01152 | 12296 | TGAccacactggccccgggctggctcctgctcccctccccctctcccgtcagcaccaccccaagagagtggagtcctcgggaactgctggctggagccaagccaactccagatgtgcacagctgggccctggcctgcctgagacctgggagtcaatgggaaacaagcagatgttggcccaccagagg | 0.393 | 0.658 |
| | | | | | | TGAcctggggaactagccacacaggaggccgggagacagggactggtgagaatggggctgagtggaggaggtggtgatat | | |
| RT_016292.1 | DAAM2 dishevelled associated activator of morphogenesis 2 | PGELATQEAGRQGLVR | NP_001188356. | XP_00671 | 12479 | ttaa | 0.226 | 0.373 |
| | | | | | | TGAatctttcctgttgaagaaatccatgttatagaaaagaactttgcagtcagacattcgtcatgggaaagttcagaaaa | | |
| RT_018283.1 | DACH1 dachshund family transcription factor 1 | VLLRELPEFCVLMFFK | NP_004383.4, N | XP_01153 | 5689 | aaataaagtccttttaagggaacttcctgaattttgtgtattaatgttctttaaaagtttaagtattctacaaaaaaaaaaaaaagttttctccattgattttcacctgtggttcataccagagacctgagaatgtttgtaaatgtacaagtatcaaagttcttacagttaattactgcaacttgctgctggacaattgta | 0.121 | 0.44 |
| | | | | | | TAActctcagcctcaagggaaaacttcctcctagtgcagccctatgctttaaaaacagttcctggtggcctttctgaaag | | |
| | | | | | | gctgggtcccaggttgtcacggtgcggaactggaggccgcggtggcttctggccgaggctgggctcttccctggatgagg | | |
| | | | | | | acctgggagccgcctgggaggacagccccagaaagggagcccgagaccaggcgtgtcgccgacatgcaaatgggttgttt | | |
| RT_013666.1 | DENND2A DENN/MADD domain containing 2A | LGSSLDEDLGAAWEDSPRKGAR | NP_056504.3 | XP_00525 | 6495 | tggtggttgggtttttttttttatcttagatattaa | 0.12 | 0.536 |
| RT_013306.1 | DGKD diacylglycerol kinase, delta 130kDa | RLASWALLAWPR | NP_690618.2, N | XP_01151 | 2001 | TAGcctctgtcctctcagcctgtggcctccacatccccgccgccgaggcctagcctccgccctctcagcctgtggcctctgcgcctcctgccactgaggccctgggcagatgctgcagcccgccccccttctcatggtgctacttcctctgtcagctacagaaagcctccgtgacaccgtccaccagagctctggg | 0.227 | 0.151 |
| RT_022377.1 | DHODH dihydroorotate dehydrogenase (quinone) | LSSVPWSDHK | NP_001352.2, | XP_00525 | 13137 | TGAggacagcgtctgacgggaagcctgatctggaaccttcccaaggactcaggcaagcctttgtggctggatcatgagaggagggactccatcttgagccatgtcccccagccatggcatggctgcactgtaaacgccaatcggggggtcaccaggatcaaccgcaggctttcttcagtcccttggtcag | 0.035 | 0.345 |
| RT_007395.1 | DOCK11 dedicator of cytokinesis 11 | GNADVRDNETDLSQEYLELCK | NP_653259.3 | XP_00526 | 4740 | TGAggaaatgcagatgtacgtgacaatgagactgacctttctcaggaatatttggagctgtgcaaatgttaa | 0.306 | 0.345 |
| RT_023489.1 | DOCK4 dedicator of cytokinesis 4 | TFSVGTLITYSAPSMNGIK | NP_055520.3, | XP_01151 | 786 | TAAgtcacttttctatgtacctgcgatgcattctttgcccgtttacaaaataagaagtatgatgagaagacatttagtgtaggcactttaataacttactcagctccttcgatgaatggaattaaaacttgcttattaaatatcatgttgcacaatattaaaagttgctga | 0.054 | 0.17 |
| | | | | | | TAAgaaaagccatcttcattcgtggagactgtggccctgcaaccctggagaaggacttgctggtacttaaaaaatgggac | | |
| RT_013182.1 | DOCK8 dedicator of cytokinesis 8 | EKPSSFVETVALQPWR | NP_982272.2, N | XP_01151 | 2611 | atttgccacccaggactga | 0.054 | 0.103 |
| | | | | | | TAGggccccaccctcacccggagctcccggaggactcttgagggccgctcctcccccgcagagctttggcgtctaaaccg | | |
| | | | | | | ggtgtgtgtaaatcggtgtcatcgcttgtgtcttgctgggcgcatggttgctgtcccccctcctccgtctgggaccgagga | | |
| RT_017352.1 | DOHH deoxyhypusine hydroxylase/monooxygenase | SSPAELWR | NP_001138637. | XP_01152 | 1607 | gctgccgttgtgctgtgactccctga | 0.047 | -0.103 |
| | | | | | | TAGaatatcaggaaaaaggccacatctatggtaattaatggcagaaaagctggagagttggattctgcggtgctgctgac | | |
| RT_009530.1 | DPPA4 developmental pluripotency associated 4 | ATSMVINGRK | NP_060659.3 | XP_01151 | 23756 | aggtgaactctggtcctctgcacctgtttatgggccatgcagactggtggggtggcagatgttagcctaa | 0.046 | -0.134 |
| | | | | | | TGAagaaagaagaattcagtctcactcagtgaatgtcgcaggccatttctaaaagtgctacagaggacagacagggtttt | | |
| RT_024102.1 | DRAM1 DNA-damage regulated autophagy modulator 1 | NSVSLSECR | NP_060840.2, | XP_00526 | 626 | gaggccaccctga | 0.267 | 0.437 |
| RT_014648.1 | DUSP18 dual specificity phosphatase 18 | AIPRAPALESEVQIYC | NP_689724.3, N | XP_00672 | 4973 | TGAgccatcccacgagcccctgcattggagtcagaggtacagatctattgttga | 0.109 | 0.066 |
| | | | | | | TAAcgcaagcgacctgctgccttccttcccactgcttgtcttcagtgtgcccggctgggcaggggtgcggtggtggtggc | | |
| RT_024715.1 | DUSP22 dual specificity phosphatase 22 | LGRGAVVVADEDR | NP_001273484. | XP_01151 | 3721 | cgatgaggacaggaaagggagatag | 0.23 | 0.307 |
| | | | | | | TAGgaggacaggcctgcagcatcctggtctcgggaggcttctgtcattgctcacacacagttcagatttccacctcttta | | |
| | | | | | | tagacaagaagtgaatttgcctggggcagaacacccacccaaagagtccccacttaacaatacccccccccacggcaag | | |
| RT_006993.1 | EDN3 endothelin 3 | FPPLYRQEVNLPGAEHPPK | NP_996917.1, N | XP_01152 | 11113 | aatgcccaaatccgaatgaccccagttttcctaatgagtaa | 0.153 | -0.258 |

| | | | | | | | | |
|---|---|---|---|---|---|---|---|---|
| RT_023670.1 | EFCAB6 EF-hand calcium binding domain 6 | TPLLCAQRDDSHR | NP_942153.1, N | XP_00526 | 10512 | TAGacaccccctgctgtgtgcccagcgggacgacagccacagggcctgtttcaaggctaataaaatcctataa | -0.017 | 0.162 |
| | | | | | | TGAtccctcctctcaccccatgcccaatcagggtgcaaggagcaaggacggggccaaggtcgctcatggtcactccctgc | | |
| | | | | | | gccccttcccacaacctgccagactaggctatcggtgctgcttctgcccactttcaggagaaccctgctctgcaccccag | | |
| RT_001248.1 | EPHA1 EPH receptor A1 | TLLCTPENLFVLK | NP_005223.4 | XP_00671 | 5193 | aaaacctctttgttttaaaagggaggtgggggtag | 0.158 | 0.103 |
| RT_026532.1 | ERCC4 excision repair cross-complementation group 4 | TVMAVFLSHACTFQRLLAR | NP_005227.1, | XP_01152 | 10931 | TGAacagtgatggctgttttcttatcccatgcctgtacttttcagcggctccttgccagacatcataggtcattattaattattggtttgctatttcattcttttccaatgctcttaatgattgtacggtggaccagaagccaggattcctctctgaactctgcagttaggcatcacttgaacttgcctgtgcctgctctttt | 0.218 | 0.384 |
| | | | | | | TGAaagatgtatggtgatcttgaaacctccagacacaagaaaacttctagcaaattcaggggaagtttgtctacactcag | | |
| RT_003934.1 | ESRP1 epithelial splicing regulatory protein 1 | MYGDLETSR | NP_001116297. | | 8815 | gctgcagtattttcagcaaacttga | 0.122 | 0.056 |
| | | | | | | TGAcacgggccaaggagccagtggggaagtgcgtgtgttgggccaggtaggatacgtacacctcttgcctcagagcagcc | | |
| RT_002612.1 | EVPL envoplakin | TPLASEQPHPR | NP_001979.2 | XP_01152 | 9163 | tcatcccaggcagtgggtcttccctctgtccaaccactgttttattattttactaacatggtga | 0.216 | 0.421 |
| | | | | | | TAGtttctccgtccctacacagggagctcctccccaagggtagatcggaccgttcatgctgcctataggcattatgtccc | | |
| | | | | | | tcaaaaaaaaactcctttgcctgcatcctgtgtacaacatgacatttttaaccaatccaatctaaaaatgtgccagaatc | | |
| | | | | | | cacctgtggcccgaatcgtgtttggttcctctttctactccactgcagatgaccaaacctgtcccgctgccactttcctc | | |
| RT_018437.1 | FAM192A family with sequence similarity 192, member A | TVHAAYR | NP_079222.1, | XP_00525 | 8802 | actgatattgggaggagggcaaggcccagccgaagttccactaaaaatgccccaggagaatag | 0.236 | 0.104 |
| | | | | | | TGAcagagaggtgtgagggaggcgaaggccctggcctccgaggatctgggtggcgtctgctgccctaggtgtgctggaag | | |
| | | | | | | tccacttggaagaactgttccggaggcgctgggtcgggatgccgaacctctcctgatccgccggcagcaacgagccatta | | |
| | | | | | | aaactgcagttcctgaccacgcactgcttcgcaggctccgagccctgcatcctccacagcccccgccttgctcactgttg | | |
| | | | | | | ggccctcagggcgggcagggtggctgtgagtcccaggtgtcatcttcctgctctgcgactttctctggagaccttgggcc | | |
| | | | | | | tttggcctgtggggcactggggatggtaacctcactgccccgtcactcccttcaaaggcgacagacccaagcccacgtca | | |
| | | | | | | ggagaggagcgtggggtcagctccagcagggtcggggtcagtgcctgtgtctggtgggggccactcaggagagttgggtt | | |
| | | | | | | ttcagggcagagatggcttgattccacctggtggggtgggtggaccctggagactctcaggtgatgggccagcctgccc | | |
| RT_011736.1 | FAM213B family with sequence similarity 213, member B | AEMGLIPPGGVGGPWRLSGDGPA( | NP_001182667. | XP_01153 | 21033 | caaggctctttactga | 0.316 | 0.33 |
| RT_012764.1 | FANCE Fanconi anemia complementation group E | GPPSWCSITSFLK | NP_068741.1, | XP_00524 | 4680 | TGAccatctcaccaagggaccaccctcttggtgctccatcaccagcttcctgaagggcatttctttcttcaccaccttgtcttga | 0.371 | 0.658 |
| RT_025932.1 | FBL fibrillarin | SSALSGLR | NP_001427.2, | XP_00525 | 120 | TGAagttcagcgctgtcaggattgcgagagatgtgtgttgatactgttgcacgtgtgtttttctattaa | -0.004 | 0.153 |
| | | | | | | TGAggacacagacctgcttcagggccaagcctggatgggtgctcagaggagctaaggtggaaatactgcctgaaaagact | | |
| | | | | | | ccgcccctccccctcccctcgaacaaatgggtcccgggcctccaaaacacggtacctccagaacacttcagctccacag | | |
| RT_026099.1 | FBXO10 F-box protein 10 | GHRPASGPSLDGCSEELRWK | NP_036298.2, | XP_01151 | 4883 | cccatagcaagaagaaagtgggcaatgtttttcagggagaagggatagaagctggtgggtttccagtcactcgaagtctataa | 0.195 | 0.345 |
| | | | | | | TAAagtctagaggaggacattgcacctttgctgctgctgctatcttccaagagaacgggactccggaagaagacgtctcca | | |
| RT_024737.1 | FBXO21 F-box protein 21 | LHGALGTCCTRK | NP_055817.1, N | | 2900 | cggagccctcgggacctgctgcaccaggaaagccactccaccagtagtgctggttgcctcctactaa | 0.081 | -0.025 |
| RT_011219.1 | FGFRL1 fibroblast growth factor receptor-like 1 | GPMARRNGQHPR | NP_001004356. | XP_01151 | 7158 | TAGacggcaccgtatctgcagtgggcacgggggggccggccagacaggcagactgggaggatggaggacggagctgcagacgaaggcaggggacccatggcgaggaggaatggccagcaccccaggcagtctgtgtgtgaggcatag | -0.033 | -0.155 |
| RT_007474.1 | FMNL2 formin-like 2 | TCVLLMVPK | | XP_01150 | 2309 | TGAggagaagcgtcaggcggcgctttgatgatcagaacttgcgttctgttaatggtgccgaaataa | 0.136 | 0.345 |
| RT_020024.1 | FRK fyn-related Src family tyrosine kinase | TLEKNIK | NP_002022.1 | XP_01153 | 386 | TGAacactggagaagaatatcaaataa | 0.046 | 0.384 |
| | | | | | | TGAactgcagttggacctcctgaccaagcactgcgttctcacactagtgtgccttgcaaattgtatttgtcttccgagag | | |
| RT_019443.1 | FRMD4B FERM domain containing 4B | SHTSVPCKLYLSSERMFVLET | NP_055938.1, | XP_00671 | 6767 | gatgttcgttttagaaacctaa | 0.206 | 0.1 |
| RT_026450.1 | GDF9 growth differentiation factor 9 | QMVLLKP | NP_001275753. | XP_01154 | 7924 | TAAcaaatggtcctcttaaaaccttga | 0.24 | 0.468 |
| RT_014494.1 | GGT5 gamma-glutamyltransferase 5 | TVARTRPR | NP_001289393. | XP_01152 | 6149 | TAAgactgtaggaggtgcagaggggactccaagaccgtggccagaaccagacccagaggcccttcttcctga | -0.034 | -0.123 |
| | | | | | | TGAgcggtggcaccccctcctgctggtgccagtgggagactgccgcctcctcttgacctgaagcctggtggctgctgccc | | |
| | | | | | | cacccctcactgcaaaagcatctccttaagtagcaacctcagggactgggggctacagtctgcccctgtctcagctcaaa | | |
| | | | | | | accctaagcctgcagggaaaggtgggatggctctgggcctggctttgttgatgatggctttcctacagccctgctcttgt | | |
| | | | | | | gccgaggctgtcgggctgtctctagggtgggagcagctaatcagatcgcccagcctttggccctcagaaaaagtgctgaa | | |
| | | | | | | acgtgcccttgcactggacgtcacagccctgcgagcatctgctggactcaggcgtgctctttgctggttcctgggaggct | | |
| | | | | | | tggccacatccctcatggccccattttatccccgaaatcctgggtgtgtcaccagtgtagagggtggggaaggggtgtct | | |
| RT_003442.1 | GLB1L2 galactosidase beta 1-like 2 | SISLSSNLR | NP_612351.2, | XP_00671 | 5181 | cacctgagctga | 0.044 | -0.025 |
| RT_007096.1 | GLYATL3 glycine-N-acyltransferase-like 3 | PSKKLQWFYYFP | NP_001010904. | XP_01151 | 11307 | TAGcccagtaaaaaactgcagtggttttattactttccctga | 0.129 | 0.21 |
| | | | | | | TGAgcggggggcgccgtccaggccgagcgcagactgtttaggactcagcagacccagcaagaggcatctgccctttcccc | | |
| | | | | | | agccacctccccagcaagcaacctgaaatctcagcagatgcccaccatttctctagatcgcctagtctcaacccataaaa | | |
| RT_024798.1 | GPR17 G protein-coupled receptor 17 | AGGAVQAER | NP_001154887. | | 1128 | aggaagaactgacaaaggggatccatcggccacccctctgcaggggcttgtga | 0.133 | -0.025 |
| | | | | | | TGAgcatcaaacaaatctcttcactgtttctttttttaggactccctttgcaaggagcaactgtaatattgtgggactaac | | |
| RT_023626.1 | GRID2 glutamate receptor, ionotropic, delta 2 | SLHCFFFR | NP_001501.2, N | XP_01153 | 5438 | atggatgtaa | 0.19 | 0.292 |
| | | | | | | TAGgcacagctggggtggggacctcaggcctgggggctgggcagaggaaagcaaaggagattggaaggaacgtcccctgt | | |
| | | | | | | ccccgcactgggcttggggaccagagctgccacctgcctgttgggccaggagcctcctgcccttacctgccaggaagcca | | |
| | | | | | | gcaggctctcaggccagctgcttgggcttcatcctcctcagatcttctgtgggtttctaaagctgccagccgagatagcc | | |
| | | | | | | aaggccaaaggaagcacatgcctctctcaggccaaactcacctgcccctcaactctcctccagagtcagaagtttctgcc | | |
| | | | | | | gcagccctgcagagggcacagaaaatggaagacagctcttatattgccatttcttccacaagagcccaggcctcctacag | | |
| RT_000656.1 | GRIK3 glutamate receptor, ionotropic, kainate 3 | AQLGWGPQAWGLGRGK | NP_000822.2, | XP_01153 | 865 | cttga | 0.122 | 0.07 |
| | | | | | | TAGcagctactccgagcgccgtttacagttttgcacagtgatcttccccattgtccactcaagtcgtggcctggggaaca | | |
| RT_006083.1 | GRK6 G protein-coupled receptor kinase 6 | APFTVLHSDLPHCPLK | NP_002073.2, | XP_00671 | 3763 | cagacggagctgtccccagtgtcctccgtccctcagcccctggcctggctga | 0.05 | 0.107 |
| | | | | | | TAGagggatgctttaaccaaccctcttcctccccgtcattgtactgtaactgggacagaagaaataatggggatatgtgg | | |
| RT_002849.1 | H2AFV H2A histone family, member V | RDALTNPLPPR | NP_958844.1, N | | 14036 | aatttttaacaacagttaa | 0.214 | -0.102 |
| RT_012910.1 | HAGH hydroxyacylglutathione hydrolase | GRPAPSADLGIR | NP_005317.2, NP_00103 | | 2154 | TGAggccgccctgcaccttcagcggatttggggattaggctcttttag | 0.028 | 0.119 |
| RT_012805.1 | HAUS2 HAUS augmin like complex subunit 2 | SSTLFLLNYV | NP_060567.1, N | XP_00672 | 7221 | TAGtcatcaactttatttttgcttaattatgtgtag | 0.019 | 0.106 |
| | | | | | | TAAgtcaattctttgggcctactccccgactccctccctgaagatgcttctcaggctcccatgagaaaacaatctatact | | |
| RT_006180.1 | HEATR9 HEAT repeat containing 9 | QSILIFR | NP_689994.2 | XP_00527 | 5817 | tatcttccgtgaataa | 0.115 | 0.17 |
| | | | | | | TAGgcttcagctgggactgccagggccgcggccgcccgcacgtccgcgggtcccggccgcgccgccgcgcgcgcccctg | | |
| RT_023343.1 | HOXB8 homeobox B8 | ASAGTARAAAAR | NP_076921.1 | XP_00525 | 1334 | cccgagagagctctggccccgctag | -0.2 | -0.215 |
| | | | | | | TGAccggtggcttctcgctgcgcagcctccagggcccgaggtgatcccggtggtgggatccacgctgggcctaggatggg | | |
| | | | | | | acgtttttcttcttgggcccggtccacaggtccaaggacaccctccctccagggtcctctcttccacagtggcgggccca | | |
| RT_007294.1 | HPN hepsin | STGPRTPSLQGPLFHSGGPTQPRDH | NP_892028.1, N | XP_00672 | 1019 | ctcagccccgagaccacccaacctcaccctcctgac | 0.314 | 0.341 |
| | | | | | | TGAgggcgcgccggcgcggtgccagcgccgtctgcccggccccgccctctttcggttcaggggcctgcggagcgggttgg | | |
| RT_010197.1 | HSPA12B heat shock 70kD protein 12B | VGAGETIVLQSAPFHALQPR | NP_443202.3, N | XP_00526 | 3550 | ggcgggggaaacgatagttctgcagtctgcgcctttccacgccctccagccccgggggagataa | 0.142 | 0.028 |
| RT_012273.1 | IDS iduronate 2-sulfatase | VLPTMDGK | NP_001160022.1, NP_000 | | 3218 | TGAgttttgccaaccatggatggcaaatgtgatgtgctcccttccagctggtga | 0.075 | 0.289 |
| | | | | | | TAGagacctgcctccccttgcacccgaggccctgcagcagggagctcagcgaggcagagggtggggctgcacagaggg | | |
| | | | | | | gaacatcagctgcagctctgcaccaggccggtccctggggactggggcgctcctccctcaggctttctccctcagtcttg | | |
| | | | | | | gcttctccagggctctggggtgtctggagctaggcttggccctaccattctggggccatttccaccacagttggggctct | | |
| RT_004577.1 | IFFO1 intermediate filament family orphan 1 | GRGWGCTEGNISCSSAPGR | NP_542768.2, N | XP_00671 | 3036 | cctgccttcacgcgtgggtgtctgctacttccccatctttaa | -0.066 | 0.215 |
| | | | | | | TGAccggccgcatccttgcacgccccgcggagctctctccatggtagcccagggcgctggcagaccatgctccgcctcca | | |
| | | | | | | ccagggccacagaggagcggaggggcctatgggggaggagcggagggccctgttggggaaggttgggttttggacccca | | |
| | | | | | | gggataagcttttccgatgtcacaatgtggaggaaagcacctgggggacaacagtgctcgtgcaggtggggcttgggaaa | | |
| | | | | | | tgcacgtcccttccccttactccccgccaaaacccacatcccagcctctggatcctggggaaggttccagtccctggaga | | |
| RT_016397.1 | IGHMBP2 immunoglobulin mu binding protein 2 | WQTMLRLHQGHRGAEGPMGEER | NP_002171.2, | XP_00527 | 4725 | atacccagggcctcaaacttga | 0.278 | 0.341 |
| | | | | | | TAAagccctcccgcgcccccaccccagacccgagccaccccaggaaaagcacaaggactgccgccttctcgctcccgcc | | |
| RT_011586.1 | IKZF1 IKAROS family zinc finger 1 (Ikaros) | KSTRTAAFSLPPAA | NP_001278768. | XP_01151 | 10190 | agcagcatag | -0.107 | 0.033 |
| | | | | | | TGAgcagagctccaccgcagtcccgggtgtctgcggccgcaacgcaacggacactggctggaaccccggaatgagccttc | | |
| | | | | | | gaccctgaaatccttggggtgcctcgaggacgactggccgaaaagccgcattcctgcctcacaggccggaagtcccagc | | |
| | | | | | | ccagtccccgcgcgcgtccctcttcctcctcatactttcccttgactgagagctcctctaacccctgttctgatgggga | | |
| | | | | | | gggcggtcttcccacttcctctccagaactccagaaagagcagtgtgcttatgcttcagtccaggctggagaggttgggg | | |
| | | | | | | ccggggtagggaggcaggagccatgtcagttctgaaggagggtgaggcggtgggggattgcagggggcggctgagagaaa | | |
| RT_023045.1 | IL17RE interleukin 17 receptor E | IPCLTGRKSQPSPR | NP_705616.2, N | XP_00671 | 1412 | acctccttgggggccagggattccctttcccactctga | 0.154 | 0.292 |
| RT_008965.1 | IL33 interleukin 33 | RELGLPDPLK | NP_254274.1, N | XP_00525 | 770 | not found | | |
| | | | | | | TGActgtcactttggcatcctctgatgaactcacacatgcctcagtgcctcagtgaaaagaacagggatgctggctcttg | | |
| | | | | | | gctaagaggtgttcagaatttaggcaacactcaatttacctgcgaagcaatacacccagacacaccagtcttgtatctct | | |
| | | | | | | taaaagtatggatgcttcatccaaatcgcctcacctacagcagggaagttgactcatccaagcattttgccatgtttttt | | |
| | | | | | | ctccccatgccgtacagggtagcacctcctcacctgccaatctttgcaatttgcttgactcacctcagacttttcattca | | |
| | | | | | | caacagacagcttttaaggctaacgtccagctgtatttacttctggctgtgcccgtttggctgtttaagctgccaattgt | | |
| RT_025506.1 | IL5RA interleukin 5 receptor, alpha | LTSSCIYFWLCPFGCLSCQL | NP_783853.1, N | XP_01153 | 4995 | ag | 0.208 | 0.623 |
| | | | | | | TAGccaaaacagctggcaggactgccggtcacacaccagcacgtcccacctcgtgctcacggattttattacacagatag | | |
| | | | | | | tggcggcaatggcctcagcccagcccaccctcacctgcttttccagcccacaaagggggacgatcacggcccagcaaaag | | |
| | | | | | | cgatgctgagaggggaaacagtccagagtccaacagcagaacttgggggaagcggtcggggtggccaggaacataaacta | | |
| | | | | | | tgtatagggccggggccttctgcccagggctcccctggaccaggacgccaggtagggcagggaacctcagtagtcctcc | | |
| RT_020939.1 | JMJD8 jumonji domain containing 8 | GGQEHKLCIGAGGFCPGLPWTR | NP_001005920. | XP_00525 | 7736 | acccagcattctcagagatgaatgcgtcaatacctccttcatagccaagttggggatga | 0.182 | 0.435 |
| | | | | | | TGAgtccgcgcgcggggcgcatgcgcggccaccctcccccaagggcgggctcgcggggggtgtcgtgggcgccccggacttggagagggtgcggccctggggacccccctccccgagtgtgcccaggaactcagagggcgcggcccccggggattc | | |
| | | | | | | cccccccccgagggtgcccaggactcgacaagctggacccccctgctcccgggggggcgagcgcatgaccccccgccctc | | |
| RT_010302.1 | JUND jun D proto-oncogene | ARPPGIPPPRGCPGLDK | NP_005345.3, N | | 752 | gcgctgcctcttttcccccgcgcggccgcccccgtgttgcacaaacccgcgcgtctcggctgccccctttgtacaccgcgccgcggaagggggctccgagggggcgcagcctcaaaccctgcctttcctttactttttacttttttttttttttctttggaagagagaagaacagagtgttcgattctgccctatttatgtttc | 0.187 | 0.348 |
| | | | | | | TAGcgcctgcgccgcccacacggagacccccctgccccctcccagctgcagcgtcgggaccccccgaggtgcgccaagggggtg | | |
| RT_009378.1 | KCNG2 potassium channel, voltage gated modifier subfamily G, mem | RPHGDPLPPPAAASGPPR | | XP_01152 | 16826 | gggggcgtctggcctgggggagcggctcctgccggccgcgtcctcggcccctcgtgcgtgagcagccccagaacttggcggggccctgcctgactccccgtggcagcgctgggcaaagtcactggcctttgtcctctgccccaccccttccttggatttttaattttctacgcctag | 0.195 | 0.485 |
| | | | | | | TGAaaaatccttccaacattcactttattgaacagcttggtggactggaagggtccctccaagaaacaaatctgcatctc | | |
| RT_000486.1 | KCNU1 potassium channel, subfamily U, member 1 | KILPTFTLLNSLVDWK | | XP_01154 | 14563 | agcactgcctttt | 0.108 | 0.373 |
| | | | | | | TAGgccccagaggtcgcactccgcaggaggactgaggccatgtgccattctcccgggcccagcgcccggccgtcacccca | | |
| | | | | | | cccgatgacctgcatgaagccagcagcacccagagccactcctgctgccctagaactagcggttagaagaatccgctgtt | | |
| RT_003017.1 | KIAA0513 | RTEAMCHSPGPSARPSPHPMTCMI | NP_001273495. | XP_00525 | 10909 | cctccctcatctcctctgcctgtgtctgcgaccccatccatgtgccaaagtgtcccttgggtcacacagctaa | -0.046 | -0.156 |
| RT_021510.1 | KIAA1191 | SPLSRDSGK | NP_001274264. | XP_00526 | 8368 | TAGagccctctttccagggattctggtaaaggtggtttcttgcatcccactccccttttaccttggctttga | 0.014 | -0.011 |
| RT_003800.1 | KIAA1683 | DPGSLQWGLRGR | NP_079525.1, N | XP_00526 | 3408 | TAGgaccctggctccctgcagtggggacttcgtgggaggcactcatggctctgggtctaatgaataa | -0.065 | -0.167 |
| | | | | | | TAAgaagctttgaggctaccaggcaggggagtccccctacccacaacctcttccctgaaagggatgagggggaagagagg | | |
| | | | | | | tagcagccagagccaggacccagggttggggctgccggctgacccggagcccctggagcaggaggctggggcagagggcc | | |
| | | | | | | ctaggccaagcccaccctgggcaccagggacaatcctcttccccaccaccggccctcaggctggcatctctgcccccagc | | |
| | | | | | | tccaggaggggccagacagaagcagccattgggcatctcaggtttgagggggatatgggccgggaactacccagaagcat | | |
| RT_010079.1 | KMT2D lysine (K)-specific methyltransferase 2D | RGSSQSQDPGLGLPAD | NP_003473.3, | XP_00526 | 6602 | ctgggaggcagcagggtgggggaagaggatgtgtggccgggcctcacagccctgctgctcccactga | 0.021 | 0.103 |
| RT_005240.1 | LAMTOR2 late endosomal/lysosomal adaptor, MAPK and MTOR acti | NDHLEGR | NP_054736.1, NP_001138 | | 2906 | TAAcggcattggtggaagctggggtcagaaaagagaaatgaccatttggaggggcggggcctcctagaagaaccttcttag | 0.198 | 0.216 |
| | | | | | | TGAgggtgcccgggcaagtcttgccagcaaggcagcaagacttcctgctatcaagcccatggaggaaagttactgctga | | |
| RT_017114.1 | LIPG lipase, endothelial | TSCYPSPWRKVTAEDPPNGR | NP_006024.1, | XP_00525 | 5528 | ggacccacccaatggaaggattcttctcagccttgaccctggagcactgggaacaactggtctcctgtga | 0.063 | 0.028 |
| | | | | | | TAGgtggctgccaacgcgcacgctcgggtccgaggctgctcccctggagcggcgcccctgcgccctcagcccgagcagcg | | |
| RT_023299.1 | LMTK2 lemur tyrosine kinase 2 | VAANAHARVRGCSPGAAPLRPQPE | NP_055731.2 | XP_01151 | 13730 | acatccactcgccatttgctga | 0.16 | 0.035 |
| RT_001154.1 | LOC102723730 adhesive plaque matrix protein-like | SPWPTGLR | | XP_01150 | 17842 | TAAttccctatttctcagcctctgctggtgaagtttctgcctcttacttcagaagtccaagtggtgaagagcccttggcccacaggtctcagggtggtaggaacgaaaaggccgctctga | -0.025 | 0.207 |
| | | | | | | TGActggtccacgcacggccaaagcatgaggaagctcatcgtgggattcatttttcttacattttggacttacacagtaa | | |

| | | | | | | | | |
|---|---|---|---|---|---|---|---|---|
| RT_013953.1 | LOC105373926 putative HERV-K_Xq28 provirus ancestral Pol protein | GIHFSYILDLHSKGFN | | XP_01151 | 8461 | gggcttcaactga<br>TGAcggctcaccataaaagccaaaatggggacctgtcaaaaatgttttagccaaactatggcttggctcaggaagatgcc | 0.399 | 0.623 |
| RT_000939.1 | LOC643355 uncharacterized LOC643355 | KMPRPK | NP_001257874. | XP_00527 | 8225 | aagaccaaaggctgctgaactttaa<br>TGAacaacccaggccatggggtggaaacggcttgggcttcaagcctctgaaaccaaccctctgcagcagctccatccatt | 0.209 | 0.394 |
| RT_024143.1 | LRIT2 leucine-rich repeat, immunoglobulin-like and transmembrane | TTQAMGWK | NP_001017924. | | 801 | tatccagcaagtatatattgagcacccactgtgttccaagcactga<br>TGAggcggcggggccgggcgggcgaggggcgtggagcccccacccaggtcccagcccgggcgcagcctgaccgggacccctccctcccacagcccagcccaccttctgggaccacgcagggaattggggagaggtggcttccagccccatctggggctc | 0.205 | 0.384 |
| RT_007580.1 | LRRC4B leucine rich repeat containing 4B | GGGAGRARGVEPPTQVPAR | NP_001073926. | XP_00672 | 1548 | ggacccccagtaaggacagggtggggctccaggagcggagtctctgggatcctcgcctcacccccgccaatccccggtga<br>TAAattccagctctgatcaggcaactggttggtgactggagagcattgtttcataggcttttctctgtcctatctgggag<br>cgctgcttcttcccctgccttccgagagacgaagaccgtggcgagcttggcgcttaggggctcccgtgccatggctcacc<br>ccagggagccccagcagccaccaggtgcctctgtctgcagacccctggcccgggctggcgccgacgctcagaacctgcag | 0.16 | 0.119 |
| RT_026848.1 | LRRFIP1 leucine rich repeat (in FLII) interacting protein 1 | TVASLALR | NP_001131022. | XP_01151 | 10123 | gtacttcataagcacacaggggcctcgagggagctctgtgtctga | -0.019 | 0.204 |
| RT_006265.1 | MAOB monoamine oxidase B | RERVSVITLSSYCIWDMSLGK | NP_000889.3, | XP_01154 | 6600 | TAAagagagagggtgtctgtaatcacactctcttcttactgtatttgggatatgagtttggggaaagagttgcagtaa<br>TGAgagctgaatgagggcatcataggccagacaggcccaaggatggatgaatggagaggacaaaggcagcttctgacacaccagccccaggacctggggcgactggaggaagccaggcgagtggggcccaggactggttccagtgagagaaaccaaccacaggcacccaagcactaccagac | 0.26 | 0.259 |
| RT_023252.1 | MAP3K6 mitogen-activated protein kinase kinase kinase 6 | ELNEGIIGQTGPR | NP_001284538. | | 5739 | gtgggaggggtatcctgggaagagcccccctaggtctttttaaaaaatttttttgttggccgggcatggtggctcatgcctgtaatcccagcactttgggaggccaaggtgggcagatcacgaggtcaggagatcaagactatcctggctaa | 0.059 | -0.036 |
| RT_013519.1 | MAP7D1 MAP7 domain containing 1 | EGLPWIR | NP_060537.3, N | XP_00671 | 2843 | TAAgagggtttgccttggatccgggcacagttgtga<br>TAAgtagcaacgccactccttctcagtcttcatcgatcaatgacatttcatccatgtccactgagcagacgctggcctca<br>gacacagacagcagtcttgatgcctcgacgggacccccttgaaggctgtcgatgataggttagaaatagcaaacctgtcag | -0.005 | -0.214 |
| RT_011694.1 | MAPK9 mitogen-activated protein kinase 9 | VATPLLLSLHR | NP_620707.1, N | | 3409 | cattgaaggaactctcacctccgtgggcctga | 0.151 | 0.396 |
| RT_022686.1 | MARK2 MAP/microtubule affinity-regulating kinase 2 | QAARSGGGGGGPAGR | NP_001034558. | XP_01154 | 659 | TAAcaggctgccaggagcgggggcggcgggggcgggccagctggacgggctgccggccgctgcgccgccccacctgggcgagactgcagcgatggattggtgtgtctcccctgctggcacttctcccctccctggcccttctcagttttctcttacatgtttgtggggggtgggagattgttctccagcacc | 0.181 | 0.152 |
| RT_022228.1 | MCAT malonyl-CoA-acyl carrier protein transacylase | LQGAQMR | NP_775738.3 | XP_01152 | 10785 | TGActgcaggggggctcaaatgcgatga | 0.257 | 0.623 |
| RT_027052.1 | MED1 mediator complex subunit 1 | EPYFLKETGPEEK | NP_004765.2 | XP_00525 | 10723 | TAGgaaccttatttcctaaaagaaacagggccagaggaaaaaaaactattgataagtttatag | -0.112 | 0.058 |
| RT_014431.1 | MED12L mediator complex subunit 12-like | RHDVLCLH | NP_443728.3 | XP_00671 | 5306 | TGAatctgcaagaggagaagacatgacgttttatgtttgcactga | 0.257 | 0.44 |
| RT_022201.1 | MEI1 meiotic double-stranded break formation protein 1 | SSGLEGPEVER | NP_689726.3, | XP_01152 | 9595 | TGAtcctcaggacttgaaggcccagaagtggagagagaatga<br>TAGcagctacctgctcagccctgcaccatgctgctggggaagacagtggccccagccccgtcaggcagggccagggacag | -0.087 | 0.103 |
| RT_017809.1 | MGAT5 mannosyl (alpha-1,6-)-glycoprotein beta-1,6-N-acetyl-glucosa | SHAGTLARA | NP_002401.1 | XP_00526 | 2131 | aagtcatgcagggactctggcaagagcctga | 0.125 | 0.107 |
| RT_013659.1 | MID1IP1 MID1 interacting protein 1 | KMVAGDGR | NP_067065.1, NP_001092 | | 1487 | TGAggcgtggcgcccgtggctgcccagcaccttcttcgacccatctcaccctctctcattcctcaaagcttttttttttttcctggctggggggcgggaagggcagactgcaaactggggggctgcgtacgtgcaggaggcgcggtggggctgcgtggaggaggggggccacgtgtgagagagaagaaaa | 0.154 | 0.119 |
| RT_001122.1 | MIS12 MIS12 kinetochore complex component | LLSSQKEEPVK | NP_001245146. | XP_00525 | 2641 | TAAttgctcagtagtcaaaaggaggagcctgtcaaaaagtag<br>TAGcgaatcctccctgcactgcattcgcacaaactgcttcctttcttgattcgtagatttaataacttacctgaaggggt | 0.066 | 0.116 |
| RT_019879.1 | MITF microphthalmia-associated transcription factor | RILPALHSHK | NP_937802.1, N | XP_01153 | 3922 | tttcttgataattttcctttaa | 0.278 | 0.488 |
| RT_021866.1 | MPV17L MPV17 mitochondrial membrane protein-like | EVKDSLK | NP_001121895. | XP_01152 | 10879 | TGAgaagtcaaggactctcttaaagggaccacatttttacctaa | 0.219 | 0.281 |
| RT_021358.1 | MSL3 male-specific lethal 3 homolog (Drosophila) | NVVGSVR; ATALSSLALWVPGE | NP_523353.2, N | XP_00527 | 1444 | TAAaatgttgttggttctgtaagagcaactgctctgtctagtttggcgctctgggttccaggtgaataa | -0.003 | -0.089 |
| RT_010471.1 | MTR methionine synthase | FAVLGTMVMK | NP_000245.2, N | XP_00527 | 10332 | not found | | |
| RT_020347.1 | MTSS1 metastasis suppressor 1 | VHKKCAGGE | NP_055566.3, N | XP_00671 | 4627 | TAGgttcacaagaaatgcgccggtggggaatga<br>TGAgagccacagctgacaaagtctgcatgtccgaggacggcccctgcactggagctgggcgccagagctgcagagctagt<br>gttcggccctcagagaaggatccagaatcaaaagctcaagagtgacgtgaggtgggcaccggccccaagtgcagagtcaa<br>ggcagggagaggccggctggagccaggcccccctcgcacgcagcccccaaatcatggacgcacctgtggggagcaccacat | 0.134 | 0.067 |
| RT_025500.1 | MYO9B myosin IXB | LKSDVRWAPAPSAESR | NP_004136.2 | XP_01152 | 1548 | ctccacctgcggcctcacatctccccactcccctttttgtacgtttaa | 0.079 | -0.036 |
| RT_006029.1 | NCAM2 neural cell adhesion molecule 2 | QQYYRGLNNTTK | NP_004531.2, | XP_01152 | 14703 | TAAcaacaatattacaggggcttgaacaacactacgaagagtatttggattgcgtga<br>TGAgtggggagcaccgaggcaggggtgggggaatgtgtactgaggagccgtgtgtctgctcctggctggcccggcctaat<br>aaagcagtgttgccatctcatcttcccctaaaaacccttttatgtacacctgcgcagagaagagggctgcctggcctccctgggccgctggtacagatcacacacacacacagattaaacgcagcccgtttaatgatggggcccagactgcagtggctcaagacaggacactgcggaaagccgccatgt | 0.29 | 0.468 |
| RT_009713.1 | NCAPH2 non-SMC condensin II complex, subunit H2 | LNAARLMMGPR | NP_001171940. | | 1370 | taatccgtga<br>TGAggggctgcgccacgccacggccccgctggagctggggaccacagactggaccggctctcttcatgcccagccccgg | 0.233 | 0.257 |
| RT_014759.1 | NCKAP5L NCK-associated protein 5-like | LSSCPAPGDGDPSLK | NP_001032895. | XP_00671 | 3669 | agacggggacccctccctgaagggaccaaggaggcaggtggataagaaggtgaaaagggggtccctggcacaccccaccacccactgcttcggcggatga<br>TGActcagaacagggcggggggggggggcggtgtcaggtcccagcgagccacaggaacggccctgcaggagcagggcggctgccgactcccccaaccaaggaaggagcccctgagtccgcctgcgcctccatccatctgtccgtccagagccggcatcctt<br>gcctgtctaaagccttaactaagactcccgccccgggctggccctgtgcagaccttactcaggggatgtttacctggtgctcgggaagggaggggaaggggccggggagggggcacggcaggcgtgtggcagccacacgcaggcggccagggcggccagg | 0.164 | 0.028 |
| RT_015970.1 | NCOR2 nuclear receptor corepressor 2 | HPCLSKALTK | NP_001070729. | | 13497 | gacccaaagcaggatgaccacgcacctccacgccactgcctcccccgaatgcatttggaaccaaagtctaaactga<br>TAGgcatttcaaagcatcacacttgtctgcaggacttacagatcctgcagtcaatgtttcggtttagactctccactgtt<br>acctaagttctcaagctgcctatggtttttctgtgtcaatgtgatttatggtagtaccatcctttctcctttgggtttta<br>aaataagttgcagaacagacactttaaaagcttctgcaatattatttctgtgcctagagtctttctccattataaacatg | 0.265 | 0.714 |
| RT_006054.1 | NEB nebulin | AFQSITLVCR | NP_004534.2, N | XP_01150 | 727 | ttttaa<br>TAAgatttgagtccattgcaaaaggttaagccatatgacaatttcaaaatgcatgtgattggcagcttcaaaacagaacg<br>ggttctcccatgggggctccagacattgtattttactttgtgcaatatgaggggactgcatgcaagctcagggtgctccc<br>tcctcagtctttgggggattcaaatgcatgatattgtatgtacctgggaaatttgccgatttcctaagctgttggaaggg | 0.092 | 0.07 |
| RT_001896.1 | NEFM neurofilament, medium polypeptide | VLPWGLQTLYFTLCNMR | NP_005373.2, N | | 19012 | ggtcacttaagggggggatgtcttgagatgtattatgcaaagtaccaactga<br>TGAccagccccacccccctccccagcagccagagaagcagtggcccggggcagttccgagggcagtgcctgcagtcaagtg | -0.018 | -0.182 |
| RT_019335.1 | NFASC neurofascin | QWPGAVPRAVPAVK | NP_001005389. | XP_00524 | 284 | gccgggtcaggcgtggtga | 0.407 | 0.658 |
| RT_016847.1 | NFIC nuclear factor I/C (CCAAT-binding transcription factor) | LPKLFISSPPAAR | NP_995315.1, NP_001231 | | 2713 | TAGcaaaggtcttcttccctcgccccttctccatcgtcccaggaatcccagggggcagcacagccggcccccggcccacgttttcggtggaaaattagagtgaacaagaacacccctgccgactcccagcccggccaaaaagacaaaacacatagacgcacacactcaggaggaaaagaaaaaaca | 0.104 | 0.424 |
| RT_008486.1 | NLRC5 NLR family, CARD domain containing 5 | WPPQDLWNPAK | NP_115582.4, | XP_00672 | 274 | TGAtggccccctcaagacctttggaatccagccaagtga | 0.086 | 0.065 |
| RT_015312.1 | NPL N-acetylneuraminate pyruvate lyase (dihydrodipicolinate syntha | EDCLSRRTLNY | NP_001186985.1, NP_001 | | 6156 | TGAgaagactgtctctcccgcaggaccctgaattattga<br>TAGggcgcctgccctctgctggctaggaccgggcccagcccagcccttcctgaagccagaagcgttgctgagtgtgttcc | 0.22 | 0.281 |
| RT_016124.1 | NPLOC4 NPL4 homolog, ubiquitin recognition factor | GACPLLARTGPSPALPEAR | NP_060391.2, | XP_01152 | 846 | ctgtaactgccccatagtgggcagccctggaggaacaaggggctggctgtcctgggctccctga<br>TAGagaggtgggggaggtgacagggagctaggcagccgtgctccctccagtgcggacttgtctccctctgagggaggcaa<br>gaggctggcgatcagggatcttgttgcattgggagcaggggcggctctcctggtccccaggagagatgctttgaggagca | 0.029 | -0.103 |
| RT_009589.1 | NUDT18 nudix (nucleoside diphosphate linked moiety X)-type motif 1 | APSSADLSPSEGGKR | NP_079091.3 | XP_01154 | 600 | ttcctctagattgcacaagggacagtgcctttaaccaagcgaggagtccaaagctcaggacctga | 0.128 | 0.215 |
| RT_003245.1 | NXN nucleoredoxin | GCMAFAKSLAFGEGSARVSWAE | NP_071908.2, N | XP_00525 | 391 | TAGcgtggctccggcctcctgagacgttatttaaaactcagccttctcctcctcccccctccttccttccgcccttggacttacccagcgtgccccgaatcccaccacccaagtgtccagcctctctgtggtgccttgtttctgcagtaaactcctcagccagcaccctggggtgtggaatcagcagcggcagagtc<br>TAAtgcattacagctgtgagaaagagcactgtggctttggcagctgttctacaggacagtgaggctatagccccttcaca | 0.25 | 0.203 |
| RT_016280.1 | OLFML1 olfactomedin-like 1 | CITAVR | NP_940876.2, | XP_00525 | 2996 | atatagtatccctctaatcacacacaggaagagtgtgtag<br>TGAgtggagacctgtgctccccggagaggggcagcagtgcgggaggggctttgcacagcagctcctgcaactgacccagt<br>ccgcaaatatttattgggggccagcccagggctgggactgggcatgaggtggtcaccaggattgagcttcctcagcaccc | -0.12 | -0.022 |
| RT_014428.1 | OLFML2A olfactomedin-like 2A | VETCAPRRGAAVR | NP_872293.2, N | XP_00671 | 974 | agtgggtaatacttgcttccacttgcagagcaccgtgccaagcacttcccacacacttacccgtttga<br>TAAattaaaatcgtactcataatcagctctgcatacatctgaagaacaaaaacatcaacgtcttttgtccagcctctttt | 0.315 | 0.257 |
| RT_010142.1 | OPA1 optic atrophy 1 (autosomal dominant) | IVLIISSAYI | NP_570850.2, N | | 1229 | tcttctgctgttccacctttctaa<br>TGAagggggatggcatctggtagggagggaggtatagttgggggacaggggtctgggtttggctccaggtgggaacaagg | 0.054 | 0.204 |
| RT_015055.1 | PAQR7 progestin and adipoQ receptor family member VII | WEQGLVK | NP_848509.1 | XP_01153 | 3864 | cctggtaaagttgtttgtgtctggcccacagtga | 0.138 | 0.12 |
| RT_005430.1 | PARP10 poly(ADP-ribose) polymerase family member 10 | APEGGVGDGGQQRPR | NP_116178.2, | XP_01151 | 3568 | TAAccgaaggggccaccctctggcctcctgcttcccaggctcccagctccgcacaggctgatgctccccgcccccaactgtggccgcctgagctgtccccggggacgcccctgcctccctctgcgggctccagaaggcggtgtgggggatggcggtcagcagcggccgaggggggccgggctaggtccc<br>TAGttaccgcggggaccacatcaagcttcaggccgacagcttcggcctccacatcgtccccgtctgaccccaccccggag<br>ggagggaggaccgacgcgacgcgatgcctcccggccaccgccccagcctcaccccatcccacgacccccgcaacccttca<br>catcaccccccctcgaaggtcggacaggacgggtggagccgtgggcgggaccctcaggcccgggcccgccgcccccagccc | 0.177 | 0.163 |
| RT_017560.1 | PAX2 paired box 2 | VGQDGWSRGR | NP_001291498. | | 7788 | cgcctgccgcccctccccgcctgcctggactgcgcggcgccgtga | 0.08 | 0.388 |
| RT_014634.1 | PCBP2 poly(rC) binding protein 2 | NNADSSIIPFCCSPPPMIHLCSF | NP_005007.2, NP_001122 | | 4813 | TAGaacaatgcagattcatccataatccctttctgctgttcaccaccacccatgatccatctgtgtagtttctga<br>TAGatagccggctttgccatccgggcatgtggccacactgcccaccaccgacgatgtgggtatggaacccccctctggata<br>cagaacccccttcttttccaaataaaaaaaaaatcatccagggcttggtgctttgtatcctagcttgtattcctttgaaaagcaacaggaggaaattggccaatatccagcactggggactgaggctttcctgtaaatgcaaggggttgggtggggagtgg | -0.054 | -0.075 |
| RT_026547.1 | PEBP4 phosphatidylethanolamine-binding protein 4 | IAGFAIR | NP_659399.2 | XP_01154 | 6085 | ggggaatag<br>TAAaagattttctgtagtgtttttgaaaagtttgcagcttatgtaatagcagataaaatttctaattgtaaaatgttaaa | 0.365 | 0.444 |
| RT_014747.1 | PHF14 PHD finger protein 14 | FAAYVIADKISNCKMLNCK | | XP_01151 | 10015 | ttgtaaaatctaa<br>TGAatacaaaataaaacaaataatttgtatttatggagtggaaaacagcttatttcaaggaaaaatatgactgccttatt | -0.055 | -0.122 |
| RT_013650.1 | PIAS2 protein inhibitor of activated STAT, 2 | NMTALLLLVPYCQVR | NP_775298.1 | XP_00672 | 3105 | attgctagtgccatattgtcaggtgcgaatggtgctcaaaagttgtatcattaaaatttga | 0.22 | 0.666 |
| RT_010003.1 | PICK1 protein interacting with PRKCA 1 | GLGGGRSPSPAPCGRSHCPR | NP_036539.1, N | XP_01152 | 2783 | TGAgtgccccgcggctgtggtgccgggggcagggtgcgtgggaggacggagcctgggagcggggcggggccgccgcgcaagggggcgacgcataaaggcctgctggcttggggcgcctgcctccctgctcctctgtcctcgcacagcgaacctgggctcctgcccaggacaggcaccagggtcatg | 0.28 | 0.257 |
| RT_023912.1 | PLAC1 placenta specific 1 | SSGLGSPEDAISR | NP_068568.1 | XP_01152 | 1924 | TGAtcctcaggtttggggtctcctgaagatgctatttctagaattagtatatag | 0.131 | 0.103 |
| RT_006100.1 | PLCB4 phospholipase C, beta 4 | THSQTSEHKLHG | NP_000924.3, | XP_00526 | 5813 | TGAagcagcaaacccacaaagcatcaaaagactcactcacaaacttctgaacacaaactccatggatga | 0.127 | 0.211 |
| RT_023238.1 | PLCZ1 phospholipase C, zeta 1 | QLMINDISLAMHRNK | NP_149114.2, | XP_00525 | 1169 | TAAcagctaatgataaatgacatatcattagctatgcatcgcaataaaacagccaaaatgaatgtctttatgttttctttttaaacataaagaaaaataattactcataa<br>TAAcatgatgaagaaactgacaggacagctctgtatttcatatttatacataaaggacttgaggagagaaaaaacacttt | 0.028 | 0.152 |
| RT_009701.1 | PLD5 phospholipase D family, member 5 | GLEERKNTLICLFF | NP_689879.2, N | XP_01154 | 2529 | aatatgtctcttttttag | 0.334 | 0.184 |
| RT_008256.1 | PMPCA peptidase (mitochondrial processing) alpha | RSEFPETAK | | XP_00526 | 2176 | TAAagagcactcaagttagtaggcgagaagggctcagtagaggactgctacattgtacgtggagtagcagtggctcggaatgccgctgtaccgttgctggctctcgggcaggagtctcagccccaggtttccggggcccccatgagtgaccttaggaggtctgaatttcctgaaactgccaagttgtatgt<br>TAGaggcctctgctaaagagcccggatgcagcgtcttccgtgggactgtccccaaggctgaggctcctgccaagtcctag<br>gggcctctgtacctgccctgctggaagccctgacttccccggggccccaggctgtgttagggttctctgggcctcttctt | 0.075 | 0.259 |
| RT_023570.1 | PNPLA7 patatin-like phospholipase domain containing 7 | SPDAASSVGLSPR | NP_689499.4, N | XP_00671 | 2907 | tgtaccagcagccctgcatacagggccctgtgagccccctgcagtcctgtga<br>TGAgaccaaaaagtttctcctactccagctggccctgtattcccctggaaggaagggaatcatgccttctatatacaga<br>cagattgccttcagaagagtggaagaaaatctccactatcaatgaacccagactcttgtcttcttcaagagcaagggcct | 0.063 | -0.102 |
| RT_009079.1 | POU2F3 POU class 2 homeobox 3 | ISTINEPR | NP_001231611. | XP_01154 | 4268 | ccggagatccaaactgtgattga | -0.045 | 0.055 |
| RT_020178.1 | PPP1R12B protein phosphatase 1, regulatory subunit 12B | LASSISPP | | XP_01150 | 97 | TGActtgctagtagcatctcacctccttga<br>TGAggatatattcaggggtccccaggaactcctcggtccagagaccaagtggtggccttggaaagcagatgtcaggcttt | 0.383 | 0.655 |
| RT_020123.1 | PPT2 palmitoyl-protein thioesterase 2 | GYIQGSPGTPR | NP_619731.2, N | | 11122 | ggtgtgcctgtga<br>TAAaggctatggtttattgcaggggattgggagggggggcgggaaaacatggagaattaagtcagataatgctggcagcca<br>aaggggcaaaatggcctgtgacattatcctgttcagagcttggagatgtacaagggacataggagcaatttacactgaca | 0.174 | 0.345 |
| RT_023177.1 | PRRC2C proline-rich coiled-coil 2C | SDNAGSQRGK | NP_055987.2 | XP_00524 | 19643 | cacagctgctgtaccagtga<br>TAGctcaggaccttggctgggcctggtcgtcatgtaggtcaggaccttggctggacctggaggccctgcccagccctgct | -0.016 | -0.058 |
| RT_027064.1 | PTP4A3 protein tyrosine phosphatase type IVA, member 3 | HLRALVSEERGAPR | NP_116000.1, N | XP_00525 | 8250 | ctgcccagcccagcaggggctccaggccttggctggccccacatcgccttttcctccccgacacctccgtgcacttgtgtccgaggagcgaggagcccctcgggccctgggtggcctctgggccctttctcctgtctccgccactccctctggcggcgctggccgtggctctgtctctctgaggtgggtcgggcg<br>TAAgaaagcaaaaagatctgggatatgtgttggaaaactgctttcccttatgttcactgtgccataatgctgctcgcagg | 0.19 | 0.491 |
| RT_009826.1 | PTPN4 protein tyrosine phosphatase, non-receptor type 4 | KNEELK | NP_002821.1, | XP_01150 | 10201 | aaatggcattttacaaaaaaaaaaatgaagaactcaaaaaaactttgaaaacttcagcactgttgcactttatgttttaa<br>TAGctcaatgggatggggaacctgccggagtccagaggctgctgtgaccaagcccccttttgtgtgaatggcagtaactg | 0.052 | 0.103 |
| RT_023418.1 | PTPRT protein tyrosine phosphatase, receptor type, T | LNGMGNLPESR | NP_008981.4, N | XP_01152 | 10957 | ggctcaggagctctga<br>TGAcctggacacagacagagacagagccaggcccggccctcccgcccccgacctgaccacgccggcctagggttccagactggttggacttgttcgtctggacgacactggagtggaacactgcctcccactttcttgggacttggagggaggtggaaca | 0.357 | 0.491 |
| RT_016814.1 | PVRL2 poliovirus receptor-related 2 (herpesvirus entry mediator B) | HWSGTLPPTFLGLGGRWNSTLDFS | NP_002847.1, | XP_01152 | 11516 | gcacactggacttctcccgtctctag<br>TGAcgggctttgtctgtggaaggagacagaaggaaattgacccactctcctatctccagacctttacctagcccctccag | 0.418 | 0.373 |
| RT_002300.1 | RASAL2 RAS protein activator like 2 | ALSVEGDR | NP_733793.2, N | XP_01150 | 7997 | gtttacagaatgttgctacttcacaatggcgatgtggtga<br>TGAcagcctagtgagggctgcaagaccatcctgcccggaccacagaaggagagttggcggtcacagagggctcctctgcc<br>aggcagtgggaagccctgggtttggcctcaggagctgggggtgcagtgggggactgccctagtccttgccaggtcgccag<br>caccctggagaagcatggggcgtagccagctcggaacttgccaggccccaaaggccacgactgcctgttggggacaggag | 0.261 | 0.394 |
| RT_014669.1 | RASSF7 Ras association (RalGDS/AF-6) domain family (N-terminal) m | SQLGTCQAPK | NP_003466.1 | XP_00525 | 869 | atgcatggacagtgtgctcaagctgtgggcatgtgcttgcctgcgggagaggtccttcactgtgtgtacacagcaagagcatgtgtgtgccacttcccctaccccaacgtgaaaacctcaataaactgcccgaagcagcttgagtgtgtgtggaggctgcgctgggcagggtctgaggtcttgaacttgagc | 0.289 | 0.33 |
| RT_013497.1 | RBM14 RNA binding motif protein 14 | APIMITCGRLR | NP_001185765.1, NP_001 | | 4196 | TAGccgagctctctgattaccgccgtttatcagagtcgcagctttcgttccgccgctcgccgacaaagtcctcgctggattaccgtcgcctgcccgatgcccattccgattacgcacgctattcgggctcctataatgattacctgcgggcggctcagatgcactctggctaccagcgccgcatgtagggccatc<br>TAAgccgccactgcaaaaaaaaacagggtcagaaaattacttgggtgggtagacttaggaattttctacttggtaaaagc | 0.152 | 0.119 |
| RT_024372.1 | RFPL3 ret finger protein-like 3 | AATAKKNR | NP_006595.1, N | | 7405 | attatacagtcataggagaaagatatgggacatttctataatctatattctaatttga | 0.076 | -0.112 |

| | | | | | | | | |
|---|---|---|---|---|---|---|---|---|
| RT_016562.1 | RGMA repulsive guidance molecule family member a | SPGVSTMAPQNAR | NP_064596.2, N | | 2448 | TAGacgcgtagatgtggagggaggcgcgggctccgtcctctcggcttccccatgtgtgggctgggaccgcccacggggtg<br>cagatctcctggcgtgtccaccatggccccgcagaacgccagggaccgcctgctgccaagggctcaggcacggacccctc<br>cccttctagtgcacgtgacaaggttgtggtga | -0.048 | -0.155 |
| RT_020305.1 | RGN regucalcin | GQVFFPAR | NP_690608.1, N | XP_00672 | 13379 | TGAggacaggtcttctttcctgccagagggagctctgaagacaactag | 0.238 | 0.345 |
| RT_020623.1 | RGS4 regulator of G-protein signaling 4 | FSPEGRGMKCQDSMLWK | NP_001095915.1, NP_005 | | 2129 | TAAttctcacctgaaggcagagggatgaaatgccaagactctatgctctggaaaacctga | 0.216 | 0.207 |
| RT_015540.1 | RILPL1 Rab interacting lysosomal protein-like 1 | LSDLRMDIEKLTPNSK | NP_847884.2, | XP_00671 | 15904 | TGAccttggcccatctccaccctccaacctggactgcccgcccaccagcgcctgcaaccgaactgcagcccaggggtcatt<br>gctgcctcaagcctctcggtgcagatgcaccctgaaaactgacccctcaaacagactgtctgatttgaggatggacattg<br>aaaaactgacgccaaactctaaagaaatgtttatttataccagggctatcactgtttctaa | 0.197 | 0.373 |
| RT_016579.1 | RNF183 ring finger protein 183 | EAVLTTELHSK | NP_659488.2 | XP_01151 | 367 | TGAgtgctgttcccagacaagaaaccaaaccttttttcggttgctgctgggtatggtgactacggagcctcatttggtattgtcttcctttgtagtgttgtttatttttacaatccagggattgttcaggccatgtgtttgcttctgggaacaattttaaaaaaaaaacaaaaaaacgaaaagcttgaaggactgggag | 0.144 | 0.257 |
| RT_016183.1 | RNF212 ring finger protein 212 | SFEVISK | | XP_01151 | 15579 | TGAagttttgaagtcatttcaaagacaaagttgtgtttttaccgcgacttccatgcctcctgcgggacctgctcaccttg<br>gggcagtgacacctga | 0.188 | 0.153 |
| RT_014053.1 | RNF43 ring finger protein 43 | SQNFGRPR | | XP_01152 | 14116 | TAAtcccagaactttgggaggcctaggtgggtggatcccttgaggccaggagttcaagaccaacctggccaacatggtga<br>aaccctatctctactaaaaatacaaaaattagccgggtatggtggcgggcacttataa | 0.056 | -0.075 |
| RT_014228.1 | ROBO4 roundabout guidance receptor 4 | TVSLRLPR | NP_001288017. | XP_00671 | 6043 | TGAaccgtgtccctgagacttcccagacgggaatcagaaccacttctcctgtcccacccacaagacctgggctgtggtgtg<br>tgggtcttggcctgtgtttctctgcagctggggtccaccttcccaagcctccagagagttctccctccacgattgtgaaa<br>acaaatga | 0.338 | 0.159 |
| RT_014154.1 | RPL13 ribosomal protein L13 | SPPGDLESVGSHAGSPR | NP_000968.2, N | | 3890 | TAAagccctcctggggacttggaatcagtcggcagtcatgctgggtctccacgtggtgtgtttcgtgggaacaactgggc<br>ctgggatggggcttcactgctgtgac | -0.067 | 0.033 |
| RT_002043.1 | RRAS2 related RAS viral (r-ras) oncogene homolog 2 | QMITMLALDEEAG | NP_001170785.1, NP_036 | | 6220 | TAGaatcccttcagttttagctaccaacggccaggaaaagccctcatcttctcttttctctcctcagtttacatcttgttggtacctttctagccttagacaaatgatcaccatgttagccttagacgaagaagctggctag | -0.013 | -0.134 |
| RT_016126.1 | SAPCD2 suppressor APC domain containing 2 | SLWAAWAPR | NP_848543.2 | XP_01151 | 1984 | TAGtccttgtgggccgcgtgggcccccagggccagcctggcactcagcccttcgaggtgggcgccccatcgcacccacc<br>ctctctggctggagaccccggcaggcccaggcacagtcccggagtgggcgccttcctgccgcccttgccagatgggctc<br>cccaggcctgcccccggctggtccccgcaccgagcgcttga | 0.021 | -0.119 |
| RT_024884.1 | SECISBP2L SECIS binding protein 2-like | LSISSCGR | NP_055516.2, N | | 5603 | TAActcaggaaatgtcggctctctatctccagctgtggaagggttgcagccattaccttttatgcttcatctcaacattt<br>tgcactgtccagtatttaa | 0.277 | 0.536 |
| RT_024986.1 | SEL1L2 sel-1 suppressor of lin-12-like 2 (C. elegans) | TLLPICHLGDLVPRYE | | XP_01152 | 593 | TAGgatgcggatcacaggaaaacctgctcacgggaatcagttcactccaaggtatccccactaaataaacagtttcccat<br>ccaaatcaagagtccgctgcagtttcccgctcagggctgtgatgggaaacagacaagacttggggaccatcctagagaaa<br>caacttcagacttcctgcttagtagagaaaaccctttgcgggatgctgggaactggtcacctaagaccttgttaccaatc<br>tgtcatcttggtgacctagtcccacgttatgaataa | 0.051 | -0.226 |
| RT_022798.1 | SERINC4 serine incorporator 4 | DTQGVISVWAEVNTMNWKK | NP_001244960. | | 5671 | TAAgtccttttcacaaactggggttcccctgacattgtactcctagagttggctcaaggggagctgtcagcccagctca<br>atacctcaaggacacacagggagttatctccgtttgggctgaagtcaatactatgaactggaagaagtggtcaaacacag<br>tctaa | 0.178 | 0.17 |
| RT_023546.1 | SETBP1 SET binding protein 1 | LHLGPRELTR | NP_056374.2 | XP_00525 | 8012 | TAGggcgggtctgggcgtctgcacctggggcctagggaactgacacgtgggaagcgcagtgagccggggcgggggcggaa<br>tcccccgctgcagggacacccacgcccttctctccagaagccgggcaggcagaatccggccagacgacggggctga | 0.015 | -0.103 |
| RT_006972.1 | SF1 splicing factor 1 | RGEQMVLGK | NP_973724.1, N | XP_01154 | 143 | TGAgttatttttgtggacggaatcggaacacgctggctccatatcgtgaaatttttattaattttttcttttttcctttgttacttctttatcttttcctttcttcagactccgtccaaggagatgctctccccggtcttctgctgcaatttagattcctttcccttctctccagttctccttcccttaccaaggagaggggagca | 0.323 | 0.289 |
| RT_014701.1 | SGOL2 shugoshin-like 2 (S. pombe) | ESKQKYSIKKMKCLMK | NP_689737.4, | XP_01150 | 268 | TGAagtgaatttatggattctggtttttctgaatttcaaagcataaggaatcaaaacagaaatatagtatcaagaagat<br>gaaatgcttaatgaaaaggtttttttttgtttctttggcctttcatggagtgttgatttgtccattcttaatgtttatt<br>aa | 0.044 | 0.153 |
| RT_006725.1 | SGSM1 small G protein signaling modulator 1 | ERGPGVCSQAWSSVRK | NP_001035037. | | 5791 | TGAggggcacctcaccccggcagcctcagccaagctgcccctgccccgctcctctgcttacttttcctcctggctggatg<br>ggcaccccgggagcggggtcctggtgtctgttcacaagcgtggagttcagtgcgcaaagaaactaccctgacttttactt<br>ctgggcagatggggtggagggagtacccctcaattcagccttacattttcctgtttga | 0.073 | 0.028 |
| RT_008135.1 | SH3RF2 SH3 domain containing ring finger 2 | DTNSLASK | | XP_00526 | 13815 | je ne trouve pas le peptide | | |
| RT_017390.1 | SLC38A8 solute carrier family 38, member 8 | WAASAGQEGALR | NP_001073911. | XP_01152 | 918 | TGAtgggcagctagtgccgggcaggaaggggccctccgggggctgaccctacgtggctgctgtatgcagccaggagaccg<br>atgccatttcttttcctcataa | 0.088 | 0.065 |
| RT_013448.1 | SLC9A3R2 solute carrier family 9, subfamily A (NHE3, cation proton a | TLGLSLPR | NP_001123484. | XP_00525 | 1085 | TGAgccccttcctgcctgtctcgggaccctgggaccccctcccgcacggaccttgggcctcagcctgccccgagctccccc<br>agcctcagtggactggagggtggtcctgccattgcccagaaatcagccccagccccggtga | 0.091 | 0.066 |
| RT_018371.1 | SLC9A8 solute carrier family 9, subfamily A (NHE8, cation proton ant | GLAEMRASSSPFKT | NP_001247420. | XP_01152 | 5222 | TGAcgccaggtgccaaggcttcaggcaggcaggcccaggatgggcgtttgctgcgcacagacactcagcaggggcctcgc<br>agagatgcgtgcatccagcagccccttcaagacataa | 0.189 | 0.485 |
| RT_018499.1 | SMOC2 SPARC related modular calcium binding 2 | AHTPKAVPR | NP_071421.1, NP_00115 | | 12481 | TAAatggctcatacccgaaggcagttcctagacacatgggaaatttccctcaccaaagagcaattaagaaaacaaaaacagaaacacatagtatttgcactttgtactttaaatgtaaattcactttgtag | 0.184 | 0.172 |
| RT_018427.1 | SNX29 sorting nexin 29 | RHPHGPGCTGELASRGVITHR | | XP_01152 | 6366 | TAGcggcatccccatggcccagggtgcacgggggaattagcctctcgcggagtcatcacgcatcgactgaattccctggt<br>gaaaactgagttagccagttgttcctaa | 0.251 | 0.171 |
| RT_021657.1 | SPACA7 sperm acrosome associated 7 | SLLEPPPTSLRNR | NP_660291.2, | XP_01153 | 13958 | TGAggccgcagccccagaccccctgcgcaggagaggaggcctgctagaacccccacccaccagcctccggaacagggcacttgtgtgcacacgcccacgttctctgaaccattccacataa | 0.012 | 0.119 |
| RT_009922.1 | SPAG11B sperm associated antigen 11B | AGMLKYPK | NP_478108.2, N | XP_00527 | 335 | TAAaatataagctcccggaaggcaggatgttgaagtatcccaaggcttaaaggaatgtgtggcttatagtaggtgttc<br>aataaatatttgttga | -0.046 | -0.089 |
| RT_005096.1 | SPAM1 sperm adhesion molecule 1 (PH-20 hyaluronidase, zona pellu | LAEMNNMSILK | NP_001167515. | XP_01151 | 3094 | TAAttgcgcaggttagctgaaatgaacaatatgtccatcttaaagtgtgctttttcgactaattaa | 0.092 | 0.116 |
| RT_002817.1 | SPATA21 spermatogenesis associated 21 | QELLLGFAFVAW | NP_940948.1, | XP_01153 | 4868 | TGAccctctggaggcctgactctaggcagctgctcaaggcctgtagacagatgggccaccaggcaggagctgctgctggg<br>atttgcttttgtggcctggtga | 0.219 | 0.432 |
| RT_006931.1 | SPDYE3 speedy/RINGO cell cycle regulator family member E3 | HLYSQLLK | | XP_01151 | 2077 | je ne trouve pas le peptide | | |
| RT_019025.1 | SPOCK3 sparc/osteonectin, cwcv and kazal-like domains proteoglyca | LMTVEINK | NP_001238896. | XP_01153 | 588 | TGAttgatgacagttgaaatcaataaattctacatttctaatatttacaaaaatgatagcctatttaaaattatcttctt<br>ccccaataacaaaatgattctaaacctcacatatattttgtataattatttgaaaaattgcagctaa | 0.272 | 0.294 |
| RT_004780.1 | SPTB spectrin, beta, erythrocytic | QVGLAGAAESGRAETGQPRPWPPL | NP_001020029. | XP_00526 | 11484 | TAGcaggtggggctggcgggcgcggcggagtcgggacgcgcagagaccgggcagcccaggccttggcctcctctcgccgc<br>cgcccgccggtcgaatcgagatgagtcgagtcgcgggccgggcccccgggccccacgctgcactgcaaaagctgccttcg<br>cccgcgtgtcctgggccgacgcccccctccatgccctcgcccgcgctggctccgctccagggacaggggctcccagctcct<br>ctggcgcctttggggcctccaggcctgggggcaggaaacggttctctgaggcacacgccctcctctcccaccctcctcct<br>gagccgagggggcacaggcccccttacccccaaccaaaaaaaaaaaaaaaaaaagagctggagaaggctgatggggttcttc<br>aaacgctgcccgttcctaaagcaagtcttgcttcgggtcacctcccacctggtggcagccagggaaaggggaaaggaaga<br>agacactggaaatgcatggccagcccccctaggggcatga | 0.137 | 0.107 |
| RT_026612.1 | STIL SCL/TAL1 interrupting locus | PFNSLPF | NP_003026.2, N | XP_01154 | 11171 | TAAccttttaactccctgcccttttaa | 0.112 | 0.195 |
| RT_009155.1 | SUPT7L SPT7-like STAGA complex gamma subunit | EMFCPDLLDPTEK | NP_001269661. | XP_00526 | 6091 | TAAaaggaaaagagggagatgttttgtccagacctactagacccaacagaaaaggtttttgtattagaatctgtttccttaaaaattgatttgactcctgttcttaaacacaagtggtttttcctaattccagaggaactggacgtcaccaaacaaggttgcattttactttgcatccagtttttatagttttccaca | 0.058 | -0.122 |
| RT_010002.1 | SWI5 SWI5 homologous recombination repair protein | AGSSPLVHSSQGQK | NP_001035100. | XP_00525 | 5056 | TGAgcaggctcatcgcccctgtccacagctcccagggacagaagggtgtggacatgataaacactgagagcccaaccag<br>cacacctacagagtttccagcgagacaatgccagaagcacttttctagagagcccaagcccaatcaggaggaggctagag<br>aagaggcgggcagggaggaaggggcgttcccagctccaaggttcctgaaatgtcgcagtgatacttgtgtaggggtacat<br>gtactttatttgtcaataaacgtatctgtacactga | 0.325 | 0.292 |
| RT_002713.1 | SYCE2 synaptonemal complex central element protein 2 | CAGHCFRSTDQQR | | XP_01152 | 5976 | TGAcaccttggggtcagatgtgcaggccactgcttcagaagtacagaccaacagagatggtga | 0.325 | 0.421 |
| RT_026555.1 | SYT15 synaptotagmin XV | PFAQHRGSALGADHPR | NP_114118.2, | XP_01153 | 12022 | TGAccctttcgcccagcaccgcggttccgctttgggagccgaccatcctcgcagttgcatgtga | 0.336 | 0.432 |
| RT_026388.1 | T T brachyury transcription factor | AQVPKDAVTFCRGSQW | NP_001257413. | XP_01153 | 4595 | TGAagcagcaaggcccaggtcccgaaagatgcagtgactttttgtcgtggcagccagtggtga | 0.237 | 0.211 |
| RT_019228.1 | TCTE3 t-complex-associated-testis-expressed 3 | MVWLRLR | | XP_00671 | 20255 | TAGctcattacagattcagacaagaagtgcggcggaagcttacacctgcacgtggtgggacagaggcaagaacggatggt<br>ctggctgaggctgagagaaccaagctggattaggatcggcttgagggcccggcgtcgtcagtggtga | 0.379 | 0.491 |
| RT_007777.1 | TET3 tet methylcytosine dioxygenase 3 | AVSVVLLPSYLGAGWGCR | NP_001274420. | XP_01153 | 13737 | TAGgtgcccagggagccagcgtacctcagcgtcgggcctggcccgagctgtctctgtggtgcttttgccctcatacctgggggcgggttgggggtgcagaagtctttttatctctatatacatatatagatgcgcatatcatatatatgtatttatggtccaaacctcagaactgacccgcccctcccttacccca | 0.008 | 0.035 |
| RT_026131.1 | TEX26 testis expressed 26 | IVQMKK | NP_689538.1, | XP_01153 | 1363 | TGAaaagggaaaatagtacaaatgaagaaaatctga | 0.169 | 0.373 |
| RT_013769.1 | THUMPD2 THUMP domain containing 2 | TVTQMNLELK | | XP_00671 | 7176 | not found | | |
| RT_001050.1 | TMEM106B transmembrane protein 106B | KLEEMDLK | NP_060844.2, N | XP_00524 | 1964 | TAAaaactggaagagatggatttaaagaagaaatatctattgatatttcctatactctcaatgaagaggtatttcctaat<br>aggagaccttaaattgaacaaacctaaagtttacacttctaagagtacagttaaaagtatgtggacctgcagttcttgta<br>actctccactctgtgttaatgatatatttgtactaggatcttttacttga | 0.046 | 0.195 |
| RT_012578.1 | TMEM155 transmembrane protein 155 | RMTSHSHLQNLK | | XP_01152 | 121 | not found | | |
| RT_017166.1 | TMEM185A transmembrane protein 185A | WTGTHSLR | NP_001167563. | | 1538 | TAGatgccacttccggggacagagcttaagtggactgggacgcactctctccgccttcctctgccccctcgttcaccccg<br>cagaccagaaccagtactggagctgggtctccaggtacgtccatctcatgccttgtttgcatccagcgcctatcagccac<br>tcaccacgacgggacgcggaagtggcaggtga | 0.008 | 0.127 |
| RT_019558.1 | TMEM54 transmembrane protein 54 | TLQATLLCDQARIPGAGLR | NP_277039.1 | XP_00527 | 13000 | TGAgagatgatgtcctgcccaggcccgatggccactaggaccctgcaagcaactctgctctgtgaccaggccaggattcc<br>tggagctggcctgagagggctcaatggaccctcggggacccaagtggggctttcaaccctctcccccaccacccagccca<br>ctgcactgaaatga | 0.14 | -0.036 |
| RT_022387.1 | TMEM92 transmembrane protein 92 | VTSCLESCHQQPPPQCLLDQAR | NP_001161687. | XP_00672 | 5318 | TGAgtcacctcctgcctggaatcttgccatcagcaacctcctccccagtgcctcctggatcaagctagagactgctggca<br>ccccaggaatgtccctgcccatcctgccgtgtctctgttcattcttggatttaacttattactttttctgcttctgtttc<br>caccccagctgcctctcttgtcctga | 0.304 | 0.348 |
| RT_004139.1 | TNFRSF25 tumor necrosis factor receptor superfamily, member 25 | RSGGPCRSPK | NP_683871.1, NP_683868 | | 10644 | TGAcacggcgcccacttgccacctaggcgctctggtggcccttgcagaagccctaagtacggttacttatgcgtgtag | 0.343 | 0.421 |
| RT_024642.1 | TNFSF14 tumor necrosis factor (ligand) superfamily, member 14 | GALDMGLTRGELR | NP_742011.2, N | XP_00525 | 5902 | TGAaggaaggagcgtggtgcattggacatgggtctgacacgtggagaactcagagggtgcctcaggggaaagaaaactca<br>cgaagcagaggctgggcgtggtggctctcgcctgtaa | 0.123 | 0.12 |
| RT_019102.1 | TNK2 tyrosine kinase, non-receptor, 2 | DASGEPEGLPEGIT | NP_005772.3, N | XP_00526 | 7353 | TGAgatgcgtctggagagccagagggcctgctgaaggaatcacctga | 0.092 | -0.004 |
| RT_005492.1 | TOX4 TOX high mobility group box family member 4 | SYLLGKCPR | NP_055643.1, NP_00129 | | 1913 | TAGtccttcctgttctccaagccagtgaagagttatctgctgggaaagtgtccaagagcctgtttttga | -0.102 | -0.119 |
| RT_015033.1 | TPCN1 two pore segment channel 1 | APEGPSQGDPSPGSSR | NP_001137291. | XP_01153 | 8686 | TAGcccagcgcccgaaagccgtctcttctatgcaataacacaatagtattactctactgcgatgtacggaactgcggtgtgtgtacacatactcacgtatatgcacatatttatatacaggaagaaaaaaagacagacaagatggggcttggtttataaccaccttgccctgtcttccttaactccagaagcca | 0.143 | 0.21 |
| RT_019778.1 | TPCN2 two pore segment channel 2 | LAYTGAR | NP_620714.2, | XP_00527 | 9965 | TGAcgtccgggctgccgtcccagcaggggcggcaggagagagaggctggcctacacaggtgcccgtcatggaagaggcggccatgctgtggccagccaggcaggaagagacctttcctctgacggaccactaagctggggacaggaaccaagtcctttgc<br>gtgtggcccaacaaccatctacagaacagctgctggtgcttcagggaggcgccgtgccctccgctttcttttatagctgcttcagtgagaattccctcgtcgactccacagggacctttcagacaaaaatgcaagaagcagcggcctcccctgtcccctgcagcttccgtggtgcctttgctgccggcagccctt | 0.186 | 0.426 |
| RT_002189.1 | TPM1 tropomyosin 1 (alpha) | PWFSLLASCLRARHTLCFLLYR | | XP_00525 | 2616 | TGAccctggttctctctcttagcatcctgccttagagccaggcacacactgtgctttctattgtacagaagctcttcgtt<br>tcagtgtcaaataaacactgtgtaa | 0.262 | 0.432 |
| RT_024775.1 | TPM3 tropomyosin 3 | LSPFLLCSGSAPFTPR | NP_689476.2, | XP_00671 | 2836 | TAAttatcaccgtttctgctctgttctggatctgcccccttactcctcggggaacccaaggccccactctcgctctggattccatttgggtcagcctggctggtccccaaggcattaggatgggggagcaaaaagcaacttatgtattttcttccaccccccaccccaaattaa | 0.264 | 0.433 |
| RT_000591.1 | TPRA1 transmembrane protein, adipocyte asscociated 1 (lipid metab | EEASRPRVPRGGGPGQGTFCGQ | NP_001136118. | XP_01151 | 66 | TGAcatcgcttccatgccctgccacactggcagcatcaacagcacagacagcgagcgctggaaggccatcaatgcctgag<br>ggcagctgccagggcctgtggaggacaggccagagaggaggccagcaggcccagagtcccaggggaggaggaccaggtcaagggacgttctgtgggcagtagc | 0.262 | 0.362 |
| RT_020019.1 | TRIM68 tripartite motif containing 68 | GLELGLAPMGLGGPSH | NP_001291425.1, NP_060 | | 6601 | TAAgaaagctaccaccctaaccagagaggcttggaattggcctggcccccatggggcttggaggaccgagccactga | 0.15 | 0.103 |
| RT_006402.1 | TRIM7 tripartite motif containing 7 | VTVANRAGEQGLWTPEGVPFLRSH | NP_976038.1, NP_976039 | | 3507 | TGAggggcactgctggggagctcctgtctctgggctgccggtgggaggggatgtcgccctccccagagatgcctggtccgtctgggtctgccctccgtgctcctgaccctgctgcccaagagagcctgctacagacacaaccccgaggcaggagagtgactgtggccaaccgagcaggggaacaggg | 0.102 | 0.028 |
| RT_014420.1 | TSPEAR thrombospondin-type laminin G domain and EAR repeats | GRGCPAATGVAR | NP_659428.2, N | | 422 | TGAggccgcggctgcccggcagcaactggggtggccaggtggggcgggacccagcacctcccccaactgaccctggacg<br>tccaggtgggcccagcctagggcatcccgggcctgggcatcaggcagggtgcaggtggggccatctgcggccctggttcg<br>ggcagggacatccatccacctctgagtccttctga | 0.073 | 0.119 |
| RT_013052.1 | TSPO translocator protein (18kDa) | VPGPPGTAAAPAGAITLVMWWPSF | NP_000705.2, N | | 1818 | TGAgtgcccggcccaccagggactgcagctgcaccagcaggtgccatcacgcttgtgatgtggtggccgtcacgctttca<br>tgaccactgggcctgctag | 0.183 | 0.257 |
| RT_008971.1 | TSTA3 tissue specific transplantation antigen P35B | TGSGASGPSAGR | NP_003304.1, | XP_01151 | 9638 | TGAagctggaagacaggatcaggtgccagcggaccatcggctggcagagcccagcggccaccacccgtcaaccctgccaggagctga | -0.078 | 0.211 |
| RT_021416.1 | TTC13 tetratricopeptide repeat domain 13 | LFFKTWNYK | NP_001116307. | XP_00671 | 3290 | TAGttctgctgctgtatttatacaagtaaagggccggaccttcttgcttcttaagttatttttttaaaacatggaattataaagaaattaggtcctctattacttttgataccaatttttataggaattga | 0.125 | 0.162 |
| RT_021425.1 | TTC39B tetratricopeptide repeat domain 39B | WKNDLVNK | NP_001161813. | XP_01151 | 4525 | TGAtcagaacttgaaggatggaatttttcctcaattagcatcggcatctgctatagattcaaccttatattaaagtggaaaaatgatcttgtgaataagagacaaaaaaacagttttttggtcaatattttctatcatctgagagatttactgttggtctacataatttttttcctatggaatggcattctatattatc | 0.324 | 0.329 |
| RT_003336.1 | TTC7A tetratricopeptide repeat domain 7A | QQGTWVR; REGGAGQR | NP_001275882. | XP_01153 | 17188 | TGAcgacgctgcagccgcagggagggaggggctggccagaggggagaggcagcagggaacgtgggtcagggtggggcaacagtggcatcaggtgcggggcctcagggaaatacatctttag | 0.4 | 0.71 |
| RT_022743.1 | TUBGCP2 tubulin, gamma complex associated protein 2 | KGVGSAGTGANGSR | NP_001243547. | | 6900 | TGAgccctggctgtgacaggaaggaagggtgtggggtcagcagggactggtgcaaatgggtccagaattttcaaatcgaa<br>tgctctgtgttctgtctttgcagtttaa | 0.052 | 0.066 |
| RT_023684.1 | TUFT1 tuftelin 1 | AISIFLQ | NP_064512.1, N | XP_01150 | 16246 | TGAgctgcctggagatggttgctgccattgctgctgcctctgcctcggagaagcccactgccccttgttggctgttaacac<br>tgcctttgacttcctgactgtccctggctgcacccaggacttcgggctcctgtgtctcaccattcccaagcccctggcc<br>actctaagctgggcagcaggagcacgagcaccattcaaggcactgcagcccctttggaagacattgtcctgcaagcagga<br>gccagggcaatatctatattcctacagtga | 0.196 | 0.008 |

| | | | | | | | | |
|---|---|---|---|---|---|---|---|---|
| RT_007176.1 | UBA2 ubiquitin-like modifier activating enzyme 2 | TEMPLNRTLLLFSLSGQNQIVMSFV | NP_005490.1, | XP_00525 | 2127 | TGAacagaaatgcctctaaacagaaccctcttactatttagtttatctgggcagaaccagattgttatgtcctttgttcc aaagggaaaaaattgacagcagtga | 0.25 | 0.384 |
| RT_009758.1 | UBL5 ubiquitin-like 5 | RILIFLPR | NP_001041706.1, NP_077 | | 9141 | TAGatgagaatcctcatcttcctgccccgctttcctctcccatcctcatcccccacactgggatag | 0.097 | 0.127 |
| RT_022085.1 | uncharacterized LOC105371419 | APPGHSDRGRGVR | | XP_01152 | 4152 | TAAcagccacacccgaccagtgacagcagggccctggccagggctcggggggtcacctggggcaagggacctccagagaggaatctgagggcccctcctgggcacagtgacagagggcggggtgtgaggaagccgactggaagagccgctgttcacacaa attcccagagggaagggcagggaaggacagcgtag | 0.172 | 0.152 |
| RT_000368.1 | UNG uracil DNA glycosylase | SSAEGWPLR | NP_550433.1, NP_003353 | | 4500 | TGAtcatcagctgaggggtggcctttgagaagctgctgttaa | 0.157 | 0.329 |
| RT_023844.1 | UVSSA UV-stimulated scaffold protein A | AGPSALAISTFSLCQCLR | NP_065945.2, | XP_01151 | 5089 | TAGagagcggggcccagtgcactggccatcagcactttctccctctgccagtgtctcaggacagcagagtgggcgtgggt ctgggcagtaaccatgctttgtctattactgtgtttgatgtaaa | 0.028 | 0.215 |
| RT_012400.1 | VASP vasodilator-stimulated phosphoprotein | TTQECIVPTPHPTWK | NP_003361.1, | XP_00525 | 921 | TGAccacagggacccagaagacccgcttctcctttccgcacacccggcctgtcaccctgctttccctgcctctacttgacttggaattggctgaagactacacaggaatgcatcgttcccactccccatcccacttggaaaactccaagggggtgtggcttccctgctcacacccacactggctgctga | 0.494 | 0.658 |
| RT_001270.1 | VDAC3 voltage-dependent anion channel 3 | ASDLSLEVKRNEPTMFWP | NP_001129166. | XP_00671 | 4076 | TAAtgtggtttgaggaaagcatcagatttgtccctggaagtgaagagaaatgaacccactatgttttggccttaa | 0.149 | -0.081 |
| RT_022463.1 | VMO1 vitelline membrane outer layer 1 homolog (chicken) | AAVVSRR | NP_001138413.1, NP_001 | | 21723 | TGAgccgctgtggtgtcgcggcggcgcctacctagtggctttctcgcttcgcgtggaggcacccacgaccctcggtga | 0.064 | 0.066 |
| RT_017770.1 | VPREB1 pre-B lymphocyte 1 | SAEEAEPLSISWSCSVAVNPHWR | NP_009059.1, NP_00129( | | 4587 | TGAactgaagacagcagaggcacgcatccccttggagagactgtcatggaagagggtggagtcgccgcccgaagcgccgaggaggctgagccactcagcatctcctggtcctgcagtgttgctgtaaatccccattggagactgcattag | 0.129 | 0.1 |
| RT_013079.1 | WDFY3 WD repeat and FYVE domain containing 3 | FNKLSGDHGL | NP_055806.2, | XP_01153 | 7738 | TGAagattcaacaagctgagtggagaccatggtctgtag | 0.237 | 0.437 |
| RT_019157.1 | WDR19 WD repeat domain 19 | ADTMLLR | NP_079408.3, | XP_01151 | 8174 | TGAttggcacgtgcagatacaatgctcctgagaagacagcattttccacaggaggctgtttcctcccctggtggatttaa gagacggtcctttctggatacagagaaatga | 0.269 | 0.294 |
| RT_004152.1 | WNT7A wingless-type MMTV integration site family, member 7A | MLSLSFLLR | NP_004616.2 | XP_01153 | 7432 | TGAgccccgtgtgcacaccaccctcccgctgcaagtcagattgctgggaggactggaccgtttccaagctgcgggctccc tggcaggatgctgagcttgtcttttctgctgaggagggtacttttcctgggtttcctgcaggcatccgtgggggaaaaaa aatctctcagagccctcaactattctgttccacacccaatgctgctccaccctcccccagacacagcccaggtccctccg cggctggagcgaagccttctgcagcaggaactctggacccctgggcctcatcacagcaatatttaac | -0.001 | 0.066 |
| RT_022489.1 | ZCCHC5 zinc finger, CCHC domain containing 5 | GGPHVGQSTNMRPPLFQYPCLIHY( | NP_689907.1 | XP_01152 | 9291 | TAAgggggaccccacgtgggccaatctacaaatatgcgtccccctctcttccaatatccatgtctcatacactatggcttactgttgaaatccaactgggaagaagataattctttgagcaagcaatggtag | 0.007 | -0.15 |
| RT_023327.1 | ZNF140 zinc finger protein 140 | NYECMEFFKK | NP_001287707. | XP_01153 | 894 | TGAatttacactgcaaagaaaaactatgaatgtatggaattttttaaaaagaagtataatgccttacttcagagaactct tggaaagaagccttatgtgaaagtgatgactgtgaagtaa | 0.218 | 0.382 |
| RT_027047.1 | ZNF282 zinc finger protein 282 | GAERPRAPPSHLLECPGTTGPQMV( | NP_003566.1, | XP_00671 | 5072 | TAGggctgggctgggggagggcagggccggacggagtggatcgggggcggcctgagcaccaaccaccttgccgggtgtcctcagccaccgtctggaaatcggcaacaggcattgcactccggttgggggtcccccagggtggggcagggatcccccagat ctgtctggtctgaatggacgcccagctcatctagggtggacccagctgctggggaagagccagggggaccgcgaggagcc gagcgtcctcgggcaccgccctcacacctcctcgagtgccctgggaccactgggccacagatggtcatcaggggaagcca ccagggagtcccgaagcccttctga | -0.116 | -0.103 |
| RT_014705.1 | ZNF283 zinc finger protein 283 | VVKEYFVCVYRQLIIIRTLTLEK | NP_001284681. | XP_00525 | 4338 | TGAttgaaagttgtaaaagaatattttgtgtgtgtgtatagacaacttatcataataagaactcttactcttgagaaaccttgtgaatgtaagggttgtgcaaaagccattcatttctgtttatgggcaattatcttgctatccagcaattcatactagtga | 0.227 | 0.294 |
| RT_010661.1 | ZNF34 zinc finger protein 34 | FIAADLR | NP_001273699.1, NP_085 | | 7548 | TAAcagtggcgcggtgtccgagggcagagtccagctgagcacttcctgcatgcgcccccggcacctgactctgccctttatgtattatccacacgatgttttcacagagtgaaaggacgtttctcattaaacaaacctcttttcttaaatcaaaagcagtgcatgttcattttagagaaattgagacagaaaag | 0.201 | 0.152 |
| RT_002045.1 | ZNF410 zinc finger protein 410 (glycosylation candidate) | ATLSDLKMR | NP_067011.1, NP_00122S | | 5300 | TGAgcgtgggtgctgactcctggaagagcaactctatctgatctcaaaatgcgtatactgggaacaggatgccttagcccacaacagaaccagaatgaatctttgaaggcacaagactctgcttttgccactcttcctctttcctggtatagaagatggatgtag | -0.098 | -0.025 |
| RT_023991.1 | ZNF512B zinc finger protein 512B | VALPLLR | NP_065764.1, | XP_00526 | 3859 | TGAgcatggtgcgtggggcctggcccccatgcagccgccacaccgccctctgttcaaggcagggccagctccgggacctc tgtcctccagttctccatcccccacccccgcctcttcatccgtccagtggggcagccagccactctcccttccctgaaggtggcccttcccctgttaaggctgccacagggcacacctgggttggggctccccaaggtgtgcccagcagaagtgcaatag | 0.074 | 0.292 |
| RT_010258.1 | ZNF580 zinc finger protein 580 | RGVGAPAPGGGTSALGSR | | XP_01152 | 1229 | not found | | |
| RT_005360.1 | ZNF584 zinc finger protein 584 | RATESSSTRTLIVR | | XP_00525 | 8735 | TAAagagctacagagtcatccagcaccaggacactcatagtgaggggaaaccaagaaggcacactgagcatggggcagct ttcccacctggttccagttgtgggcaacagcaagaagtccatgtggcagagaagctgttcaaatgcagtgactgtgggaa ggtgttcttaa | 0.176 | 0.259 |

SUPPLEMENTARY TABLE 4

*Relative Fluc fold change*

**UGA Fig 1F**

| Condition | N1 | | N2 | | N3 | |
|---|---|---|---|---|---|---|
| | Norm Fluc/Rluc | SD | Norm Fluc/Rluc | SD | Norm Fluc/Rluc | SD |
| Vector | 1.0000 | 0.2409 | 1 | 0.2587 | 1 | 0.3428 |
| H-Ras V12-D5 | 1.0940 | 0.3269 | 0.5199 | 0.2041 | 0.9202 | 0.1283 |
| H-Ras V12-D12 | 0.5663 | 0.1330 | 0.3747 | 0.1497 | 0.4701 | 0.0662 |
| H-Ras V12-D20 | 0.4043 | 0.0241 | 0.2683 | 0.1672 | 0.3423 | 0.1405 |

**AQP4 Fig 1F**

| Condition | N1 | | N2 | | N3 | |
|---|---|---|---|---|---|---|
| | Norm Fluc/Rluc | SD | Norm Fluc/Rluc | SD | Norm Fluc/Rluc | SD |
| Vector | 1.0000 | 0.1645 | 1 | 0.1701 | 1 | 0.1370 |
| H-Ras V12-D5 | 1.7850 | 0.6203 | 0.8558 | 0.1782 | 1.828 | 0.3805 |
| H-Ras V12-D12 | 0.7813 | 0.0213 | 0.3300 | 0.0806 | 0.7042 | 0.1719 |
| H-Ras V12-D20 | 0.3594 | 0.0807 | 0.2036 | 0.0492 | 0.4357 | 0.1050 |

**UGA Fig 2A**

| Condition | N1 | | N2 | | N3 | |
|---|---|---|---|---|---|---|
| | Norm Fluc/Rluc | SD | Norm Fluc/Rluc | SD | Norm Fluc/Rluc | SD |
| Vector | 1.0000 | 0.1771 | 1 | 0.1162 | 1.0000 | 0.0238 |
| H-Ras V12 | 0.4557 | 0.1307 | 0.2343 | 0.1541 | 0.6687 | 0.1065 |
| Vector Gen | 1.9220 | 0.5004 | 11.3100 | 1.07700 | 5.5810 | 0.0345 |
| H-Ras V12 Gen | 1.1860 | 0.1066 | 7.9870 | 1.61800 | 4.3510 | 0.6618 |

**AQP4 Fig 2A**

| Condition | N1 | | N2 | | N3 | |
|---|---|---|---|---|---|---|
| | Norm Fluc/Rluc | SD | Norm Fluc/Rluc | SD | Norm Fluc/Rluc | SD |
| Vector | 1.0000 | 0.0864 | 1 | 0.2858 | 1.0950 | 0.1338 |
| H-Ras V12 | 0.3274 | 0.1348 | 0.5268 | 0.0705 | 0.2753 | 0.0467 |
| Vector Gen | 2.4020 | 0.4240 | 1.5830 | 0.0189 | 2.6290 | 0.6564 |
| H-Ras V12 Gen | 0.9056 | 0.0795 | 1.4380 | 0.3512 | 0.9912 | 0.0870 |

**MMuLV Fig2A**

| Condition | N1 | | N2 | | N3 | |
|---|---|---|---|---|---|---|
| | Norm Fluc/Rluc | SD | Norm Fluc/Rluc | SD | Norm Fluc/Rluc | SD |
| Vector | 0.7927 | 0.07473 | 1.0000 | 0.1428 | 1.1450 | 0.2049 |
| H-Ras V12 | 1 | 0.00017 | 1.3520 | 0.000165 | 1.3150 | 0.147 |
| Vector Gen | 0.7581 | 0.07355 | 1.0250 | 0.07029 | 1.1050 | 0.3146 |
| H-Ras V12 Gen | 1.046 | 0.1354 | 1.3480 | 0.09148 | 2.0100 | 0.0386 |

**UGA Fig 2B**

| Condition | N1 | | N2 | | N3 | |
|---|---|---|---|---|---|---|
| | Norm Fluc/Rluc | SD | Norm Fluc/Rluc | SD | Norm Fluc/Rluc | SD |
| Vector | 1.1310 | 0.1031 | 0.8210 | 0.0169 | 1.0000 | 0.1838 |
| PML-ER | 1.0000 | 0.2771 | 1.0000 | 0.1238 | 0.9085 | 0.2310 |
| Vector 4OHT | 1.1430 | 0.1341 | 0.8441 | 0.1567 | 0.9875 | 0.0128 |
| PML-ER 4OHT | 0.0657 | 0.1138 | 0.0000 | 0.0000 | 0.9542 | 0.3150 |
| Vector Gen | 15.9600 | 4.3580 | 17.1100 | 3.2800 | 17.9200 | 2.8720 |
| PML-ER Gen | 11.0100 | 1.1130 | 17.9700 | 3.8380 | 16.8300 | 2.5950 |
| Vector 4OHT Gen | 14.6700 | 1.6520 | 19.1300 | 4.4710 | 11.5500 | 0.3844 |
| PML-ER 4OHT Gen | 2.2040 | 1.2050 | 3.9580 | 0.6262 | 6.0300 | 0.6035 |

**AQP4 Fig 2C**

| Condition | N1 | | N2 | |
|---|---|---|---|---|
| | Norm Fluc/Rluc | SD | Norm Fluc/Rluc | SD |
| Vector | 1.0000 | 0.0554 | 1.0000 | 0.1457 |
| PML-ER | 0.8622 | 0.0583 | 0.9104 | 0.1158 |
| Vector 4OHT | 1.1640 | 0.2160 | 1.0570 | 0.1785 |
| PML-ER 4OHT | 0.6573 | 0.0927 | 0.7367 | 0.1292 |
| Vector Gen | 3.7630 | 0.7502 | 2.5350 | 0.2966 |
| PML-ER Gen | 2.9520 | 0.3866 | 2.9780 | 0.2403 |
| Vector 4OHT Gen | 2.7590 | 0.4541 | 2.4750 | 0.1161 |
| PML-ER 4OHT Gen | 1.8230 | 0.1799 | 2.0780 | 0.6261 |

**MMuLV Fig 2D**

| Condition | N1 | | N2 | |
|---|---|---|---|---|
| | Norm Fluc/Rluc | SD | Norm Fluc/Rluc | SD |
| Vector | 1.0000 | 0.1017 | 1.0000 | 0.07772 |
| PML-ER | 0.8475 | 0.0784 | 0.8101 | 0.1826 |
| Vector 4OHT | 1.0120 | 0.1778 | 1.1320 | 0.2045 |
| PML-ER 4OHT | 0.8001 | 0.1193 | 0.8046 | 0.09078 |
| Vector Gen | 1.4700 | 0.1510 | 1.5350 | 0.4673 |
| PML-ER Gen | 1.3340 | 0.0510 | 1.2730 | 0.01816 |
| Vector 4OHT Gen | 1.5010 | 0.0460 | 1.4530 | 0.04499 |
| PML-ER 4OHT Gen | 0.8767 | 0.1930 | 0.9071 | 0.1145 |

*Relative Fluc fold change*

**UGA Fig 3B**

| Condition | N1 | | N2 | |
|---|---|---|---|---|
| | Norm Fluc/Rluc | SD | Norm Fluc/Rluc | SD |
| Vector D12 | 1.0532 | 0.2135 | 1.0000 | 0.1817 |
| H-Ras V12 D12 | 0.6578 | 0.1054 | 0.7406 | 0.1223 |
| Vector D35 | 1.0000 | 0.0579 | 1.0000 | 0.0579 |
| H-Ras V12 D35 | 0.9209 | 0.0639 | 1.6336 | 0.1719 |

**AQP4 Fig 3B**

| Condition | N1 | | N2 | | N3 | |
|---|---|---|---|---|---|---|
| | Norm Fluc/Rluc | SD | Norm Fluc/Rluc | SD | Norm Fluc/Rluc | SD |
| Vector D12 | 1.0000 | 0.0864 | 1.0000 | 0.0479 | 1.0000 | 0.2210 |
| H-Ras V12 D12 | 0.3274 | 0.1348 | 0.5905 | 0.0455 | 0.4175 | 0.1630 |
| Vector D35 | 1.0000 | 0.0537 | 1.0000 | 0.1802 | 1.0000 | 0.1230 |
| H-Ras V12 D35 | 1.5119 | 0.3644 | 1.8362 | 0.2394 | 1.2496 | 0.2135 |

**VASP Fig 3B**

| Condition | N1 | | N2 | | N3 | |
|---|---|---|---|---|---|---|
| | Norm Fluc/Rluc | SD | Norm Fluc/Rluc | SD | Norm Fluc/Rluc | SD |
| Vector D12 | 1.0000 | 0.0120 | 1.1411 | 0.3336 | 1.0000 | 0.3467 |
| H-Ras V12 D12 | 0.7189 | 0.0187 | 1.0000 | 0.2572 | 0.5411 | 0.0613 |
| Vector D35 | 1.0000 | 0.2016 | 1.0000 | 0.3699 | 1.0000 | 0.2288 |
| H-Ras V12 D35 | 2.9382 | 0.3314 | 2.2841 | 0.2431 | 2.9616 | 0.8335 |

**HPN Fig 3B**

| Condition | N1 | | N2 | | N3 | |
|---|---|---|---|---|---|---|
| | Norm Fluc/Rluc | SD | Norm Fluc/Rluc | SD | Norm Fluc/Rluc | SD |
| Vector D12 | 1.0000 | 0.0505 | 1.0000 | 0.1064 | 1.0000 | 0.1580 |
| H-Ras V12 D12 | 0.5396 | 0.1724 | 0.4845 | 0.6852 | 0.8584 | 0.1746 |
| Vector D35 | 1.0000 | 0.0299 | 1.2092 | 0.4646 | 1.0000 | 0.3842 |
| H-Ras V12 D35 | 3.3707 | 0.3539 | 1.6540 | 0.3531 | 1.6540 | 0.3531 |

**UGA Fig 4A**

| Condition | N1 | | N2 | | N3 | |
|---|---|---|---|---|---|---|
| | Norm Fluc/Rluc | SD | Norm Fluc/Rluc | SD | Norm Fluc/Rluc | SD |
| Vector | 1.0000 | 0.1751 | 1.0000 | 0.0968 | 1.0000 | 0.2026 |
| H-Ras V12 | 0.3895 | 0.2037 | 0.6659 | 0.1597 | 0.1657 | 0.0743 |
| H-Ras V12/E6 | 0.4181 | 0.0877 | 0.8446 | 0.1520 | 0.8052 | 0.1544 |
| H-Ras V12/E7 | 2.3978 | 0.4075 | 3.9306 | 0.4586 | 3.6125 | 0.1093 |
| H-Ras V12/E6/E7 | 5.8209 | 1.1219 | 2.0074 | 0.0243 | 1.6591 | 0.3204 |

**AQP4Fig 4B**

| Condition | N1 | | N2 | | N3 | |
|---|---|---|---|---|---|---|
| | Norm Fluc/Rluc | SD | Norm Fluc/Rluc | SD | Norm Fluc/Rluc | SD |
| Vector | 1 | 0.1443 | 1.045 | 0.1559 | 1 | 0.1168 |
| H-Ras V12 | 0.2937 | 0.076 | 0.8011 | 0.07103 | 0.6803 | 0.1386 |
| H-Ras V12/E6 | 0.3333 | 0.07278 | 0.9688 | 0.0749 | 0.6761 | 0.1346 |
| H-Ras V12/E7 | 0.5688 | 0.09934 | 1.133 | 0.13 | 1.147 | 0.05639 |
| H-Ras V12/E6/E7 | 0.9557 | 0.3249 | 1.374 | 0.04987 | 1.375 | 0.2305 |

**VASP Fig 4C**

| Condition | N1 | | N2 | | N3 | |
|---|---|---|---|---|---|---|
| | Norm Fluc/Rluc | SD | Norm Fluc/Rluc | SD | Norm Fluc/Rluc | SD |
| Vector | 1 | 0.3191 | 1 | 0.2039 | 1 | 0.1326 |
| H-Ras V12 | 0.4874 | 0.02815 | 0.4709 | 0.0702 | 0.7801 | 0.07476 |
| H-Ras V12/E6 | 0.6765 | 0.01425 | 0.4316 | 0.01853 | 0.9158 | 0.05154 |
| H-Ras V12/E7 | 1.844 | 0.3842 | 0.8383 | 0.1091 | 1.342 | 0.1684 |
| H-Ras V12/E6/E7 | 1.536 | 0.11 | 1.517 | 0.0321 | 1.507 | 0.242 |

**AQP4 Fig 4D**

| Condition | N1 | | N2 | | N3 | |
|---|---|---|---|---|---|---|
| | Norm Fluc/Rluc | SD | Norm Fluc/Rluc | SD | Norm Fluc/Rluc | SD |
| Vector | 1 | 0.1713 | 1 | 0.1036 | 1.045 | 0.1559 |
| H-Ras V12 | 0.3133 | 0.06554 | 0.5884 | 0.05405 | 0.8011 | 0.07103 |
| H-Ras V12 /E7 | 0.5997 | 0.1068 | 0.8384 | 0.0461 | 1.133 | 0.13 |
| H-Ras V12/ E7Δ21-24 | 0.2692 | 0.0311 | 0.3682 | 0.0506 | 0.8253 | 0.08496 |

**AQP4 Fig 4E**

| Condition | N1 | | N2 | | N3 | |
|---|---|---|---|---|---|---|
| | Norm Fluc/Rluc | SD | Norm Fluc/Rluc | SD | Norm Fluc/Rluc | SD |
| Vector | 1 | 0.06915 | 1 | 0.2102 | 1 | 0.0437 |
| H-Ras V12 | 0.5729 | 0.07417 | 0.7011 | 0.05802 | 0.66 | 0.01713 |
| Vector /CDK4 | 0.9616 | 0.2299 | 1.127 | 0.3828 | 1.032 | 0.139 |
| H-Ras V12/CDK4 | 0.9731 | 0.08102 | 1.397 | 0.1968 | 0.6475 | 0.1597 |

*Relative Fluc fold change*

**UGA Fig 5A**

| Condition | N1 | | N2 | | N3 | |
|---|---|---|---|---|---|---|
| | Norm Fluc/Rluc | SD | Norm Fluc/Rluc | SD | Norm Fluc/Rluc | SD |
| Vector | 1 | 0.08361 | 1 | 0.3002 | 1.076 | 0.2082 |
| Vector Palbo | 1.372 | 0.1402 | 0.8938 | 0.1501 | 0.9886 | 0.1107 |
| Vector Gen | 20.4 | 2.396 | 10.59 | 1.821 | 7.828 | 1.838 |
| Vector Palbo + Gen | 11.23 | 1.73 | 1.708 | 0.1734 | 5.699 | 1.172 |

**AQP4 Fig 5B**

| Condition | N1 | | N2 | | N3 | |
|---|---|---|---|---|---|---|
| | Norm Fluc/Rluc | SD | Norm Fluc/Rluc | SD | Norm Fluc/Rluc | SD |
| Vector | 1.0000 | 0.1549 | 1.0000 | 0.1802 | 1.0450 | 0.1230 |
| Vector Palbo | 0.7022 | 0.0392 | 0.9950 | 0.2310 | 0.7346 | 0.0666 |
| Vector Gen | 1.5200 | 0.2612 | 4.8140 | 0.5472 | 3.9550 | 0.5800 |
| Vector Palbo + Gen | 1.3210 | 0.1188 | 5.2490 | 0.7914 | 1.8240 | 0.3738 |

**AQP4 Fig 5C**

| Condition | N1 | | N2 | | N3 | |
|---|---|---|---|---|---|---|
| | Norm Fluc/Rluc | SD | Norm Fluc/Rluc | SD | Norm Fluc/Rluc | SD |
| Vector D35 | 1 | 0.1425 | 1 | 0.123 | 1 | 0.2231 |
| H-Ras V12 D35 | 1.466 | 0.1292 | 1.25 | 0.2135 | 1.178 | 0.147 |
| Vector D35 Palbo | 0.573 | 0.2292 | 0.7346 | 0.06662 | 0.2108 | 0.02081 |
| H-Ras V12 D35 Palbo | 0.5395 | 0.08137 | 0.7351 | 0.1591 | 0.3108 | 0.05771 |

**AQP4 Fig6B**

| Condition | N1 | | N2 | | N3 | |
|---|---|---|---|---|---|---|
| | Norm Fluc/Rluc | SD | Norm Fluc/Rluc | SD | Norm Fluc/Rluc | SD |
| Vector D7 | 1.06067427 | 0.108277 | 1.08364861 | 0.129256 | 1 | 0.208843 |
| RPS14 D7 | 0.94751973 | 0.105604 | 0.87998426 | 0.087291 | 0.81639045 | 0.103977 |
| Vector D12 | 1.02080033 | 0.180966 | 1.07130853 | 0.064513 | 0.92031235 | 0.076799 |
| RPS14 D12 | 0.84567442 | 0.137609 | 0.83264707 | 0.130553 | 0.81490701 | 0.151407 |
| Vector D14 | 0.98017514 | 0.194747 | 1.10972634 | 0.15523 | 1.01020603 | 0.017798 |
| RPS14 D14 | 0.71437695 | 0.095787 | 0.6444314 | 0.228738 | 0.65394511 | 0.224815 |

**AQP4 Fig6C**

| Condition | N1 | | N2 | | N3 | |
|---|---|---|---|---|---|---|
| | Norm Fluc/Rluc | SD | Norm Fluc/Rluc | SD | Norm Fluc/Rluc | SD |
| Vector D7 | 1.06067427 | 0.108277 | 1.08364861 | 0.129256 | 1 | 0.208843 |
| RPL22 D7 | 0.64894303 | 0.054812 | 0.70225135 | 0.205294 | 0.72699519 | 0.094085 |
| Vector D12 | 1.02080033 | 0.180966 | 1.07130853 | 0.064513 | 0.92031235 | 0.076799 |
| RPL22 D12 | 0.83588052 | 0.227117 | 0.87195397 | 0.078338 | 0.8806377 | 0.096694 |
| Vector D14 | 0.98017514 | 0.194747 | 1.10972634 | 0.15523 | 1.01020603 | 0.017798 |
| RPL22 D14 | 0.92579054 | 0.042865 | 0.79671393 | 0.032327 | 0.61248296 | 0.12964 |

**UGA Fig S1D**

| Condition | N1 | | N2 | |
|---|---|---|---|---|
| | Norm Fluc/Rluc | SD | Norm Fluc/Rluc | SD |
| Vector D5 | 1.0000 | 0.2828 | 1.0820 | 0.1161 |
| Vector D12 | 0.8620 | 0.3315 | 0.9551 | 0.1077 |
| Vector D20 | 1.0000 | 0.1937 | 0.8995 | 0.2043 |

**AQP4 FigS1D**

| Condition | N1 | | N2 | |
|---|---|---|---|---|
| | Norm Fluc/Rluc | SD | Norm Fluc/Rluc | SD |
| Vector D5 | 0.9991 | 0.2331 | 0.931 | 0.1098 |
| Vector D12 | 0.7887 | 0.1582 | 1.004 | 0.2469 |
| Vector D20 | 0.7829 | 0.1559 | 1.066 | 0.2165 |

**AQP4 FigS1E**

| Condition | N1 | | N2 | |
|---|---|---|---|---|
| | Norm Fluc/Rluc | SD | Norm Fluc/Rluc | SD |
| Vector/Starv | 1 | 0.06281 | 1 | 0.04788 |
| H Ras V12/Starv | 0.4751 | 0.05073 | 0.5905 | 0.04545 |
| Vector /FBS | 0.928 | 0.2528 | 0.8961 | 0.213 |
| H Ras V12/FBS | 0.5003 | 0.1221 | 0.669 | 0.01713 |

**Nat. Rth candidates Ctrl F**

| Condition | N1 | | N2 | | N3 | |
|---|---|---|---|---|---|---|
| | Norm Fluc/Rluc | SD | Norm Fluc/Rluc | SD | Norm Fluc/Rluc | SD |
| HPN-Vector | 1 | 0.075251 | 0.9999999 | 0.050515 | 1 | 0.253091 |
| HPN-H-Ras V12 | 0.3838983 | 0.384538 | 0.7287598 | 0.218933 | 0.5758567 | 0.060332 |
| VASP-Vector | 0.9999998 | 0.008484 | 0.9999998 | 0.292373 | 1 | 0.008485 |
| VASP-H-Ras V12 | 0.7492877 | 0.05589 | 0.8763331 | 0.225433 | 0.7188824 | 0.018714 |
| ASPH-Vector | 0 | 0 | 0 | 0 | 0 | 0 |
| ASPH-H-Ras V12 | 0 | 0 | 0 | 0 | 0 | 0 |
| FBL-Vector | 1 | 0.178778 | 0.9999999 | 0.197851 | 1 | 0.144868 |
| FBL-H-Ras V12 | 0.6416453 | 0.164848 | 0.8305688 | 0.006193 | 0.9574316 | 0.076049 |

**Nat. Rth candidates Gen FigS2**

| Condition | N1 | | N2 | |
|---|---|---|---|---|
| | Norm Fluc/Rluc | SD | Norm Fluc/Rluc | SD |
| HPN-Vector | 8.0240 | 0.9762 | 3.6987 | 0.4123 |
| HPN-H-Ras V12 | 7.9446 | 2.6363 | 2.4933 | 0.1470 |
| VASP-Vector | 19.0749 | 1.8238 | 4.4482 | 0.3532 |
| VASP-H-Ras V12 | 8.2412 | 1.1154 | 3.1289 | 1.0182 |
| ASPH-Vector | 3.6987 | 0.4123 | 1.0000 | 0.4716 |
| ASPH-H-Ras V12 | 2.4933 | 0.1470 | 0.1653 | 0.0323 |
| FBL-Vector | 3.1437 | 0.4392 | 1.8573 | 0.3116 |
| FBL-H-Ras V12 | 1.7089 | 0.1023 | 1.0265 | 0.2637 |

**AQP4 FigS3A**

| Condition | N1 | | N2 | | N3 | |
|---|---|---|---|---|---|---|
| | Norm Fluc/Rluc | SD | Norm Fluc/Rluc | SD | Norm Fluc/Rluc | SD |
| Vector | 1 | 0.1713 | 1.045 | 0.1559 | 1 | 0.1168 |
| Vector /E7 | 1.204 | 0.2172 | 1.121 | 0.2537 | 1.147 | 0.05639 |
| Vector/ E7Δ21-24 | 0.8017 | 0.1072 | 1.321 | 0.0621 | 0.8594 | 0.01346 |

*Fluc/Rluc*

**CGA Fig S1C**

| Condition | N1 | | N2 | |
|---|---|---|---|---|
| | Fluc/Rluc | SD | Fluc/Rluc | SD |
| Vector D5 | 0.1513 | 0.01336 | 0.1522 | 0.0328 |
| H-Ras V12-D5 | 0.1915 | 0.03523 | 0.1917 | 0.02881 |
| Vector D12 | 0.177 | 0.02829 | 0.2036 | 0.05782 |
| H-Ras V12-D12 | 0.1877 | 0.05967 | 0.1653 | 0.01256 |
| Vector D20 | 0.1976 | 0.01948 | 0.1937 | 0.03903 |
| H-Ras V12-D20 | 0.2119 | 0.02777 | 0.2063 | 0.04968 |

**CGA Fig S2A**

| Condition | N1 | | N2 | | N3 | |
|---|---|---|---|---|---|---|
| | Fluc/Rluc | SD | Fluc/Rluc | SD | Fluc/Rluc | SD |
| Vector | 0.1586 | 0.03595 | 0.1653 | 0.0114 | 0.1709 | 0.03088 |
| H Ras V12 | 0.1553 | 0.007879 | 0.1812 | 0.06017 | 0.1478 | 0.02945 |
| Vector Gen | 0.1961 | 0.05043 | 0.2048 | 0.04365 | 0.1997 | 0.03816 |
| H Ras V12 Gen | 0.229 | 0.009903 | 0.2198 | 0.006688 | 0.1437 | 0.005979 |

**UGA Fig S2B**

| Condition | N1 | | N2 | | N3 | |
|---|---|---|---|---|---|---|
| | Fluc/Rluc | SD | Fluc/Rluc | SD | Fluc/Rluc | SD |
| Vector | 1.000 | 0.066 | 1.000 | 0.154 | 1.000 | 0.255 |
| H Ras V12 | 0.625 | 0.162 | 0.748 | 0.063 | 0.620 | 0.034 |
| Vector CDX5-1 | 0.973 | 0.099 | 1.045 | 0.094 | | |
| H Ras V12 CDX5-1 | 0.837 | 0.178 | 0.860 | 0.020 | | |
| Vector Gen | 7.038 | 0.741 | 5.910 | 0.425 | 5.430 | 1.054 |
| H Ras V12 Gen | 3.702 | 0.460 | 3.875 | 0.442 | 3.067 | 1.222 |
| Vector CDX5-1 + Gen | 16.928 | 3.800 | 14.750 | 1.686 | 14.111 | 0.822 |
| H Ras V12 CDX5-1 + Gen | 4.896 | 0.400 | 6.764 | 0.990 | 5.191 | 0.769 |

**Supplementary table 5. Real time PCR primers**

| Gene Name | 5' forward primer (5'-3') | 3' reverse primer (5'-3') |
|---|---|---|
| BRCA1 | GGTTGTTGATGTGGAGGGGCAA | CCAGATTCCAGGTAAGGGGTTC |
| CDKN1A/p21 | ACCCTTGTGCCTCGCTCAGG | GCGTTTGGAGTGGTAGAAATCTGT |
| CDK4 | CAGTGCAGTCGGTGGTACCTGA | GGCAGAGATTCGCTTGTGTGG |
| CENPA | AATGGATTCTGCGATGCTGTCTGG | TTTTCAGGCCTTTGGAACGGTGTT |
| FANCD2 | CCTTAGTAGCCGACTGAAACAGG | TGCAAGTAATGGACGCTCTG |
| GADD45A | TTGCAATATGACTTTGGAGGAA | CATCCCCCACCTTATCCAT |
| HMBS | GGCAATGCGGCTGCAA | GGGTACCCACGCGAATCAC |
| H-RAS | TTGCCATCAACAACACCAAGTC | ATCCGAGTCCTTCACCCGTTTG |
| KI67 | AGAAGACAGTACCGCAGATGA | CGGCTCACTAATTTAACGCTGG |
| MCM6 | ATCCCTCTTGCCAAGGATTT | GAAAAGTTCCGCTCACAAGC |
| ORC1L | CCCGGCTCAAGCATCYAAAGG | TCGTGCATCTCCAGACAGTCT |
| TBP | GCTGGCCCATAGTGATCTTTGC | CTTCACACGCCAAGAAACAGTGA |
